# Deep experimental profiling of microRNA diversity, deployment, and evolution across the *Drosophila* genus

**DOI:** 10.1101/125997

**Authors:** Jaaved Mohammed, Alex S. Flynt, Alexandra M. Panzarino, Md Mosharrof Hussein Mondal, Adam Siepel, Eric C. Lai

**Affiliations:** Cornell University, Department of Biological Statistics and Computational Biology, Ithaca NY 14853, USA; Tri-Institutional Training Program in Computational Biology and Medicine, New York NY 10021, USA; Sloan-Kettering Institute, Department of Developmental Biology, 1275 York Ave, Box 252, New York NY 10065, USA; University of Southern Mississippi, Department of Biological Sciences, Hattiesburg MS 39406, USA; Cold Spring Harbor Laboratory, Simons Center for Quantitative Biology, 1 Bungtown Rd, Cold Spring Harbor NY 11724, USA

**Keywords:** microRNA evolution, mirtron, microRNA birth and death, *Drosophila*

## Abstract

Comparative genomic analyses of microRNAs (miRNAs) have yielded myriad insights into their biogenesis and regulatory activity. While miRNAs have been deeply annotated in a small cohort of model organisms, evolutionary assessments of miRNA flux are clouded by the functional uncertainty of orthologs in related species, and insufficient data regarding the extent of species-specific miRNAs. We address this by generating a comparative small RNA (sRNA) catalog of unprecedented breadth and depth across the *Drosophila* genus, extending our extant deep analyses of *D. melanogaster* with sRNA data from multiple tissues of 11 other fly species. Aggregate analysis of several billion sRNA reads permits curation of accurate and holistic compendia of miRNAs across this genus, providing abundant opportunities to identify species- and clade-specific variation in miRNA identity, abundance, and processing. Amongst well-conserved miRNAs, we observe unexpected cases of clade-specific variation in 5′ end precision, occasional antisense loci, and some putatively non-canonical loci. We also employ strict criteria to identify a massive set (649) of novel, evolutionarily-restricted miRNAs. Amongst the bulk collection of species-restricted miRNAs, two notable subpopulations of rapidly-evolving miRNAs are splicing-derived mirtrons and testis-restricted, clustered (TRC) canonical miRNAs. We quantify rates of miRNA birth and death using our annotation and a phylogenetic model for estimating rates of miRNA turnover in the presence of annotation uncertainty. We show striking differences in birth and death rates across miRNA classes defined by biogenesis pathway, genomic clustering, and tissue restriction, and even identify variation heterogeneity amongst *Drosophila* clades. In particular, distinct molecular rationales underlie the distinct evolutionary behavior of different miRNA classes. We broaden observations made from *D. melanogaster* as Drosophilid-wide principles for opposing evolutionary viewpoints for miRNA maintenance. Mirtrons are associated with a high rate of 3′ untemplated addition, a mechanism that impedes their biogenesis, whereas TRC miRNAs appear to evolve under positive selection. Altogether, these data reveal miRNA diversity amongst *Drosophila* species and permit future discoveries in understanding their emergence and evolution.

## Introduction

MicroRNAs (miRNAs) are ∼22 nucleotide (nt) RNAs that play important regulatory roles in diverse eukaryotic species by promoting transcript degradation or by translational repression [1,2]. In the canonical metazoan pathway, primary miRNA (pri-miRNA) transcripts bearing hairpins are first cleaved in the nucleus by the RNase III cleavage enzyme Drosha. Upon export to the cytoplasm, these precursor miRNA (pre-miRNA) hairpins are further processed into miRNA duplexes by Dicer, another RNase III enzyme. One duplex strand, termed the “mature” miRNA strand, is preferentially retained in an Argonaute (AGO) complex and guides it to complementary mRNA targets. Its partner miRNA* strand, or “star” species, is preferentially degraded, although functional capacity of star strands has been documented. Beyond the canonical pathway, diverse non-canonical biogenesis pathways involving RNases that function in other cellular processes have been uncovered [3]. Chief amongst these is the “mirtron” pathway, in which the Drosha cleavage is substituted by the spliceosome to define either or both pre-miRNA hairpin termini [4-6].

In the evolutionary context, comparative and population genomics have been crucial to our understanding of the functional roles of miRNAs. Such efforts helped define features such as ultra-conservation of the miRNA “seed” sequence (positions ∼2-8 of the miRNA strand that mediate target recognition), and the overall higher constraint upon the miRNA and star strands compared to other partitions of the pre-miRNA hairpin [7-9]. With the availability of *Drosophila* population data, primarily from *D. melanogaster*, deeper insights into miRNA evolution on a more recent timescale emerged, such as the accelerated, adaptive evolution of miRNAs within clusters of testes-restricted expression [10-12]. Additionally, advances in high-throughput sequencing and the availability of miRNA prediction software that leverages this data [13,14], there has been a surge in the annotation of miRNAs across taxa, many of which comprise recently-evolved loci. Overall, miRNAs have now been associated with phenotypic diversity, and their expansion has been correlated with organismal complexity, body-plan innovation, and life cycle [1,2,15].

miRNA catalogs are continually augmented across broad phylogenetic branches, but these have mostly been assessed at the level of presence/absence of miRNA loci [15-17]. Much remains to be explored about miRNA evolutionary features across sets of related species, such as patterns and rates of gene emergence, decay, and expansion, and consistency in processing across orthologs. To date, only a few deep comparisons have been conducted. These include analysis of four *Caenorhabditis* species [18,19] (sRNAs from additional nematode species have been cloned [20], but not analyzed for miRNA emergence), up to six Mammalian species [21,22], and three *Drosophila* species [23,24]. A net gain estimate of 12 miRNA genes/million years (Myr) was first estimated in *Drosophila* [23]. However, this estimate was later revised to 0.82-1.6 genes/Myr using a refined, conservative collection of miRNA genes [24], which proved relatively concordant with a subsequent estimate of 0.83 genes/Myr in mammals [21]. Since these studies, the annotation of *Drosophila* and mammalian miRNAs has expanded by many fold [25]. Nevertheless, despite now thousands of miRNAs collected within the miRBase repository for constituent species of these clades, numbers of pan-*Mammalian* (94) [21], or pan-*Drosophilid* (123) miRNAs [8] have not changed much over the past decade. Thus, there are perhaps an order of magnitude more recently-evolved miRNAs than well-conserved loci in some metazoans.

A relevant consideration is that most previous studies of miRNA evolutionary flux considered them as a unitary class. However, our recent studies provide evidence that subclasses of miRNAs exhibit distinct evolutionary parameters [8,11,17]. For example, we noted that *Drosophila* mirtrons evolve more quickly than canonical miRNAs [24]. This observation was substantiated by our recent strict annotation of ∼500 mirtrons in both mouse and human, nearly all of which are specific to rodents or primates, respectively [26]. One can recognize, then, that expression levels of the minority of conserved miRNA loci dwarf those of the collective majority of miRNA loci, which include both canonical and non-canonical loci. We speculate from this that there should be diverse mechanisms that can drive characteristic evolutionary behaviors of various miRNA classes. However, a foundation to study these may require a deep empirical analysis of species-specific miRNAs across a phylogeny.

In this study, we set out to characterize class-specific properties of miRNAs within 12 species of the *Drosophila* genus, which diverged from the common Dipteran ancestor ∼60 Myr. Advantages of the *Drosophila* system include its wealth of phenotypic diversity, straightforward culture of a wide species collection, access to high-quality whole-genome assemblies, and most importantly, the increased power of fine comparative assessment of evolutionary features within sub-groups of closely-related species [27]. Building on a collection of approximately 1.9 billion sRNA sequences from *D. melanogaster* (summarized in [28,29]), we sequenced an additional ∼1.5 billion sRNAs from embryos, heads, male bodies and female bodies of the other 11 species. This comparative dataset permits evolutionary miRNA analysis at an unprecedented scale. In particular, we elaborate myriad features of miRNA annotation and evolution, and show how these differ with respect to miRNA biogenesis types, tissues with an animal, and between different branches of the fruitfly phylogeny.

## Results

### Compendia of sRNA data across 12 *Drosophila* species

We previously annotated *D. melanogaster* miRNAs from ∼1.9 billion small RNA reads, spanning >100 different developmental stages, tissue types, cell lines, and genetic and environmental manipulations [28,29]. While this scale is not currently feasible to achieve across other Drosophilids, we sought parameters of data collection that would permit deep annotations in other species, and thus serve as an empirical foundation for comparative analyses of miRNA evolution.

Our experience with *D. melanogaster* suggested that mixed embryos, adult heads, male bodies and female bodies are efficacious for broad capture of miRNA diversity. To test this, we performed recovery analyses of *D. melanogaster* miRNAs by subsampling data from these four tissue types (see **Methods**). At an aggregate depth of 100M reads, we recovered 94-98% of conserved (128) miRNAs with at least 30 mature miRNA reads and three miRNA* reads from 100 simulation experiments (**Figure 1A**). Of the 135 miRNAs that emerged recently in the melanogaster-group, we recovered 21-27% of miRNAs using these miRNA/star thresholds. Increasing this depth to 200M reads resulted in a maximal recovery gain to 30-34% of newly-evolved *D. melanogaster* miRNAs (**Figure 1B and/or Supplementary Figure S1**). In other words, doubling the sequencing depth allowed recovery of at most only 7% more newly-evolved miRNAs. Thus, we considered 100M reads from the four tissue types as a strong empirical foundation across these species that would permit broad insights into miRNA evolution.

**Figure 1:**
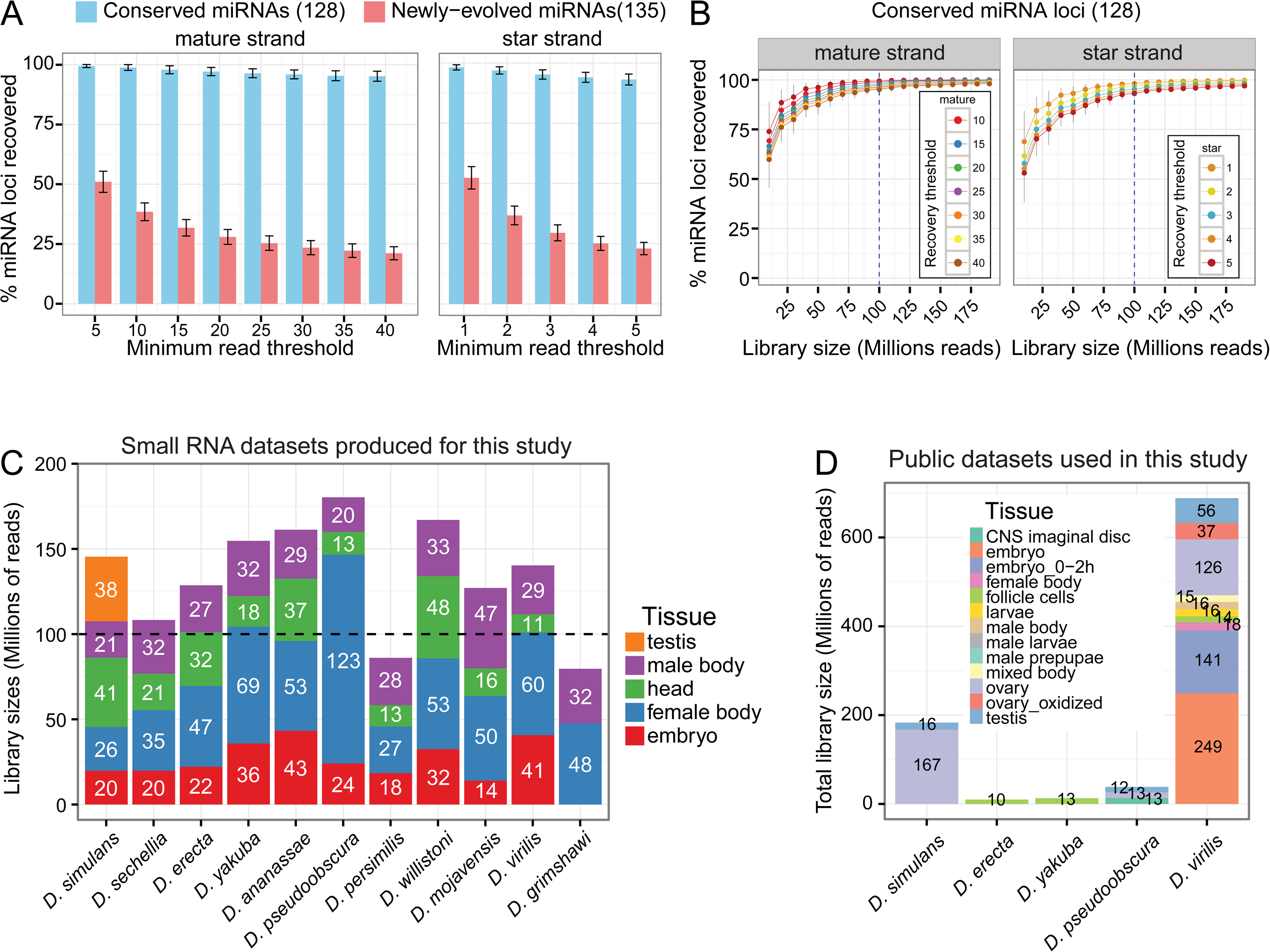
Summary of small RNA sequencing data and analysis of sequencing depth sufficiency. (**A**) The recovery rate of known *D. melanogaster* miRNAs using sets of 100 million (M) total reads sampled randomly from *D. melanogaster* head, mixed embryo, male-body, female-body public libraries. These libraries mimic those we sought to create for the 11 other *Drosophila* genomes. Bars represent the fraction of conserved or newly-evolved *D. melanogaster* miRNAs recovered at various miRNA and miRNA* minimum read thresholds, and error bars represent the standard error of the recovery rate across 100 different samples of 100M reads**. (B)** Saturation curve of miRNA (mature and star) strand recovery at varying minimum read depth cutoffs. Based upon these results, we sequenced 100M reads per species. **(C)** Actual read sequencing depth and library conditions profiled within this study, or **(D)** acquired from public repositories for 11 *Drosophila* species. Datasets for *D. melanogaster* are not shown.

We deeply sequenced 52 small RNA libraries (∼1.5 billion total reads) from these four tissue types across 11 species, exceeding 100M reads for nearly all species (**Figure 1C**). As expected, read lengths of most libraries peaked at 21-22 nts, representing miRNAs, and most body libraries showed an additional 24-28nt peak representing the piRNA population (**Supplementary Table S1, Supplementary Figure S2**). These datasets broadly extend the limited collection of publicly available sRNA data from other fly species, primarily *D. simulans* (∼200M reads) and *D. virilis* (∼700M reads), which we aggregated with our libraries (**Figure 1D**, **Supplementary Table S2**).

### A genus-wide catalog of *Drosophila* miRNA annotations

Several strategies to annotate miRNAs from small RNA data have been developed over the years. These collectively have distinct merits, but no single strategy suffices to discover the full range of confident miRNAs, especially ones with atypical structures and/or non-canonical biogenesis. Therefore, we deployed a multi-pronged framework (see **Methods**) including (1) miRDeep2 (to cast a wide net of candidate hairpins with evidence of cloned small RNA duplexes), (2) an independent set of predicted hairpin structures (especially useful for identifying miRNAs from extended hairpin precursors that are disallowed by miRDeep2), (3) intron annotations (to identify mirtrons and tailed mirtrons, which are systematically overlooked by canonical miRNA finders such as miRDeep2), and (4) whole-genome alignments to identify putative miRNA orthologs across multiple *Drosophila* species (to “rescue” miRNA loci from the candidate pool that have confidently cloned orthologs). Since all initial computational scans include substantial false positives, we subsequently utilized stringent criteria (e.g., abundance, distribution, and patterns of sRNA read alignments indicative of Drosha and Dicer cleavage) and systematic visual inspection of all loci before assigning final annotations to various categories (**Supplementary Figure S3**).

We first queried miRBase (v21) loci for Drosophilid orthologs whose cloned small RNAs had not previously been explicitly identified. This exercise served as an initial check on the overall quality of the datasets, since a basic inference from genomic conservation of a miRNA locus is that it is likely processed into mature small RNAs. Our data support the first cloning evidence for 592 unannotated orthologs of conserved Drosophilid miRNA loci (**Figure 2A**). 512 of these loci were cloned at thresholds of at least 30 miR reads and 3 miR* reads, which would have supported high-confidence de novo annotation. The remainder were cloned at lower thresholds, and would initially have been segregated as “candidates”, but could be recovered based on their orthology to loci well-cloned in other species (80 candidate-rescued, and 42 candidate miRNAs). This supported a rationale to “rescue” certain candidate miRNA loci that fall below high stringency thresholds, but that could be reasonably considered as genuine based on high sequence orthology to a confident miRNA annotation in one or more other species. On the other hand, our deep datasets supported our decision to demote 47 annotations from Drosophilid miRBase loci. These were primarily *D. pseudoobscura* loci for which our present data indicate previous annotations were based on non-miRNA reads (**Supplementary Figure S4**, Supplementary Table S3). Our reassessment of these miRBase loci emphasizes the rigor of current miRNA scoring criteria.

**Figure 2:**
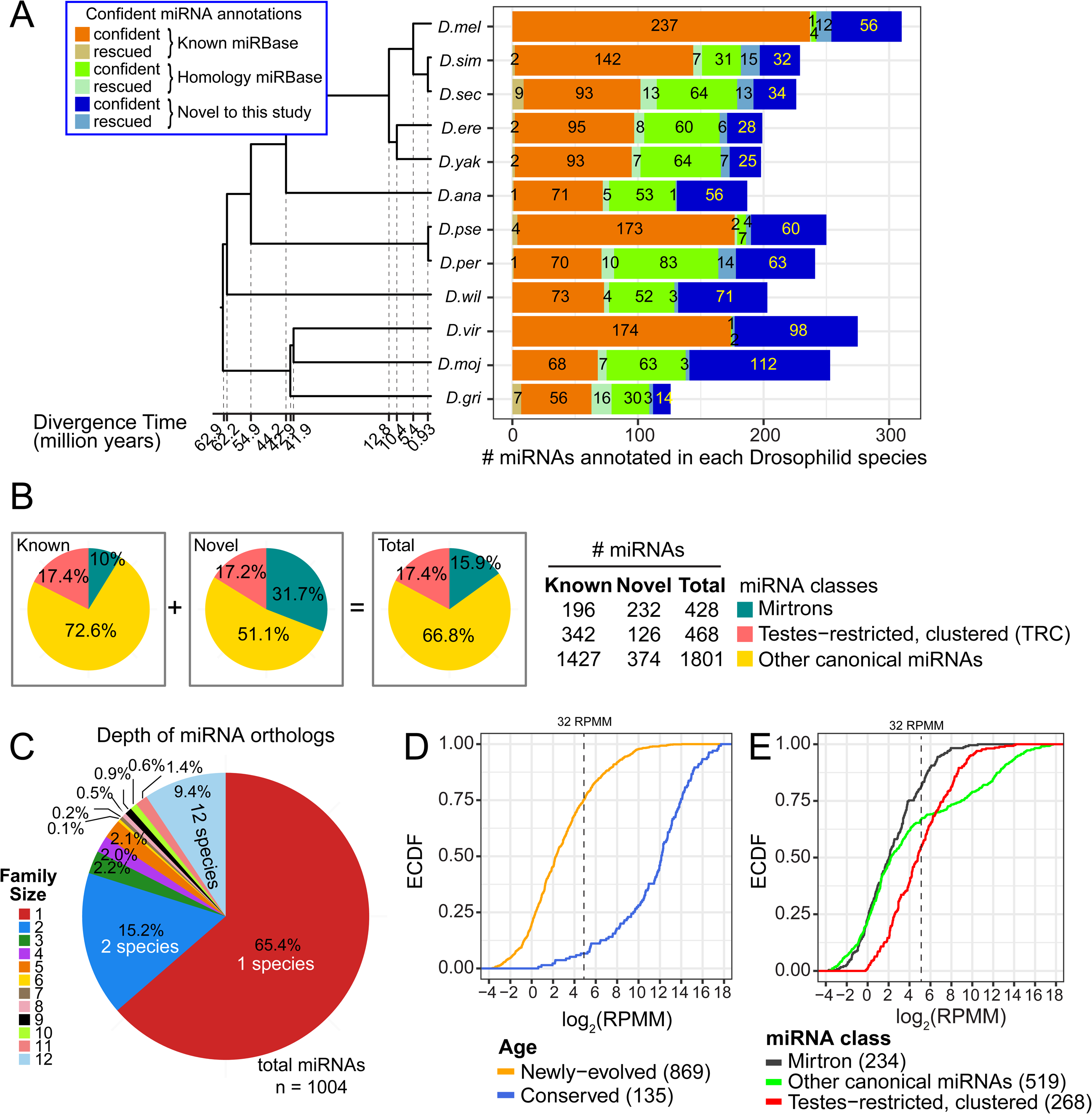
Summary of all known and novel miRNAs recovered within 12 *Drosophila* genomes. **(A)** Counts of known and novel miRNAs recovered or identified, respectively, at our two highest confidence classes – “confident” and “candidate-rescued.” MiRNAs from a third confidence class - “candidate” miRNAs - are shown in *Supp. Fig S5*. **(B)** The proportion of miRNAs recovered within three classes defined by biogenesis pathway, and testes-restricted, clustered status. Pie charts are provided for all novel or known annotations, and for the merged collection. **(C)** The distribution of alignment sizes upon assignment of all miRNAs into 1031 alignments. Paralogous miRNAs were assigned to single species alignment. The majority of miRNAs identified are singletons (species-specific) or doubletons (clade-specific). **(D)** Cumulative distribution function of alignment expression. Alignments are segregated based upon age and miRNA class. Empirical CDFs are plotted using the maximum expression values computed across all constitutive members of each alignment. RPMM = <u>R</u>eads <u>p</u>er <u>M</u>illion mapped <u>m</u>iRNA reads.

For the remainder of our analysis, we grouped the newly-cloned orthologs of miRBase loci along with extant miRBase miRNAs, so as to emphasize the truly novel collection of miRNAs that is unique to our *de novo* annotation effort. In particular, our data support the annotation of 649 novel, confident miRNAs across 12 Drosophilid species (**Figure 2A**). Perhaps not unsurprisingly, many of the highest expressed novel miRNAs are from the *virilis* subclade, which is most distant from *D. melanogaster*. The example of *dvi_264* was cloned at >100,000 reads (**Figure 3A**). At such depth recovery, we observe not only precision of 5p and 3p reads, but we also observe cloned loop reads, which provide evidence of a dominant diced product and indicate 3′-trimming of the predominantly cloned 20 nt species (**Figure 3A**). However, perhaps unexpected was that even in species relatively close to *D. melanogaster*, we still recovered scores of novel confident miRNAs, even though we sampled their small RNAs at <1/10 the read depth and tissue/cell diversity assayed in *D. melanogaster*. For example, *der_50* is but one example of a *melanogaster* subclade miRNA expressed only in *D. erecta* and recovered at a depth of >1000 reads, but absent from *D. melanogaster* (**Figure 3A**). Such observations provide indications of evolutionary flux that we explore later in this study.

**Figure 3:**
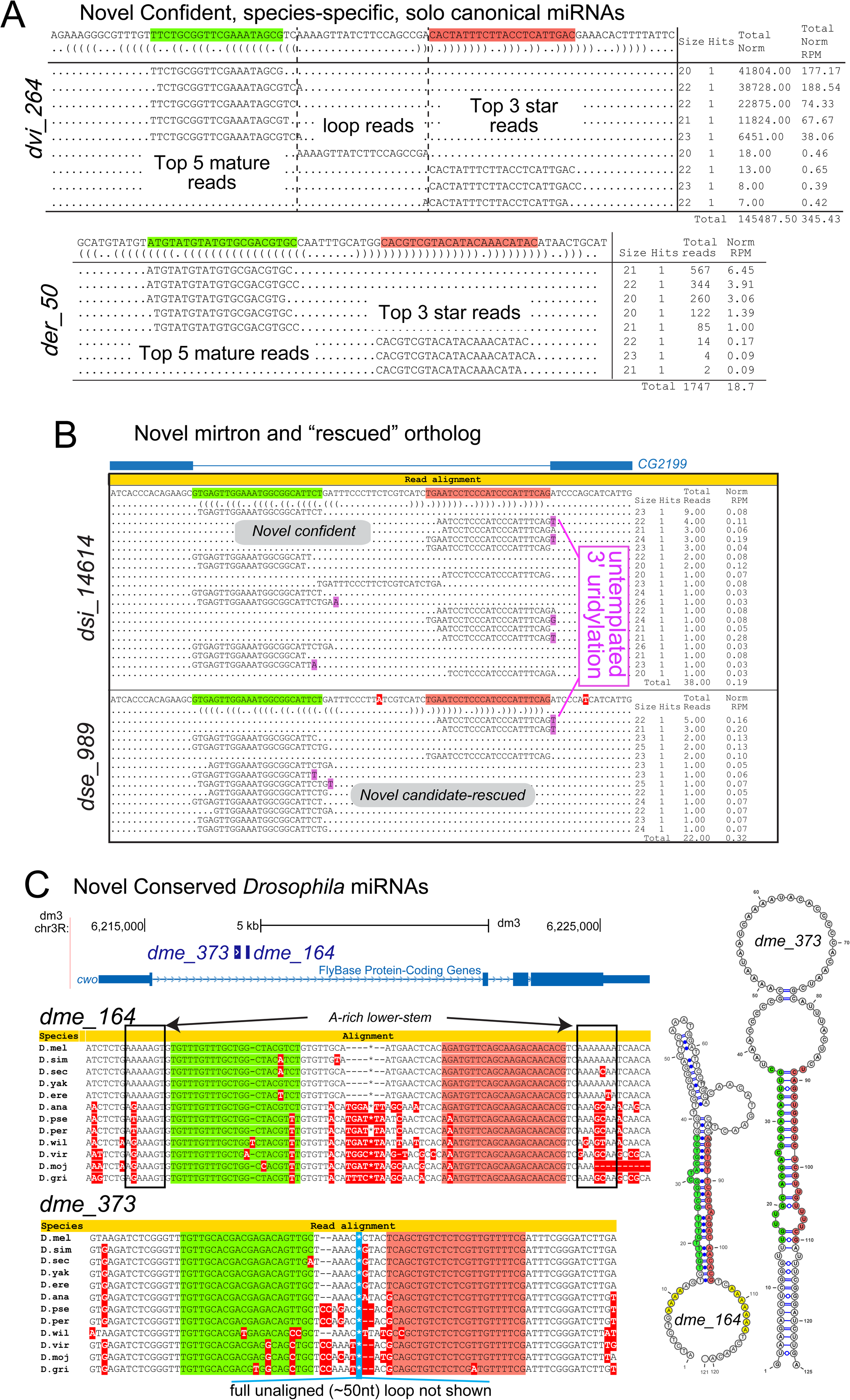
Examples of novel miRNAs identified in this study. **(A)** *dvi_264* and der_50 are examples of confident, canonical miRNAs. Small RNAs were cloned from both arms of their pre-miRNA, and an alignment of these sequences reveal patterns of precise 5’ end cleavage and 1- 2nt 3’ blunt end overhang in the hairpin structure. These are all signatures of Drosha- and Dicer-mediated cleavage. **(B)** An example of orthologous confident (*dsi_14614*) and a candidate-32-rescued (*dse_989*) mirtrons within the two sister species *D. simulans* and *D. sechellia*. Total cloned reads for *dse_939* fell below our threshold for “confident” classification and was effectively placed into the “candidate” confidence group. We “rescued” this locus due to its synteny with *dsi_14614*, a confident mirtron. **(C)** Two novel, conserved, canonical miRNAs identified within the host gene *cwo. dme_164* sequence alignment indicates higher conservation of the miR and miR* arms in comparison to the loop and flanking basal stem regions, a classic signature of pre-miRNA sequence conservation. However, it possesses conserved A-rich regions that flank the duplex, which are seemingly incompatible with a lower-stem needed for Drosha processing. *dme_373* exhibits a large terminal loop atypical of most canonical miRNAs.

**Figure 3B** illustrates *dsi_14614* as a novel non-canonical miRNA of the splicing-derived “mirtron” class. *D. melanogaster* mirtron-3p reads are associated with a high rate 3′ untemplated uridylation [30], and nearly all *dsi_14614-3p* reads are uridylated (**Figure 3B**). Our comparative sRNA data afforded us the unique capability of recovering candidate miRNAs orthologous to confident ones, which we termed as “candidate-rescued” miRNAs; i.e., that we consider as genuine miRNA products. As an example, *dse_989* was classified as a candidate mirtron originally due to the lack of miR* read recovery (**Figure 3B**), however, its synteny to *dsi_14614* allowed us the ability to elevate the confidence of this mirtron. From an initial collection of 765 “candidate” loci bearing small RNA evidence reminiscent of miRNAs, we re-classified 82 as candidate-rescued miRNAs or mirtrons that were clearly orthologous to confident loci (**Figure 2A**, **Supplementary Figure S5**).

The remaining 683 “candidate” loci have small RNA evidence that are reminiscent of miRNAs, but do not meet minimum criteria and are therefore set aside (**Supplementary Figure S5**). We do not presently consider these as genuine miRNA species, even though the veracity of some of these may emerge from deeper or specialized sequencing, and/or they may provide fodder for eventual evolutionary emergence. Nevertheless, our massive set of small RNA data allows us to expand the collection of 1965 known and unannotated miRBase loci of confident and rescued status with 732 novel miRNAs and mirtrons to arrive at a final collection of 2697 total miRNAs and mirtrons present within the *Drosophila* genus. A master table of these loci and their properties is provided as **Supplementary Table S4**. These annotations can be explored in the supplemental website, which provides extensive information regarding read pileups, secondary structures, aligned sequences in other Drosophilid species and so forth (http://compgen.cshl.edu/mirna/12flies/12flies_alignments.html). This miRNA compendium represents one of the largest in any single genus, and we sought to exploit these empirical data to derive insights into miRNA biogenesis and evolution.

### miRNA classes, alignments, and expression

We segregated our aggregate annotations into loci of distinct biogenesis and genomic clustering classes. We first separated canonical miRNAs from mirtrons, a group of small hairpin introns that mimic pre-miRNAs [4,5,31], and further partitioned canonical miRNAs to segregate <u>**T**</u>estes-restricted, <u>**R**</u>ecently-evolved, <u>**C**</u>lustered miRNAs (TRC miRNAs) [11]. We previously utilized *D. melanogaster-centric* miRNA annotations and *D. melanogaster* population data to provide evidence that miRNAs of these three classes evolve by distinct selective pressures, as discerned from their patterns of precursor and “seed” sequence conservation, copy number, and signatures of positive selection [11]. Our highly expanded pan-Drosophilid miRNA annotations provided opportunities to test some of these notions later in this study.

More than half (374 loci; 51.1%) of the novel miRNAs identified in our study and 72.6% (1427) of known and unannotated miRBase loci were solo canonical miRNAs (**Figure 2B**). On the other hand, TRC miRNAs comprised 31.7% (126) of our novel collection, and mirtrons accounted for 17.2% (232). These findings affirm that specific pools of hairpins, namely non-canonical and testis-restricted loci, contribute disproportionately to the aggregate catalog of miRNA substrates. When all miRNAs were evaluated together, we observed 4.5 times more non-TRC canonical miRNAs (66.8%) than TRC miRNAs (15.9%), an enrichment that reproduced across species (**Supplementary Figure S6**).

Curation of accurate miRNA orthologs was paramount to our comparative analyses. We grouped miRNA orthologs into alignments by building upon previous, manual alignments for *D. melanogaster* miRNAs [8], and assigning new miRNAs into groups via genome-wide homology identification and multi-species whole genome alignments (see **Methods**). Altogether, we grouped 2697 miRNAs into 1004 miRNA alignments. From these alignments, we observed that species-specific miRNAs (comprising the majority of loci newly annotated in this study) were the dominant class (65.4%), and miRNAs with one cloned ortholog the next largest class (15.2%). On the other end, 115 alignments (11.4% of loci) contained 10 or more members, and thus were present at the base of the Drosophilid phylogeny (**Figure 2C**).

We next assessed miRNA expression across orthologous miRNAs to understand the range and variation of miRNA expression among alignments. To compare miRNA expression and to assign an expression value per alignment, we computed the log_2_(reads per million mapped miRNA reads, RPMM) score for all loci and recorded the maximum expression level per miRNA in any individual library. We evaluated this metric, instead of the “average” expression of miRNAs, to account for the tissue-specific deployment of many miRNAs. In comparisons of miRNA age, we observed that 92.3% of conserved miRNA alignments had a maximum expression >32 RPMM [i.e. log2(5)], while 24.1% of newly-evolved miRNA achieved such an expression (**Figure 2D**). At a relaxed cutoff threshold, however, we observed that 80.2% of novel miRNA alignments achieved expression of >1 RPMM in at least one library. On the other hand, in comparisons of miRNA class, we observed that TRC miRNA alignments outperformed other classes at conservative maximum expression cutoff (>32 RPMM) (e.g., 45% for TRC versus 19.7% for mirtron and 34.1% for other canonical miRNA) (**Figure 2E**).

### Novel, deeply-conserved miRNAs

Catalogs of well-conserved miRNAs are considered largely complete, as it is generally believed that the set of “clonable” hairpins with miRNA-like evolutionary signatures were exhausted years ago. However, some conserved miRNAs continue to be found, many of which derive from unusual genomic locations or non-canonical pathways, perhaps explaining why they were overlooked earlier. For example, deeply-conserved, non-canonical, *dme-mir-10404* was recently reported to be processed from the internal spacer regions of highly repetitive rRNA loci [32]. Indeed, we find this miRNA is well-cloned from across the Drosophilid phylogeny (**Supplementary Figure S7**).

Amongst our novel miRNA annotations, a handful of loci appeared to be cloned from a broad range of Drosophilid species (**Supplementary Figure S8**). The behavior of *pasha* 5′ UTR hairpins was instructive. A feedback loop in which Drosha cleaves 5′ UTR foldbacks in *pasha/DGCR8* is conserved from mammals to fruitflies, and we previously validated this *in vivo* regulatory interaction using engineered transgenes [33,34]. It was reported that mammalian *DGCR8* 5′ UTR hairpin products of Drosha cleavage are retained in the nucleus, thereby preventing further maturation by Dicer [33]. Nevertheless, sufficient mature reads of *DGCR8* hairpins are found in mammalian deep sequencing data to justify annotation of *mir-3618* and *mir-1306*. We identified small RNA duplexes from the corresponding species ranges for both the deeply conserved (dme_422) and melanogaster group-restricted (dme_474) *pasha* 5′ UTR hairpins (**Supplementary Figure S8**). Although their resultant small RNAs are not particularly abundant, the coupling with our prior evidence of in vivo cleavage of these hairpins by Drosha provides de facto indication of the depth of our deep small RNA profiling.

Amongst novel conserved miRNAs, a notable pair is *dme_373* and *dme_164*, which are clustered in the first intron of *clockwork orange* (*cwo)*. Both loci are highly conserved, exhibit greater loop divergence relative to the hairpin arms that is diagnostic of evolutionarily constrained miRNAs, and are processed into small RNA duplexes across the Drosophilids (**Figure 3C**). However, both loci exhibit atypical features: *dme_164* harbors a conserved, A-rich lower stem that likely precludes Drosha access, while *dme_373* contains an unusually large (∼50 nt) loop (mean loop size of 209 *D. melanogaster* miRNAs is 22 nt). While corresponding reads were detected throughout the Drosophilid phylogeny, we note that their accumulation was modest and efforts to detect them by Northern blotting were negative (data not shown). Thus, while the deep conservation of these hairpins implies functional utility, it remains to be seen if *cwo* miRNAs are matured via non-canonical mechanisms, or if they serve another regulatory role but happen to be sampled in small RNA sequencing. Additional examples of conserved miRNAs are shown in **Supplementary Figure S8** with read details in **Supplementary Figure S9**.

### Evolutionary shifted processing of some conserved miRNA loci

It is generally assumed that genomic conservation of miRNA loci goes hand-in-hand with conserved processing of mature small RNAs, which in turn are locked into conserved regulatory networks. Since even a 1-nt shift in miRNA 5′ identity can redirect its target network, conserved miRNAs are inferred to maintain precise processing. However, in the absence of systematic small RNA sequencing analysis across a genus, the tenets of this assumption have not been challenged by empirical data.

We investigated the consistency in 5′ end processing for 129 well-conserved Drosophilid miRNAs using our comparative data. As expected, the strong majority of conserved miRNAs exhibit 5’ identities (**Supplementary Figure S10**). Of note, some miRNA loci are documented to generate substantial iso-miR species bearing distinct 5′ ends. In general, we observed concordance in iso-miR abundance across these *Drosophila* genomes. For example, two iso-miRs from the 3’ arm of *pre-mir-79* accumulated at a consistent abundance of approximately 3:1 ratio in all species (**Figure 4A**). Such conservation in iso-miR 5’ end processing supports the notion that multiple species from a single locus are incorporated into evolutionarily constrained regulatory networks. By contrast, the heterogeneous processing of *D. melanogaster mir-193* [29] is not consistent across the Drosophilid phylogeny (**Figure 4B**). In particular, its 5′ end shifts by two nucleotides in the virilis subclade relative to the other species. Thus, this conserved miRNA locus appears to be altering its targeting capacity in different species.

**Figure 4:**
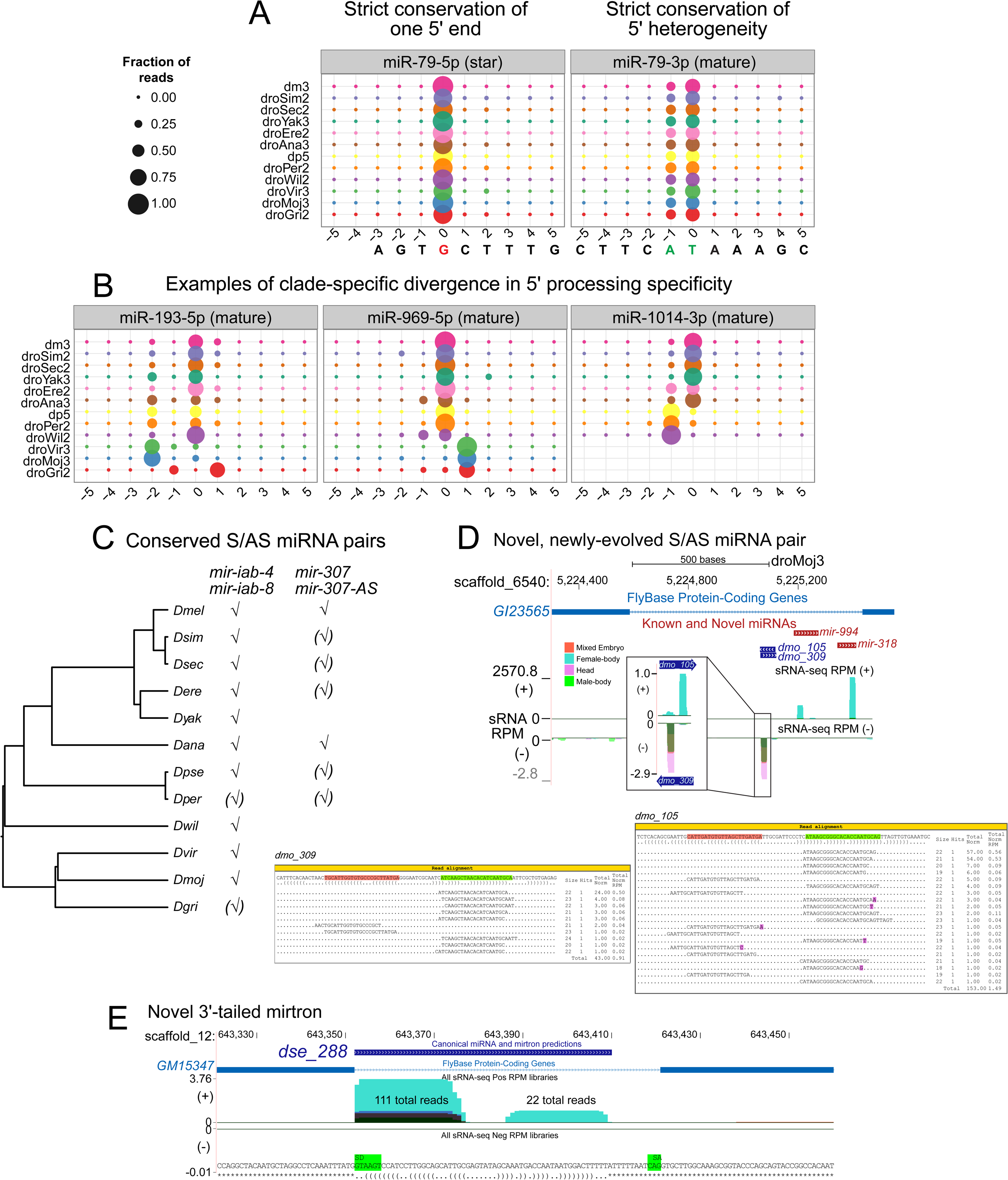
Shifted processing and alternate biogenesis pathways of Drosophila miRNAs. **(A-B)** Consistency in 5’ end processing for conserved miRNAs. **(A)** sRNA read alignment for *mir-79*, represented compactly in these bubble plots, show two miR sequences with unique 5’ ends. These represent two seed-distinct iso-miRs, that are both produced in several *Drosophila* species. Position 0 represents the proportion of reads that begin with the base of the most abundant 5’ arm sequence at either the 5’ strand (miR*) and 3’ strand (miR) for all 12 *Drosophila* genomes. Proportions shown at positions less than or greater than 0 represent proportion of reads with shifted processing. For *mir-79*, two iso-miRs are produced in similar proportions. **(B)** Panels of bubble plots depicting the heterogeneity of 5’ end processing for the miR sequence of other conserved miRNAs and mirtrons. Greater than 4 alternate iso-miR-193 sequences in *D. melanogaster* were noted previously. This heterogeneity is preserved in the genomes of the other Drosophilids, and a conserved, dominant iso-miRs is not apparent. We identified clade-specific iso-miRs for one canonical miRNA (*mir-969*) and one mirtron (*mir-1014*). Specifically, two unique iso-miR-969 sequences are each preferentially abundant in the Sophophora-group and *Drosophila-group* species, respectively, and for *mir-1014*, the melanogaster-group species produces one iso-miR-1014 sequence that is distinct from the dominant iso-miR of other Sophophorans**. (C)** *mir-iab-4/8* and *mir-307/mir-307-as* are the only two reasonably conserved miRNAs with sense and antisense transcription and processing based upon our genus-wide data. **(D)** Dozens of other recently-evolved antisense miRNAs were identified in our study however, such as *dmo_105/dmo_309* that arose adjacent to the conserved *mir-994/318* cluster. **(E)** Example of an atypical splicing derived, 3’ tailed mirtron, identified from our data, *dse_288*. It requires trimming of ∼12 nts from the debranched 3′ splice site to generate the pre-miRNA hairpin.

We identified other cases of clade-specific shifts in processing register for conserved mature strand miRNAs. For example, all *drosophila-group* species consistently processed *mir-959* into a particular species, but a different iso-miR appears substantially in the *Dana/Dpse/Dper/Dwil* ancestor, while Sophophora-group species dominantly accumulate a completely distinct, third iso-miR-959 (**Figure 4B**). We also observe evolutionary shifts for non-canonical loci. This is illustrated by clade-specific shifts in 5′ processing for mirtron-derived miR-1014-3p (**Figure 4B**). Such alterations in targeting capacity of conserved miRNAs were hidden until the availability of deep, evolutionary profiling of small RNAs across the genus, and remind us that miRNA processing cannot be well predicted from genomic sequence alone.

### Antisense miRNAs

Certain miRNA loci are transcribed and processed on both strands [35]. A marquee example in *Drosophila* is *mir-iab-4/mir-iab-8*, for which sense and antisense miRNAs have distinct and genetically overt neural functions [36,37]. Since sense/antisense transcription of this miRNA locus has been observed in beetles [38], one might assume this is a conserved feature throughout the Drosophilids. Indeed, we confidently annotated *mir-iab-4* and *mir-iab-8* in all *Drosophila* species, except for *D. persimilis* and *D. grimshawi* where the lower-expressed *mir-iab-8* locus had reads but had to be recovered through the “candidate-rescue” pipeline (**Figure 4C**).

We observed 18 other confident sense/antisense miRNA pairs, as well as several dozen candidate antisense miRNAs (**Supplementary Table S5**). A few of these involve conserved miRNAs. The most broadly conserved antisense locus was *mir-307-AS*, which was confidently or candidately detected in seven related Drosophilid species (**Figure 4C**). The modest, but clear, cross-species accumulation of *mir-307-AS* might simply reflect low expression, but alternatively it may have spatially or temporally restricted deployment. However, most antisense miRNAs were poorly conserved, and in fact a substantial fraction of them were present in species other than *D. melanogaster* (**Supplementary Figure S11**). For example, *D. mojavensis* generated a novel sense/antisense locus *dmo_105/dmo_309* adjacent to the deeply conserved intronic miRNAs *mir-994/mir-318* (**Figure 4D**). We present several additional example of sense/antisense miRNA pairs in **Supplementary Figure S11**, including loci that originated apparently de novo in individual species. Overall, it appears that only sense/antisense miRNA pairs have originated across all branches of the Drosophilid phylogeny, but few instances have been retained over substantial periods of evolution.

### Tailed mirtrons

Amongst miRNAs of non-canonical biogenesis, mirtrons form the dominant class. In the originally described pathway, splicing directly generates a pre-miRNA mimic that is appropriate for export from the nucleus and then cleavage by Dicer [4,5]. Later, alternative “tailed mirtrons” were described, for which further trimming from either the 5′ or 3′ end must occur to generate the pre-miRNA substrate [31]. In *D. melanogaster*, we identified one 3′-tailed mirtron (*mir-1017*) that is broadly conserved in other flies, along with a handful of other 3′-tailed mirtron candidates [6]. Recently, we appreciated that mammals express a few dozen conventional mirtrons and 3′-tailed mirtrons, but hundreds of hundreds of 5′-tailed mirtrons [26]. Thus, there is differential utilization of tailed mirtron pathways between invertebrates and mammals.

Of the 236 novel mirtrons annotated across the Drosophila genus in this study (**Figure 2B**), nearly all are of conventional subtype. Thus, flies do not seem to have propensity for tailed mirtrons. Moreover, while our compendia of candidates includes some potential 5′-tailed mirtrons, we do not feel confident in the read patterns to classify any of this class in any Drosophila species. Thus, this seems to be a substantial distinction from mammals. Amongst 3′-tailed mirtrons, we recovered *mir-1017* in all species excepting D. grimshawi, which has a slightly lower total library size (*mir-1017* is known to be restricted to the nervous system). We also recovered the other five previous proposed 3′-tailed mirtrons in our forward annotation pipeline [6], all of which were known supported by much higher read depth.

We also recovered 6 novel confident 3′ tailed mirtrons (**Supplementary Table 4**). An example is *D. sechellia dse_288*, which requires removal of a ∼12 nt tail following splicing and debranching (**Figure 4E**). Although it is clearly supported by >110 mature and >20 star reads, this locus is not processed as a tailed mirtron in its closely related sister species in the simulans clade. We provide details of another novel 3′ tailed mirtron from *D. erecta der_70* in **Supplementary Figure 12**. *Der_70* is notable not only as an atypical mirtron, but also for bearing a non-canonical GC splice donor within the corresponding splice junctions of the *CYLD* gene of all five melanogaster-group orthologs; the other Drosophilid species harbor a typical GT splice donor. This is the first recognized example of non-canonical mirtron splicing. Curiously, none of the eleven confident 3′-tailed mirtrons we detected have conserved processing, even in closely related sister species.

### Massive evolutionary flux of testes-restricted miRNA clusters

The second largest class of novel miRNAs we annotated classified as testis-restricted, clustered (TRC) miRNAs, based on their residence in genomic clusters and preferred or exclusive accumulation in male body/testis libraries relative to other tisssue libraries. We identified 126 novel TRC miRNAs, all of which were recently-evolved, which represented 17% of all new miRNA annotations. Collectively, this abundance of novel miRNAs clustered into nine novel genomic regions within the genomes of different Drosophila species (**Supplementary Figures S13-16**).

Strikingly, we find one or more novel TRC clusters specific to each major Drosophilid branch. Beyond the previously described miRNA clusters composed of conserved and recently-emerged testis-expressed miRNAs [11], we discovered many new TRC specific to *D. ananassae* (2 TRC, containing 34 miRNAs) (**Figure 5A**) or *D. willistoni* (1 TRC, containing 19 miRNAs), and orthologous clusters that were only traceable between closely related sister species, such as between *D. virilis* and *D. mojavensis* (2 TRC shared by these species, containing 31-34 miRNAs, with 2 additional virilis-specific clusters containing 18 miRNAs), or between *D. pseudoobscura* and *D. persimilis* (2 TRC shared by these species, containing 47-50 miRNAs) (**Supplementary Figures S13-16**). Some of these clusters dwarf the largest previously known fly miRNA clusters. For example, we expanded the membership of the *D. pseudoobscura dps_3416→dps-mir-2536* TRC to 36 miRNAs, and identified an orthologous 26-member *D. persimilis* cluster (**Figure 5B**). Small RNA expression showed significantly higher expression in male body and testis libraries than other tissues (**Figure 5B, C**). Interestingly, while miRNAs in the 3’ region of this cluster preserve their order between the two species, miRNAs near the 5’ end the cluster evolved rapidly via both local gene duplication and *de novo* miRNA emergence, as evident from precursor and miRNA sequence alignments (**Figure 5B** family assignments and **Supplementary Figure S15**).

**Figure 5:**
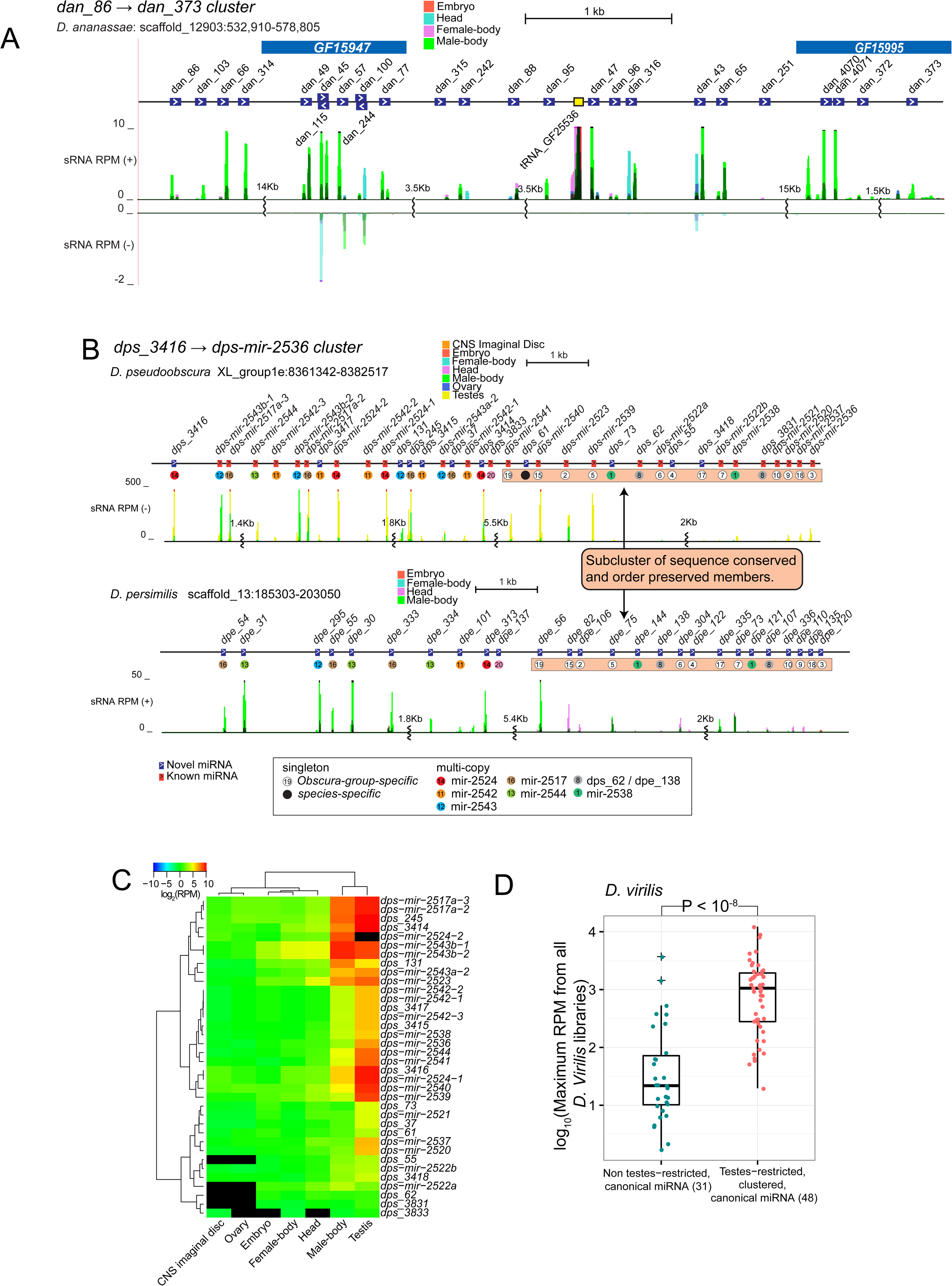
<u>T</u>estes-restricted, <u>R</u>ecently-evolved, <u>C</u>lustered (TRC) canonical miRNAs in *Drosophila*. **(A)** An example of a novel TRC miRNA cluster (*dan_86 → dan_373*) in *D. ananassae*. The majority of miRNAs show high expression in the male-body libraries. **(B)** An example of a TRC miRNA cluster (*dps_3416 → dps-mir-2536*) in the obscura-group species. The *D. pseudoobscura* cluster contains 36 miRNAs while its sister species, *D. persimilis*, contains 26 miRNAs. MiRNAs within the 3’ end region of these orthologous clusters (orange highlight) have preserved their order while miRNAs within the 5’ region show high gene duplication. Colored circles and numbers represent miRNAs of the same family. **(C)** Expression heatmap for all *D. pseudoobscura* copies reveals a predominant testes-restricted profile**. (D)** Comparison of expression difference between TRC and solo canonical miRNAs present in *D. virilis* alone or within the virilis/mojavensis clade alone. TRC miRNAs of the virilis-subgroup show significantly higher expression than their age-matched solo canonical cohorts (*Mann-Whitney Test* p < 10^−8^). All Drosophilid subclades have their own distinct TRC loci, and details of all the novel TRC loci cloned in this study are provided in **Supplementary Figures S13-S17**.

The wealth of male-body sRNA libraries for all 12 genomes and testis data for *D. pseudoobscura* and *D. virilis* permitted the evaluation of the relative expression of TRC miRNAs to that of age-matched solo canonical miRNA cohorts. In *D. virilis*, the species with the greatest number of TRC miRNAs (66 in total including known and novel TRC miRNAs), we observed significantly higher expression for TRC miRNAs than for solo canonical miRNAs (Mann-Whitney U-test p < 10^−8^) (**Figure 5D**). Although we observed a similar shift for substantially increased average expression of TRC miRNAs in *D. pseudoobscura*, this did not quite achieve significance over age-matched non-TRC canonical miRNAs due to a small number of highly-expressed loci in the latter category (**Supplementary Figure S17**).

Altogether, the massive flux of clustered testis miRNA loci across the Drosophilid phylogeny generalizes their distinct evolutionary features that we had established from studies based on a *D. melanogaster-centric* viewpoint. These data provide strong evidence that TRC miRNAs are unlikely to be evolving along a purifying selection route, but instead may be utilized for adaptive regulatory purposes.

### Distinct rates of gain and losses between miRNAs classes

Estimating rates of evolutionary turnover for genomic elements, be they non-coding RNAs, *cis*-regulatory elements, or protein-coding genes, remains an active area of investigation. miRNA turnover rates have been estimated within *Drosophila*, but these efforts have been limited by uncertainty in miRNA annotations and the number of species considered [23,24,39]. Consequently, many newly-evolved loci have been unaccounted for. In light of new evidence that miRNA sequence and structure evolution are influenced by multiple factors, including biogenesis pathways, clustering state, and testes-biased expression [2,8,11,29], we sought to understand whether such factors influence miRNA birth and death rates.

Using our updated *Drosophila* miRNA collection, we set out to characterize and compare rates across our three miRNA classes. To accomplish this, we developed a phylogenetic probabilistic graphical model with the intention of estimating miRNA gene birth and death rates by maximum likelihood (see **Methods**, **Figure 6A**). This method allows us (1) to infer universal, clade-, or branch-specific birth (*λ*) and death (*μ*) rate parameters, (2) to predict node-wise ancestral miRNA presence or absence; and (3) to estimate expected counts of edge-wise gain and loss events. We acknowledge that some small RNAs were sampled more deeply in certain species, especially in *D. melanogaster* (**Figure 1**). However, besides *D. melanogaster*, there is in general not a strict correlation between the depth of sampling and the number of confidently annotated miRNAs per species. This is due in part to the “rescue” approach (**Figure 3**). Therefore, we chose to apply our estimates of miRNA flux using our full collection of annotations, rather than by attempting to make a new set of annotations by subsampling a lower, fixed number of reads from across the species, since this would inevitably decrease annotation confidence.

**Figure 6.**
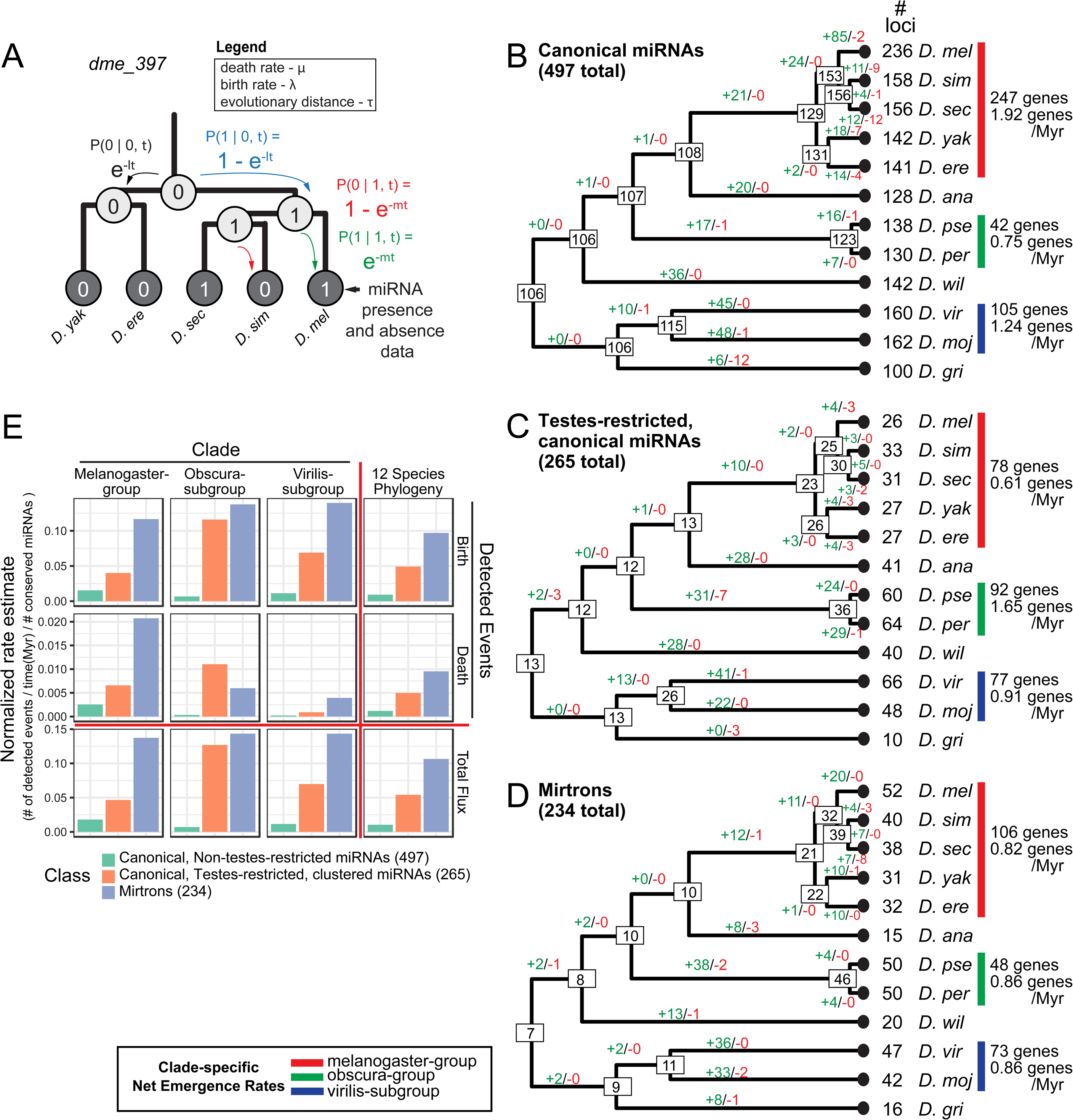
Estimation of miRNA birth and death rates in *Drosophila*. **(A)** A probabilistic, phylogenetic graphical model for estimating rates of gene birth and death. The model takes binary data representing miRNA presence or absence at the leaves of the tree and uses maximum likelihood and numerical optimization methods to estimate model parameter (*μ, λ*) values. Branch lengths (τ) are fixed. Maximum likelihood parameters are then used to reconstruct node-wise miRNA counts and edge-wise birth and death events. **(B-D)** Summary of estimated ancestral miRNA content and edge-wise birth and death events for three classes of miRNAs. MiRNA classes are canonical miRNAs **(B),** Testes-restricted, canonical miRNAs (TRC) **(C),** and mirtrons **(D)**. Estimates of edge wise birth and death events are shown in green and red, respectively. Net emergence rate (i.e. total birth - death events / Myr) are shown in each class for the melanogaster-group, obscura-group, and virilis-subgroup species**. (E)** Net miRNA gain rate for three clades- melanogaster-group, obscura-group, and virilis subgroup – are shown. Note that mirtrons and TRC miRNAs exhibit much higher rates of flux than do canonical non-testis-restricted miRNAs.

We applied our method to the pooled collection of miRNA families from our three classes and estimated model parameters (i.e. *λ, μ*). Using these rate estimates, we then reconstructed node-wise presence and absence of ancestral miRNAs, and subsequently branch-wise miRNA birth and death events, for each miRNA alignment family and across all miRNA classes (summarized in **Figure 6B-D**, see **Supplementary Figure S18** for examples of all possible tree configurations, and **Supplementary Figures S19-S21** for trees per miRNA alignments across three miRNA classes). Consistent with previous studies, the canonical miRNA class contained the largest number of ancient miRNAs, that is, those present at the root of the *Drosophila* phylogeny. Of 236 *D. melanogaster* canonical miRNAs, 106 were clearly present in the Drosophilid ancestor (**Figure 6B**). As mentioned, there are more canonical miRNAs annotated in D. melanogaster than any other fly species owing to its depth of sequencing; otherwise, the majority of canonical miRNAs that are not testis-restricted are conserved (**Figure 6B**).

The tables are turned when examining the fraction of conserved loci in the other categories of miRNAs. The strong majority of testes-restricted canonical miRNAs across the Drosophilid phylogeny, most of which are arranged in genomic clusters, are not conserved. Indeed, only 13 such miRNAs are deeply conserved, and include specific members of the *mir-972→979* cluster and the *mir-959→964* cluster. Otherwise, most fly species harbor dozens of lineage-restricted TRC miRNAs (**Figure 6C**). Similarly, the mirtron class also contains very few conserved loci. This notion was suggested earlier [24], but we now broadly extend this principle using empirical annotation of mirtrons across 12 *Drosophila* species. While the genomes of several well-profiled and deeply-mined, extant species contained >45 mirtrons (e.g., *D. melanogaster* with 52, *D. pseudoobscura* with 50, *D. virilis* with 47 mirtrons), only 7 mirtrons were present at the root of the Drosophilid phylogeny (**Figure 6D**).

Next, we computed rates of miRNA birth and death by first aggregating birth and death events across important clades (melanogaster-group, obscura-subgroup and virilis-subgroup) or the entire phylogeny for each miRNA class, and normalizing them by both the total branch length [in Millions of years (Myr)] and by conserved members of each class. These clade-specific and tree-wide rate estimates permitted additional intra-miRNA-class comparisons of total miRNA flux (i.e. birth plus death) in each of these representative clades or across the entire phylogeny. Interestingly, we saw striking rate variation across the three classes of miRNAs (**Figure 6E**). In general, canonical non-testes-restricted miRNAs exhibited the lowest rates of birth, death, and total miRNA flux in each clade and across the *Drosophila* phylogeny when compared to the two other classes. At the other end of the spectrum, mirtrons exhibited the highest rate estimates. This is attributable not only to the large collection of single-species mirtron annotation in our collection (**Figure 6D**), but also to certain atypical patterns of mirtron presence within extant species that do not group along clade boundaries. For example, *dps_22* is present within both obscura-subgroup species and *D. virilis*, but absent in other Drosophilids. This and other cases are shown in **Supplementary Figure S22**.

Testes-restricted canonical miRNAs exhibited birth, death and total flux rates in between those of canonical non-testes-restricted miRNA and mirtrons, but showed a significantly elevated death rate within the obscura-subgroup when compared to the other miRNA classes. This observation adds to our previous findings of TRC miRNA death within this clade. That is, orthologs of both *D. pseudoobscura* and *D. persimilis* are missing in part or entirety for two pan-Drosophilid TRC miRNA clusters (*dme-mir-959 → 964 and dme-mir-972 → 979*) [11].

Altogether, these findings support the previous hypothesis for the non-homogeneity in mirtron and canonical miRNA rates of evolution [2]. Moreover, we highlight differential behavior along certain lineages; i.e., with accelerated flux of TRC miRNAs within the obscura-group species.

### Multiple mechanisms underlie distinct flux behaviors of different miRNA classes

We explored several molecular strategies that could underlie the distinct evolutionary behaviors of different miRNA classes using these comprehensive novel miRNA annotations.

#### 1. cis-mutations

The impact of nucleotide changes themselves, especially those that are sparse among orthologous pre-miRNAs sequences, are little known on miRNA expression, genesis or decay. We identified compelling sets of miRNA orthologs for experimental tests, including ones exhibiting large variations in apparent biogenesis despite sometimes full genomic identity in the mature miRNA species.

For example, the melanogaster-subgroup-specific locus *mir-4984* is well-aligned across five genomes and exhibits only a few substitutions restricted to the miRNA* region of its pre-miRNA, yet large expression differences between orthologs (**Figure 7A**). Only *D. melanogaster, D. simulans*, and *D. sechellia* orthologs are processed, and no reads were mapped to the *D. yakuba* and *D. erecta* orthologs are not expressed. Even between expressed orthologs, there is a >10-fold reduced normalized RPM expression in *D. simulans* (0.06) as compared to *D. melanogaster* (0.66) and *D. sechellia* (0.82) orthologs that may be driven by a single miRNA* A-to-G substitution. We validated these expression levels deduced from the sRNA-seq data with independent Northern blot assays (**Figure 7B**). That is, we detected both the pre-miRNA and mature products *for dme-mir-4984*, yet failed to detect mature species for both *dya-mir-4984* and *dsi-mir-4984*. Another example is the alignment of *dps_41* and *dpe_2484*, which are specific to the obscura-subgroup. *dps_41* is >100 times higher expressed than *dpe_41* in the sRNA data (**Figure 7C**) and *dpe_41* is undetectable in the corresponding Northern blot (**Figure 7D**).

**Figure 7.**
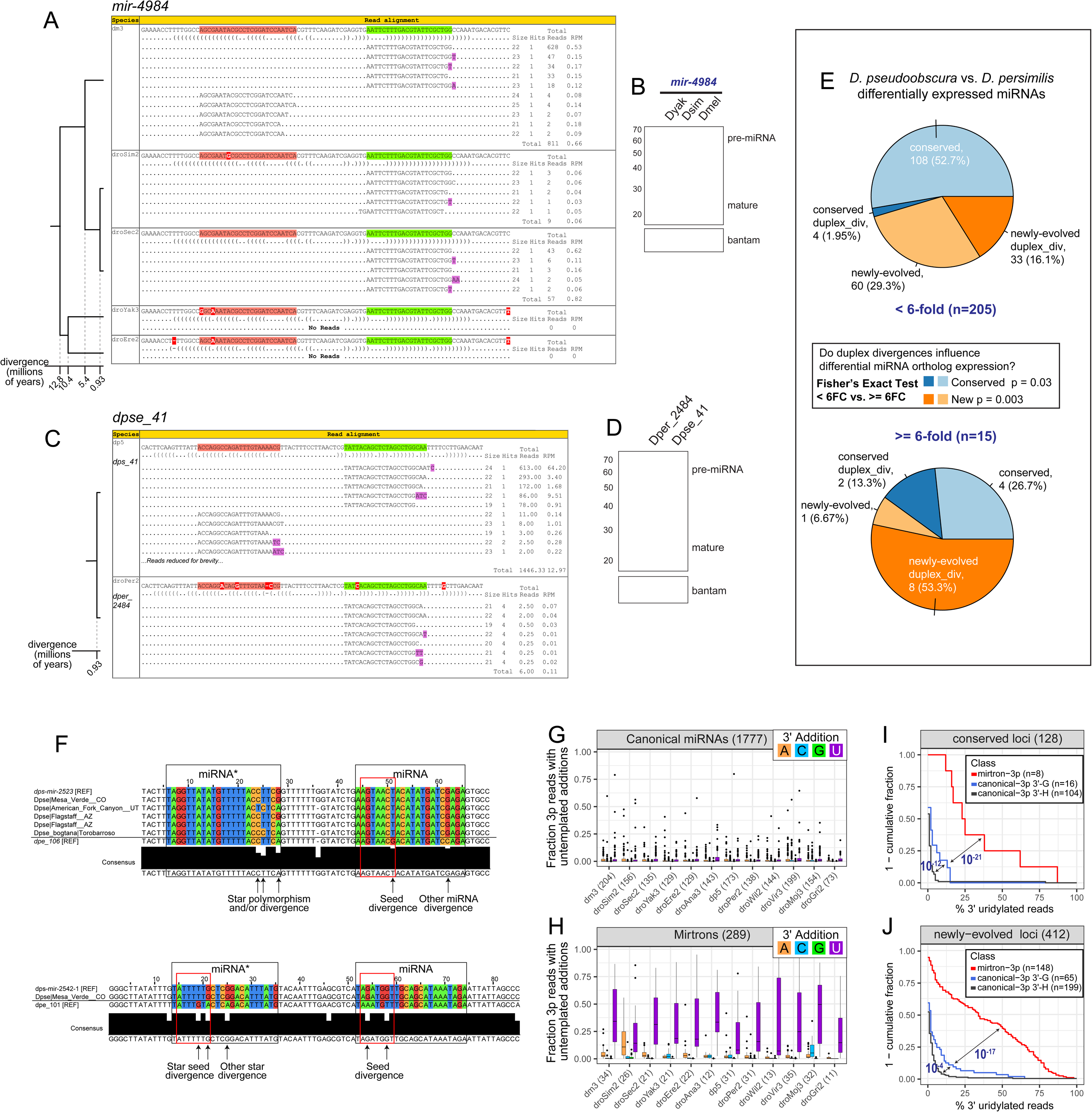
Multiple distinct cis-molecular signatures associated with miRNA flux. **(A-D)** Duplex alterations that affect miRNA processing. **(A)** *mir-4984* hairpin is similar across 5 related melanogaster subgroup species and its mature (green) arm is identical in all these species. However, small RNA sequencing indicates substantial accumulation only in *Dmel*, very modest in *Dsim/Dyak*, and not in *Dyak/Dere*. **(B)** Experimental tests of *UAS-DsRed-mir-4984* expression constructs transfected into S2 cells shows that only the *Dmel* construct was effectively processed into miRNAs. DsRed expression confirms that all constructs were expressed. **(C)** *dps_41/dpe_2484* ortholog pair, with only a few duplex divergences, exhibits divergent expression between very closely related species. **(D)** Experimental tests in S2 cells confirm differential biogenesis of these miRNAs. (**E**) Transcriptome comparison of miRNAs differentially expressed between sister species *Dpse* and *Dper*. In general, significantly more duplex divergent miRNAs are differentially-expressed miRNAs for both newly-evolved and conserved miRNAs. **(F)** Adaptive evolution of seed regions of testis-restricted, clustered (TRC) miRNAs. Shown are examples of 1-to-1 orthologs of TRC miRNAs between *Dpse* and *Dper*, including available *Dpse* population data. Highlighted are examples of seed divergence between expressed TRC miRNA orthologs between these closely related species, indicating adaptive evolutionary behavior**. (G-J)** Impact of terminal uridylation system on evolutionary suppression of mirtrons and behavior of canonical miRNAs**. (G-H)** Compared to canonical miRNAs **(G),** mirtrons **(H)** in every Drosophilid species acquire extraordinarily high rates of terminal untemplated uridylation (purple) on the 3′ ends of their 3p species, compared to any other nucleotide modifications. **(I-J)** 3′ uridylation of canonical miRNAs is sensitive to terminal hairpin nucleotide. In these graphs, miRNA loci are divided by biogenesis type (canonical vs. splicing-derived), by terminal nucleotide (3′-G vs. 3′-A/U/C, i.e. “3′-H”), and by evolutionary age. Analysis of deeply conserved miRNA loci **(I)** and recently-evolved loci **(J)** shows that canonical miRNA hairpins that end in G acquire higher levels of 3′ uridylation than do other canonical miRNA hairpins.

We broadened this analysis using *D. pseudoobscura/persimilis*, which had advantages for being a closely related (0.93 Myr) sister pair for which we had identified numerous novel miRNA annotations that might potentially be subject to expression fluctuation. We labeled miRNA alignments with >= 6-fold log_10_(RPMM) expression difference as differentially-expressed, and identified six conserved and nine newly-evolved miRNAs as such (**Supplementary Figure S23**). The conserved loci included several members of the *mir-309* cluster and seemed to be a sampling artifact given they are expressed as a highly stage-specific operon [40]. Otherwise, there was a high correlation (0.938) between the remaining 205 *D. pseudoobscura* and *D. persimilis* ortholog pairs (i.e. < 6-fold change) (**Supplementary Figure S23**).

We sought to investigate the sequence features between the 15 differentially-expressed, obscura-group ortholog pairs by investigating the patterns of the miR:miR* duplex conservation. In all cases, orthologs shared high sequence and structure similarity apart from a few substitutions. For example, we identified *dps-mir-2567* and *dpe_2484* as differentially-expressed, recently-evolved orthologs (*dps-mir-2567* = 12.97 RPM; *dpe_2484* = 0.11 RPM) (**Supplementary Figure S23**). Between these two species, we observed five duplex substitutions, one of which resides in the seed region of the dominant 5’ arm. In light of these apparent substitutions, we asked if duplex substitutions were more prevalent between differentially-expressed miRNAs than non-differentially expressed ones for both conserved and newly-evolved miRNAs. Indeed, within the obscura-group we saw significantly more differentially expressed miRNAs with duplex substitutions within the newly-evolved group (Fisher’s Exact Test [FET] P < 0.003), and within the conserved group (FET P = 0.03) (**Figure 7E**).

#### 2. Adaptive seed mutations of TRC miRNAs

Functional miRNAs are not expected not to diverge between closely related species, especially within seed regions. However, we previously used *D. melanogaster* population data and melanogaster group species orthologs to provide evidence for adaptive evolution of TRC miRNAs in this clade, including within seeds [11]. There is limited population data in other Drosophilids, but we investigated polymorphisms from whole genome sequences of 11 North American *D. pseudoobscura* strains and 2 *D. pseudoobscura bogotana* sub-species (data available from http://pseudobase.biology.duke.edu/) [41]. To identify unambiguous divergences between species, we focused on miRNAs with clear 1-to-1 orthologs, such as the miRNAs within the 3’ sub-cluster region of the *dps_3416→ dps-mir-2536* cluster.

We identified two TRC miRNAs with seed divergences in this sub-cluster. For example, the mature (3’) arm of *dps-mir-2523* contained a G-to-T substitution at the 8^th^ seed position relative to its *D. persimilis* ortholog *dpe_106* (**Figure 7F**). Analysis of the *D. pseudoobscura* population data indicated that all individuals were monomorphic for the ‘T’ allele (i.e. a fixed difference). We also observed several other non-seed positions of divergence within the star strand even within the *D. pseudoobscura* population, which is unusual and suggests fast evolution. As another example, we observe that both mature and star arms of *dps-mir-2542-1* exhibit multiple positions of seed divergence with its ortholog *dpe_101* (**Figure 7F**). Despite the impracticality of applying formal tests for evidence of natural selection given such a small sample, it is evident from these examples that several obscura TRC loci defy conventional behavior for purifying selection of seed regions and instead are quickly altering their seed regions between closely related species while still maintaining miRNA biogenesis.

#### 3. Preferential 3′ untemplated uridylation of mirtrons

Approximately 54% (232/428) of mirtron and tailed-mirtron annotations in *Drosophila* are new to our study, and are recently-emerged. This massive set of novel spliced miRNAs prompted us to ask if they exhibit characteristic properties of 3’-untemplated uridylation, as we reported in *D. melanogaster* [42,43]. In comparisons of 1777 canonical miRNAs and 289 mirtrons (**Supplementary Table S6**), we observed that mirtrons exhibited a massively greater rate of 3’ untemplated uridylation than canonical miRNAs (**Figure 7G, H**). This was evident on a per-locus basis on CDF plots (**Supplementary Figure S24A**), and was also the case even after conditioning on canonical miRNAs whose 3’ arm read ended in AG dinucleotide as with mirtrons. However, conditioning on 3′-AG enhanced the frequency of 3′ uridylation observed on canonical miRNA-3p species (**Supplementary Figure S24A**), consistent with the notion that the mirtron uridyltransferase Tailor has some intrinsic preference for hairpins terminating in 3′-G/AG [42,43].

Next, we examined whether the elevated frequency of uridylation at canonical miRNAs and mirtrons whose 3’ read ended with G was consistent across conserved and newly-evolved loci. For the conserved loci, we recapitulated previous signatures of uridylation (**Figure 7I**). Namely, mirtrons exhibited a significantly higher frequency of uridylation than canonical miRNAs (Mann-Whitney Test [M.W.T] p < 10^−21^), and comparisons among canonical miRNAs revealed that loci whose 3’ arm read ended with G were more uridylated than loci ending in a base other than G (i.e. IUPAC ‘H’ ambiguity character) (M.W.T p < 10^−12^). Of note, newly-evolved mirtrons and canonical miRNAs also exhibited the same signature as conserved loci (**Figure 7J**), and comparisons of loci within individual species revealed similar significant results in many species (**Supplementary Figure S24B**). Altogether, these findings from small RNA sequencing across the Drosophilid phylogeny broadly support the notion that adventitious access of splicing-derived hairpins to Dicer (i.e., mirtrons) continually creates a cohort of largely undesirable miRNA substrates, that are suppressed via Tailor-mediated uridylation that is in part sensitive to terminal hairpin “G”.

## Discussion

### A deep and broad empirical analysis of miRNA flux across the *Drosophila* genus

We are now 15 years into molecular evolutionary analysis of *Drosophila* miRNAs, but until now, there have been only limited attempts to address using the empirical data sampled throughout the *Drosophila* genus. In this study, we extended our curation of miRNAs from ∼1.9 billion *D. melanogaster* sRNA reads by sequencing ∼1.5 billion sRNAs from 11 other *Drosophila* species, assessing a diversity of samples optimized for miRNA discovery. Beyond the first experimental cloning of 592 orthologs of conserved miRNAs, we used a multitude of annotation pipelines and rigorous scoring criteria to annotate 649 completely novel miRNAs across 12 genomes. We carefully assessed these for ortholog relationships as a foundation for assessing evolutionary flux, including meticulous manual assignment of orthologs, paralogs, and newly-emerging members of genomic clusters (i.e. rapidly evolving TRC loci).

Overall, these data yield myriad insights into miRNA loci that are hidden from genomic alignments. These include the surprising existence of novel conserved miRNAs, unexpected clade-specific shifts in processing register, and post-transcriptional modifications of miRNAs. Beyond conserved loci, we uncover hundreds of “young” miRNAs that could not be identified by genomic sequence alone. These data allow us to quantify distinct rates of miRNA flux according to biogenesis type, genomic locale, tissue restriction, and evolutionary clade. We identify patterns of structural change and associated with flux in expression of evolutionarily nascent canonical miRNAs, providing a mechanistic basis for their instability. We also develop a new phylogenetic model to characterize rates of small RNA evolution in the presence of annotation uncertainty. Overall, we solidify the perspective that miRNAs do not comprise a unitary class, but encompass a diversity of functional loci with distinct evolutionary imperatives.

While our study provides the deepest perspective of miRNA evolutionary novelty across a genus to date, clear challenges remain for the future. We previously described evolutionary nascent miRNA-like loci in *D. melanogaster* that defy clear annotation by current standards [28,29]. In the future, analysis of a greater diversity of Drosophilid tissue and cell samples, combined with AGO1-IP sequencing, will be critical to interpret the earliest stages of miRNA emergence. At the same time, it worth appreciating that the current depth and breadth of sequencing is sufficient to identify hundreds of species-restricted, and even species-specific miRNAs with high confidence.

### Divergent rationales for rapid evolution of mirtrons and TRC miRNAs

Amongst our extensive collection of recently-emerged miRNAs, we discern two major subclasses of rapidly evolving loci, splicing-derived miRNAs (i.e., mirtrons) and testis-restricted clustered (i.e., TRC) miRNAs. We propose divergent functional explanations for their distinct evolutionary behavior, relative to the bulk collection of recently-emerged miRNAs that either evolve under mild purifying selection or lack substantial utility and evolve neutrally [8].

Mirtrons mature via the dominant non-canonical mechanism that bypasses the Drosha/DGCR8 “Microprocessor”, which otherwise serves as a molecular gatekeeper for generation of specific and accurate Dicer substrate hairpins. Mirtrons occasionally yield regulatory species that incorporate into beneficial regulatory networks, but the vast majority are not retained during evolution. We hypothesize that most mirtrons are adventitious Dicer substrates whose net regulatory capacity may be undesirable. By analogy, expressing synthetic, random, siRNAs are often used as a control situation, but probably one can imagine that they would impart fortuitous gene regulation that would not be tolerated over evolution. Indeed, molecular mechanisms involving uridylation have recently been shown to selectively suppress splicing-mediated miRNA biogenesis, which can accelerate the evolutionary flux of these miRNA substrates [42,43].

Our current studies across the *Drosophila* genus broadly confirms that the accelerated evolutionary dynamics of mirtrons is well-correlated with their tremendously high rates of 3′ uridylation and evolutionary turnover. Indeed, only six of the >400 mirtrons we annotated across the Drosophilid phylogeny were present in the fruitfly ancestor. We and others recently showed that the mechanism is mediated by the uridyltransferase Tailor, which has capacity to recognize hairpins bearing 3′-(A)G, which is characteristic for splicing-derived hairpins [42,43]. We hypothesized that this may have carryover effect on suppressing the evolutionary emergence of canonical miRNAs that happen to end in 3′-(A)G. Indeed, our broad survey using de novo miRNA annotation in eleven new *Drosophila* species provides evidence that newly-emerging canonical miRNA hairpins that end in 3′-G have more uridylation than ones than end in the other three nucleotides. This can explain the rapid evolutionary turnover of mirtrons, as well as the preferential uridylation of canonical miRNA-3p species ending in G of various evolutionary ages, which we observed to be depleted in the collection of deeply conserved, canonical miRNAs [42,43]. This supports the overall view of how molecular mechanisms that restrict the biogenesis of non-canonical miRNA substrates can affect the evolution of both non-canonical and canonical miRNA evolution.

We note also that almost all our annotations of the related 3′-tailed mirtrons, excepting *mir-1017*, have been lost from the Drosophilid phylogeny. In this case, since their hairpin tails are trimmed, they lack a characteristic 3′ nucleotide for potential identification. We have also annotated a class of confident novel antisense miRNAs, although excepting *mir-iab-4/mir-iab-8* and possibly *mir-307/mir-307AS*, it appears that all other instances of these have been purged over evolution. Thus, each of these specialized biogenesis pathways seems largely to exist to manufacture one or perhaps two miRNAs, even though biochemical capacity to make more exists in most species. Therefore, we speculate that there may be additional strategies to suppress miRNA biogenesis than we currently appreciate, especially that are suited to recognize aberrant miRNA substrates. For example, at least some 3′ tailed mirtrons are associated with untemplated 3′ additions (**Supplementary Figure S12**). It remains to be seen if this is also associated with an inhibitory role as with conventional mirtrons, and if so, how 3′- tailed substrates are recognized.

On the other hand, the extraordinarily rapid dynamics of TRC miRNAs in all subclades of the *Drosophila* genus provides strong evidence for their positive selection and adaptive evolution. Not only do TRC miRNA sequences evolve more quickly than canonical miRNA substrates of matched age, the total flux in TRC miRNA numbers between *Drosophila* subclades outpaces that of canonical miRNA loci. For example, a clear sequence ortholog of 106/497 canonical miRNAs not in the TRC class were present in the pan-Drosophilid ancestor, whereas this is only true of 13/265 TRC miRNA loci (**Figure 6C**). We entertain an alternative interpretation that some TRC miRNAs, owing to positive selection, have evolved in primary sequence so quickly that their ancestral relationships are not possible to assess. In any case, it is clear that the wholesale appearance and disappearance of massive TRC loci in different clades reflects a fundamentally different usage of these miRNAs than for maintenance of conserved seed-driven target networks as with typical canonical miRNAs.

Moreover, the atypical dynamics of TRC miRNAs are dramatically accelerated in both species examined in the obscura subclade. In fact, *D. pseudoobscura* and *D. persimilis* themselves exhibit substantial differences in their TRC repertoire, underlying nearly an order of magnitude greater birth estimate in the obscura branch than other branches of the phylogeny. The functional underpinnings of this remain to be tested, but they go hand-in-hand with the observation of dramatic proliferations of testis-restricted AGO2 paralogs specifically in the obscura subclade, and not in other Drosophila subclades [44,45].

Overall, our study provides a wealth of small RNA data that can guide functional studies of miRNA biogenesis, regulation of miRNA processing, and will underlie discovery of novel small RNA types (such as siRNAs and piRNAs). In addition, our deep and broad sampling across an entire genus provides myriad insights into the distinct evolutionary trajectories of multiple miRNA subtypes, affirming that miRNAs cannot be considered a unitary class with respect to their functional impact and utilization.

## Materials and Methods

### *Drosophila* samples and small RNA library sequencing

To analyze miRNA evolution in *Drosophila* species, we obtained cultures of whole-genome sequenced *D. simulans, D. sechellia, D. yakuba, D. erecta, D. ananassae, D. pseudoobscura, D. persimilis, D. willistoni, D. virilis*, and *D. mojavensis* strains from the UCSD *Drosophila* Species Stock Center. Adult *D. grimshawi* samples were a gift of Dr. Kevin White (University of Chicago). Small RNAs (∼18-28nt) were isolated from male bodies, female bodies, heads, and mixed embryos using polyacrylamide gel electrophoresis, and we prepared libraries as described. Libraries were sequenced on Illumina GAxII or Hi-Seq 2000 instruments.

### Recovery of *D. melanogaster* miRNAs from simulated libraries

We simulated four libraries each representing 25 million randomly sampled reads from the pooled collection of *D. melanogaster* sRNAs from male bodies, female bodies, mixed embryos, and heads respectively. We created 100 samples of 100M reads each (25M X 4 libraries), and determined the recoverability of conserved and newly-evolved *D. melanogaster* miRNAs at varying minimum mature and star read expression thresholds. These artificial libraries, allowed us to investigate the amount of miRNAs at two different age groups (i.e. conserved and newly-evolved) that could be recovered in the other 11 *Drosophila* genomes. We defined conserved *D. melanogaster* miRNAs as those with unambiguous orthologs in the obscura-group species, *D. willistoni*, or the *Drosophila*-group species. Recently-evolved *D. melanogaster* miRNAs were defined as those with orthologs within the melanogaster-group species only.

### Annotation of miRNA genes

To identify novel miRNAs and assess the expression levels of all miRNA loci, we first mapped reads from each of the 11 *Drosophila* species unto their reference genomes. All reference genomes, except for *D. simulans*, were obtained from Flybase. We utilized a revised *D. simulans* genome assembly created from an isogenic *w501* female within our analysis [46]. Reads were mapped using the Bowtie program by allowing for up to 3 mismatches (parameters: -v 3 -k 20 ‐‐best ‐‐strata). Perfectly mapped reads, and reads with 3’ end mismatches characteristic of untemplated additions, were used for the identification of miRNAs.

We supplemented existing *Drosophila* miRNA annotations from miRBase v21 with novel miRNAs and mirtrons identified in this study using a multi-stage pipeline [25]. First, canonical miRNA and mirtrons were predicted using miRDeep2 using default software settings [13]. To identify short mirtron and long pre-miRNA hairpins, two classes systematically missed by miRDeep2, we mapped sRNA datasets to introns obtained from FlyBase gene annotations, and predicted hairpin structures from the *einverted* program of the EMBOSS package [47] in a genome-wide manner per species. We used the invert_it.pl utility script from the ShortStack program [48] to filter the *einverted* results. The parameters specified to this script were: `-f 0.6 -p 30`. Introns or hairpin structures with at least one mapped read were retained and ranked by p-values calculated from a Random Forest classifier. We trained this classifier with a balanced set of positive training examples comprised of known *D. melanogaster* and *D. pseudoobscura* miRNAs download from miRBase (v21), and a negative training set composed on non-miRNA predictions identified manually in this study. We used a total of 37 features per training case representing sequence, structure, and sRNA read alignment features (**Supplementary Table S7**). Minimum free energy, and sub-optimal secondary structures were predicted using RNAfold and RNAsubopt in the Vienna RNA Software [49].

All miRNAs predicted from this pipeline were vetted manually and bioinformatically for miRNA and mirtron candidacy. In the manual phase, all miRDeep2 predictions, and intron and hairpin structures with p > 0.5 were examined for evidence of cleavage by Drosha and Dicer based on the sRNA read alignment, a hairpin secondary structure, and synteny with other miRNA predictions. Putative canonical miRNA were further classified bioinformatically using criteria based on (1) expression, (2) clonability of Drosha/Dicer products, such as the miR, miR*, loop, or MOR sequences, (3) structure pairing of the miR:miR* duplex, (4) 5’ end consistency of miR and miR* reads, and (6) ratio of background to miRNA reads (**Supplementary Figure S3**). Mirtrons were classified using the same features as for canonical miRNAs, but an additional criterion for untemplated modifications of the 3’ arm reads was specified. Canonical miRNA or mirtron predictions that met all criteria were labeled as “confident” while those that failed some or all criteria were labeled as “candidate” or “FALSE,” respectively. “Candidate” loci that were orthologous to “confident” annotations were re-classified as “candidate-rescued.” Confidence classifications for all miRNA and mirtrons are provided in **Supplementary Table S4**. Finally, novel “confident”, “candidate-rescued”, and “candidate” miRNAs and mirtrons were segregated from miRBase annotation and their unannotated orthologs.

### Identification of miRNA clusters and testes-restricted miRNAs

miRNA clusters were identified by grouping genes within a 10 Kb window of each other. Mirtrons were excluded from this classification. The majority of miRNA clusters identified in this study comprised genes with testes-restricted expression. Testes-restricted miRNAs were characterized as genes with > 4-fold log_10_(RPMM) testis or male-body expression enrichment when compared against all other tissue and developmental-timepoint libraries. If >75% of miRNA genes within a cluster were classified as testes-restricted, then all genes within said cluster were labeled <u>**C**</u>anonical, <u>**T**</u>estes-restricted, <u>**C**</u>lustered miRNAs.

### Identification of miRNA orthologs and alignments

miRNA orthologs were identified using the LASTZ program with the following parameters: H=2000 Y=3400 L=4000 K=2200 Q=HoxD55.q [50]. Hits were ranked by a score based on the consistency, continuity, and percent identity metrics from LASTZ. A 12-species sequence alignment were created for each miRNA prediction using best scoring orthologs and the Fast Statistical Aligner program [51]. Paralogs were a byproduct of this procedure because they attained lower rank during orthology assignments. All orthologs and paralogs were automatically included in our annotation pipeline, and were vetted by the same criteria.

Our process of collating accurate miRNA synteny information played a crucial role within our annotation procedure because it allowed us to “rescue” low-evidence, candidate miRNA and mirtron predictions contingent on the availability of one or more confident orthologs. Automated approaches were inept for identifying testes-restricted, clustered miRNA orthologs due to their close genomic proximity, low sequence conservation, and high rates of tandem duplication. For these miRNAs, orthologs and paralogs were identified from multiple sequence alignments of all members of a particular cluster in all species with homologs (see **Supplementary Figures S9** for examples). MiRNA alignments were further grouped into three classes: (1) TRC miRNAs, (2) solo canonical miRNAs, and (3) mirtrons.

### Birth and Death Model

To assess birth and death rate variation across classes of miRNAs and across *Drosophila* clades of interest, we designed and implemented a phylogenetic probabilistic graphical model. This model permits estimation of parameters of gene birth (*λ*) and death (*μ*) (**Figure 6A**) based upon our assignments of miRNA presence and absence in each species per miRNA family alignment. Parameter estimation required two sets of precomputed data. The first datum needed was a binary encoding of miRNA presence (1) and absence (0) as leaf node labels of the phylogenetic model. In this regard, we labeled non-miRNAs and “candidate” miRNAs as cases of absences, and “confident” or “candidate-rescued” miRNAs as cases of presences. We assigned each miRNA for which no orthologs were identified to its own singleton miRNA alignment, to be counted as an independent birth event. The second datum needed was phylogenetic branch-length estimates for the 12 *Drosophila* species phylogeny [27]. We estimated branch lengths (τ) in units of substitutions per site, using fourfold degenerate sites (i.e. sites within a codon in which all four possible nucleotide substitutions are synonymous) and the maximum-likelihood program RaXML [52]. Fourfold degenerate sites were extracted from a *de novo* 12 *Drosophila* species whole genome alignment constructed using the LASTZ and MULTIZ programs and the chaining and netting protocol used for the UCSC Genome Browser. The resulting maximum-likelihood newick-formatted tree was: ((((((dm3:0.055153,(droSim1:0.027716,droSec1:0.023941):0.024430):0.050893,(droYak 2:0.090814,droEre2:0.079010):0.032754):0.328435,droAna3:0.466508):0.162763,(dp4:0.018457,droPer1:0.018684):0.407262):0.120336,droWil1:0.593093):0.118858,((droVir3: 0.244781,droMoj3:0.335567):0.082788,droGri2:0.319783):0.118858);

Given these two datasets, we used our model to infer maximum-likelihood parameter estimates (i.e. *λ, μ*) using the standard belief-propagation algorithm to compute likelihoods, and the Broyden-Fletcher-Goldfarb-Shanno (BFGS) algorithm to obtain maximum-likelihood parameter estimates [53]. Parameter estimates were computed for the merged miRNA collection (final estimate: *λ* = 0.292, *μ* = 0.694), which we later used to compute (1) ancestral gene presence or absence states, and (2) probabilities of observable edge-wise birth and death events (**Supplementary Figure S18-21**). To assess cumulative counts of observable birth and death events per miRNA class or Drosophila clade, we computed edge-wise joint posterior probabilities (i.e. P(child, parent)) by belief propagation. For simplicity, we called birth [P(1, 0)], death [P(0, 1)], and “no change” events [P(0, 0) or P(1, 1)] if these probability estimates were >= 0.5 (**Figure 6B-D**). This method is implemented as a Java software package and available upon request.

## Data access

*Drosophila* sRNA sequencing data are under submission for access via the NCBI Gene Expression Omnibus. Read pileup, structure prediction, and 12-fly sequence alignments of all *Drosophila* miRNAs are provided as Supplementary material via an online website (http://compgen.cshl.edu/mirna/12flies/12flies_alignments.html).

## Acknowledgements

We thank Matthew DeCruz for technical assistance and the UCSD *Drosophila* Species Stock Center for fly stocks. Work in E.C.L.’s group was supported by the National Institutes of Health (R01-GM083300 and R01-NS083833) and MSK Core Grant P30-CA008748. J.M. was supported in part by the Tri-Institutional Training Program in Computational Biology and Medicine (via NIH training grant 1T32-GM083937).

**Mohammed and Flynt et al**

**Deep experimental profiling of microRNA diversity, deployment, and evolution across the *Drosophila* genus**

## Supplementary Figures and Tables

**Supplementary Table S1:** Small RNA libraries for 11 *Drosophila* species created for and analyzed in this study.

**Supplementary Table S2:** Small RNA libraries for *Drosophila* species acquired from public repositories and analyzed in this study. (**A**) Libraries for *D. melanogaster*. (**B**) Libraries for *D. simulans, D. yakuba, D. erecta, D. pseudoobscura*, and *D. virilis*.

**Supplementary Table S3:** miRBase Drosophilid loci that were demoted from miRNA status.

**Supplementary Table S4:** Master list of all known and novel miRNAs in *Drosophila*.

**Supplementary Table S5:** List of all sense and antisense miRNA pairs identified and analyzed in this study.

**Supplementary Table S6:** 3′ Untemplated nucleotide addition counts to canonical miRNA-3p and mirtron-3p species.

**Supplementary Table S7:** miRNA sRNA read, sequence, and structure features for Random Forest and logistic regression classifiers.

## Supplementary Figures Legend

**Supplementary Figure S1:**
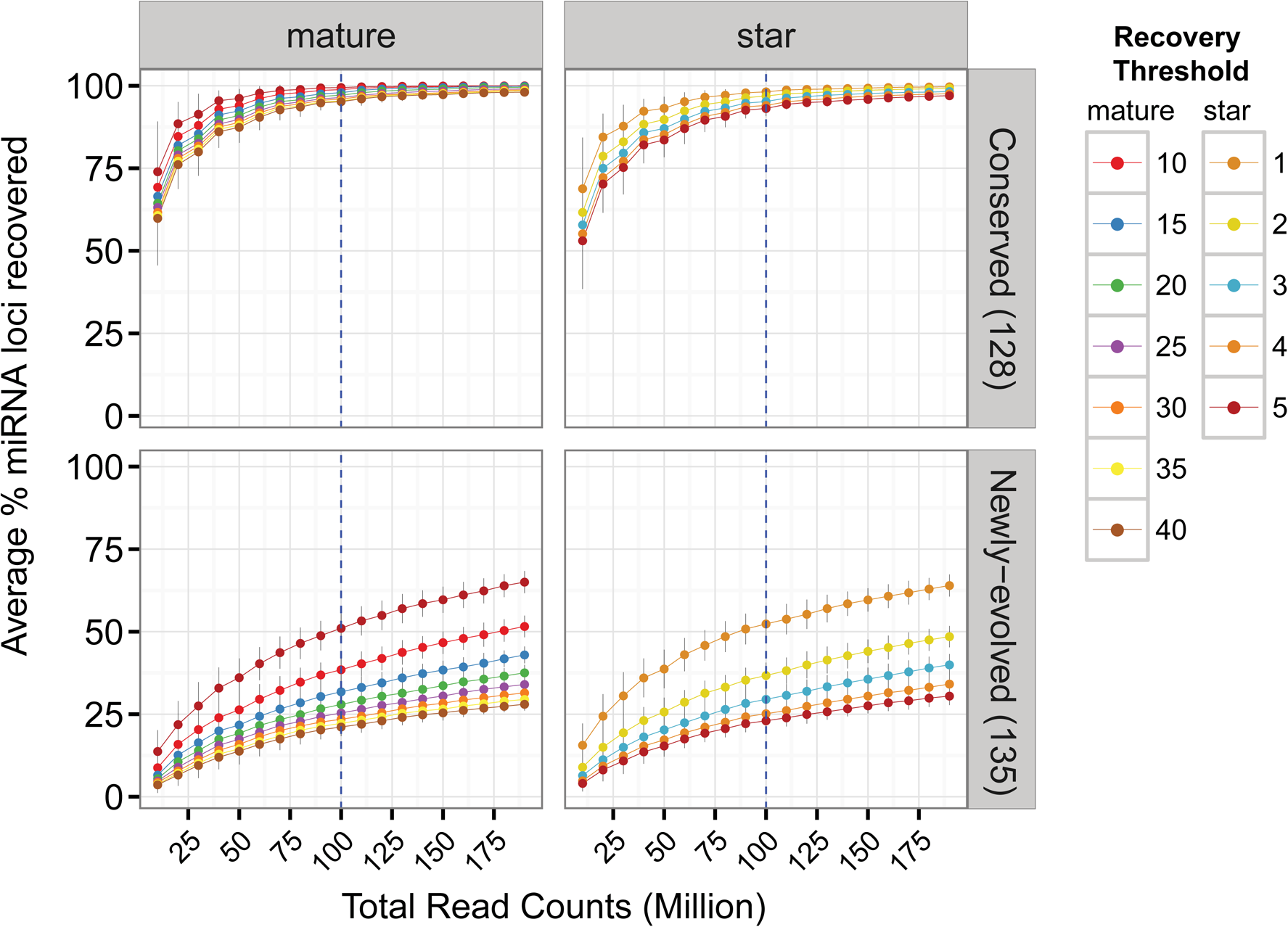
Conserved and newly-evolved *D. melanogaster* miRNAs recovered at varied read depth thresholds using *in silico* simulated libraries. These libraries are composed of randomly sampled reads across all *D. melanogaster* sRNA-seq male-body, female-body, head, and mixed embryo libraries used within this study. miRNA recovery rates are computed per read-depth sample at various miR or miR* read thresholds. Error bars depict the standard error of the recovery rate across 100 simulations.

**Supplementary Figure S2:**
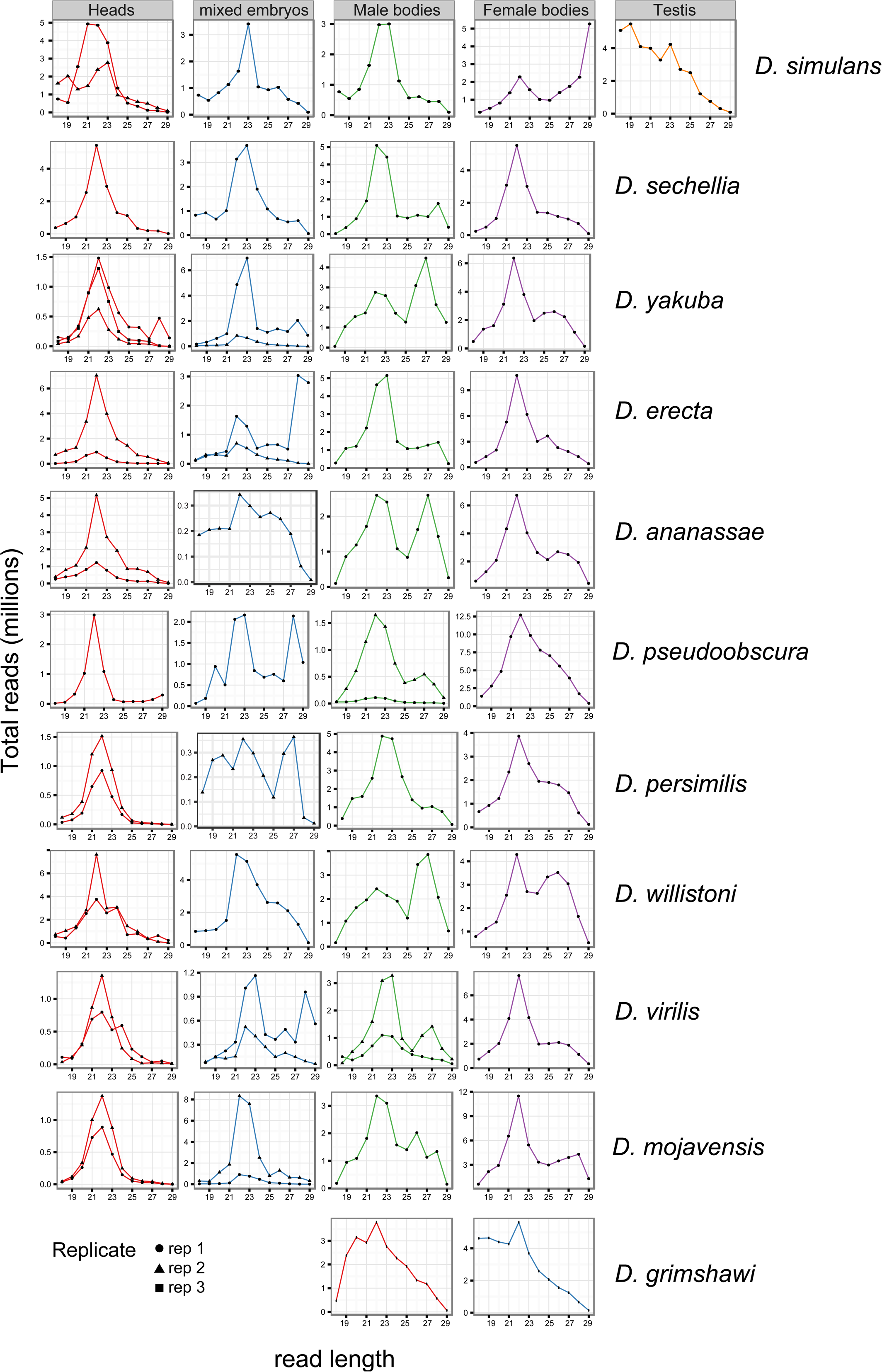
Read length distribution for all small RNA libraries sequenced in this study. We extended our previous broad and deep analysis of D. melanogaster by sampling 11 additional Drosophila species as listed to the right, by analyzing mixed embryos, adult heads, male bodies and female bodies; a testis library was also generated for D. simulans. A subset of libraries were sequenced in replicates, especially ones where the expected dominant miRNA-sized peak (21-22 nt) peak was not initially observed. A piRNA peak is seen in most of the body libraries. Due to the technical difficulty in culturing D. grimshawi, it was only feasible to generate two libraries for male and female bodies.

**Supplementary Figure S3:**
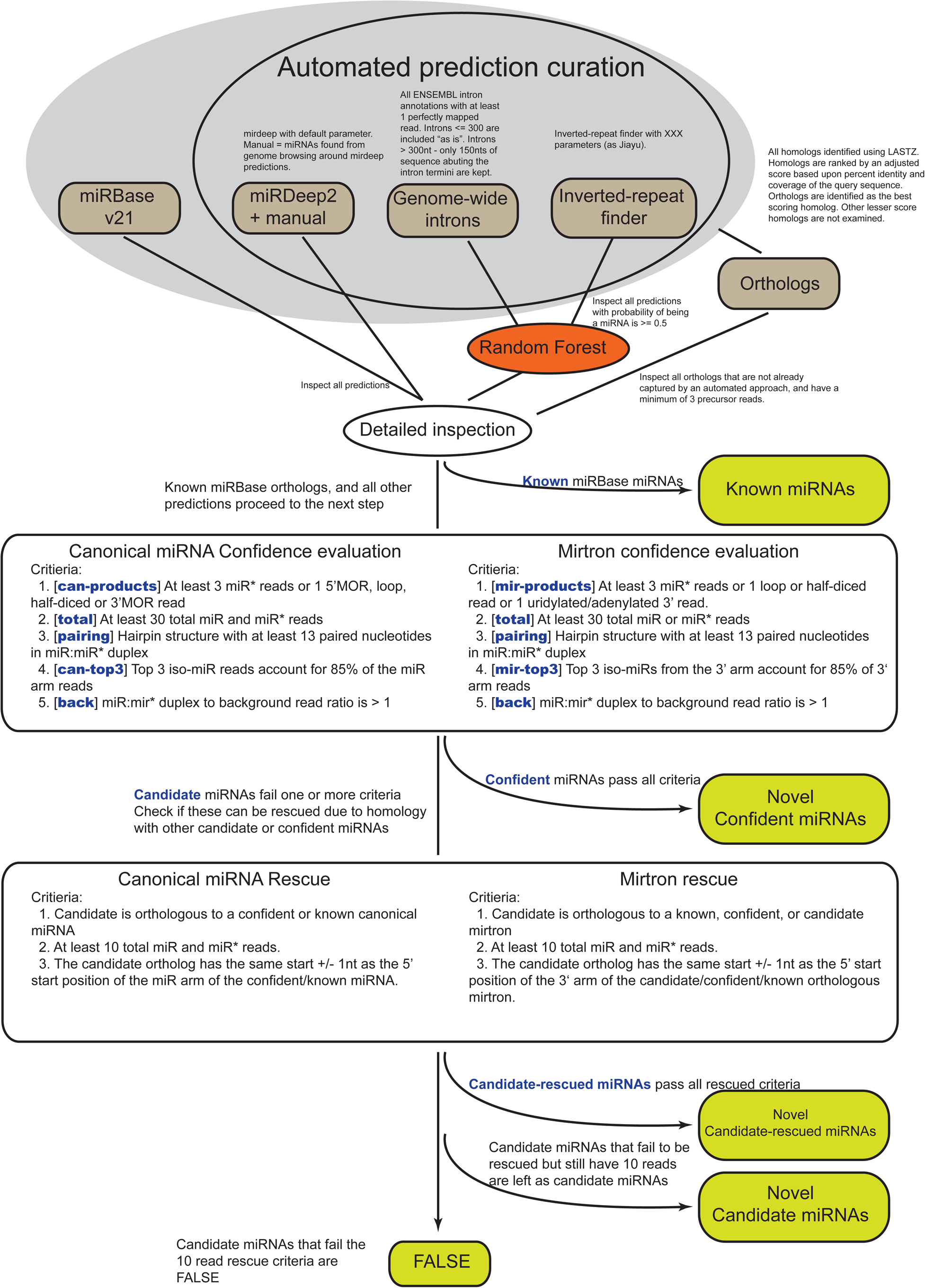
Detailed flow-chart of miRNA and mirtron identification pipeline and scoring criteria.

**Supplementary Figure S4:**
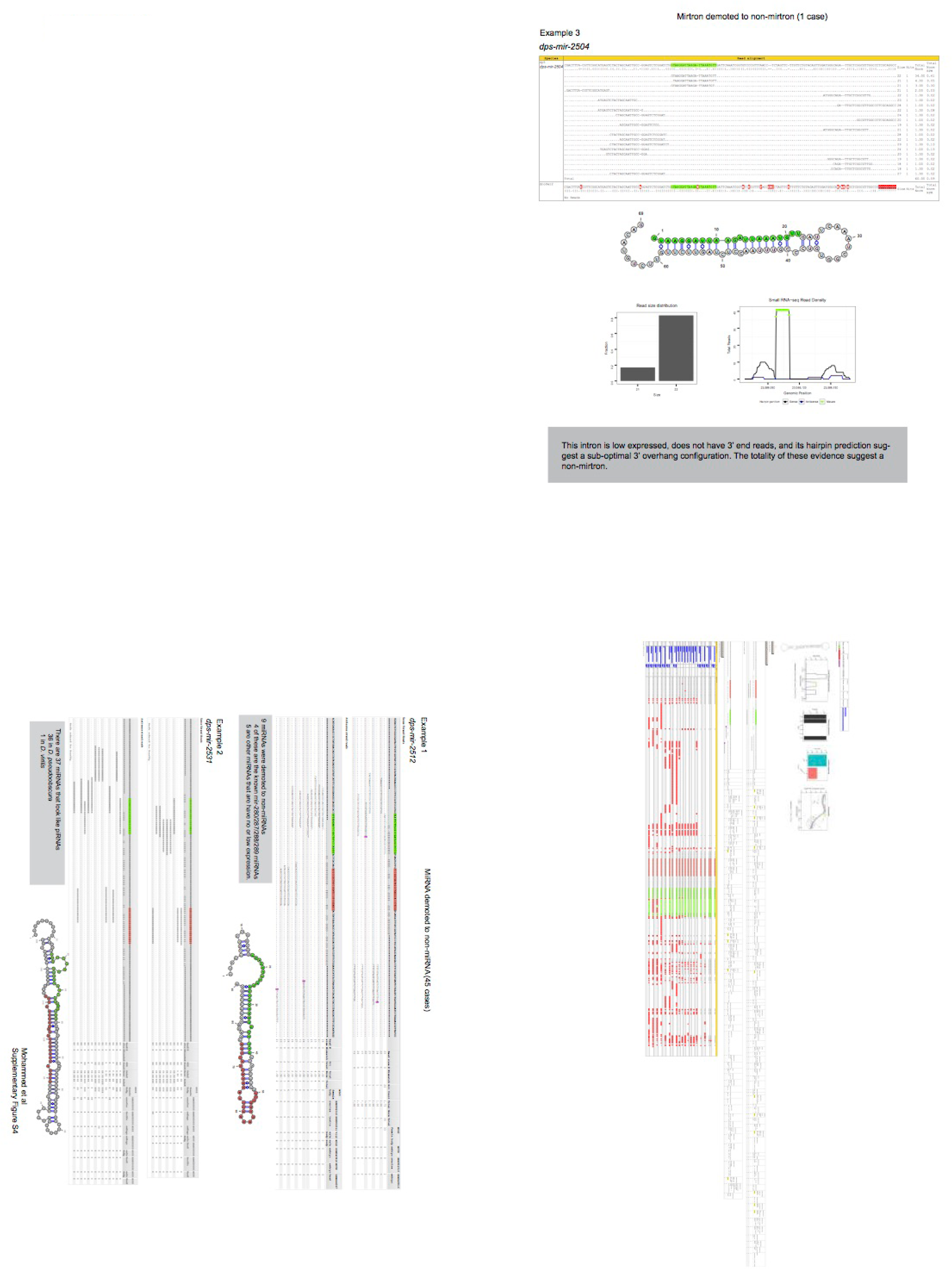
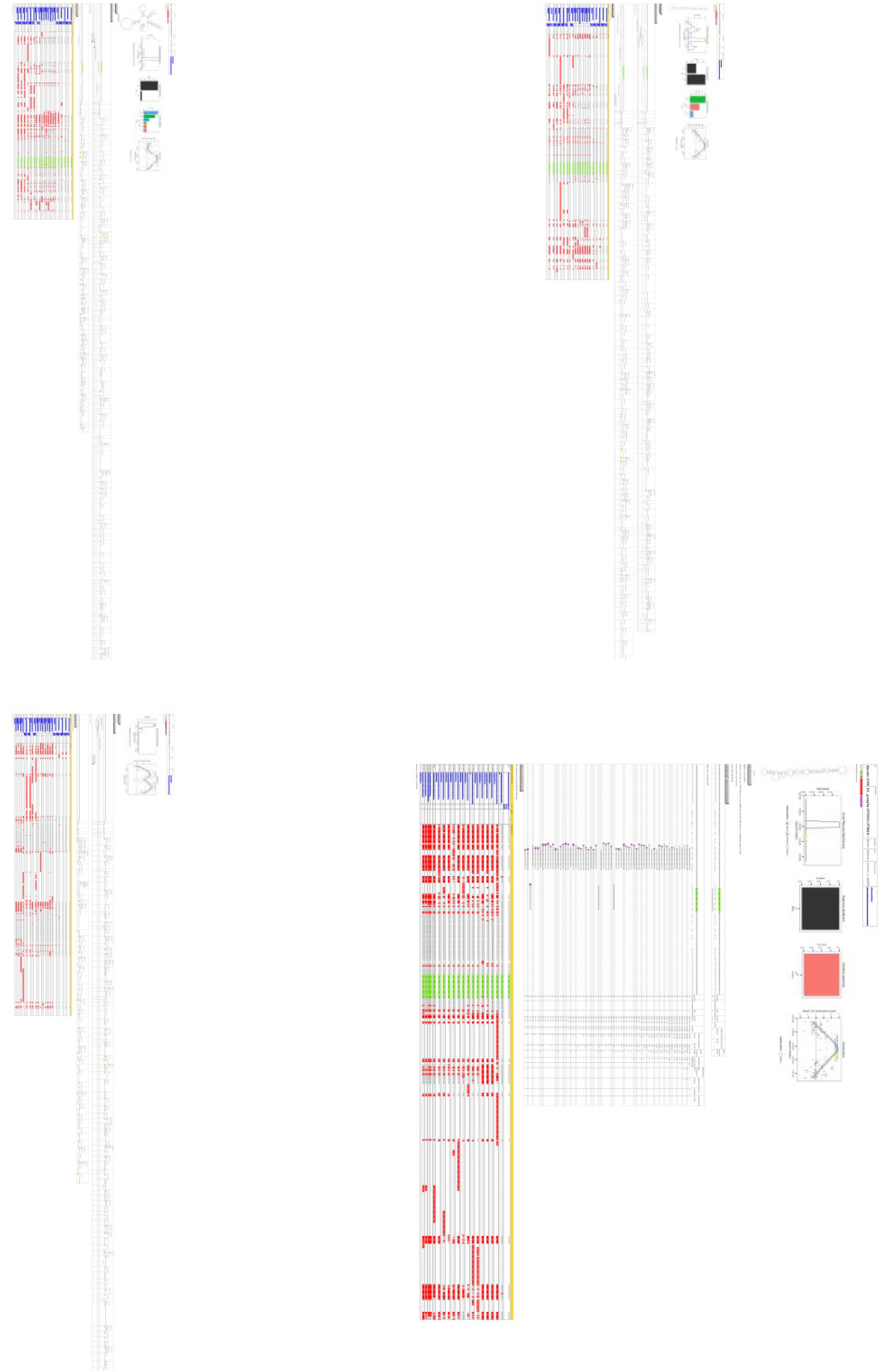
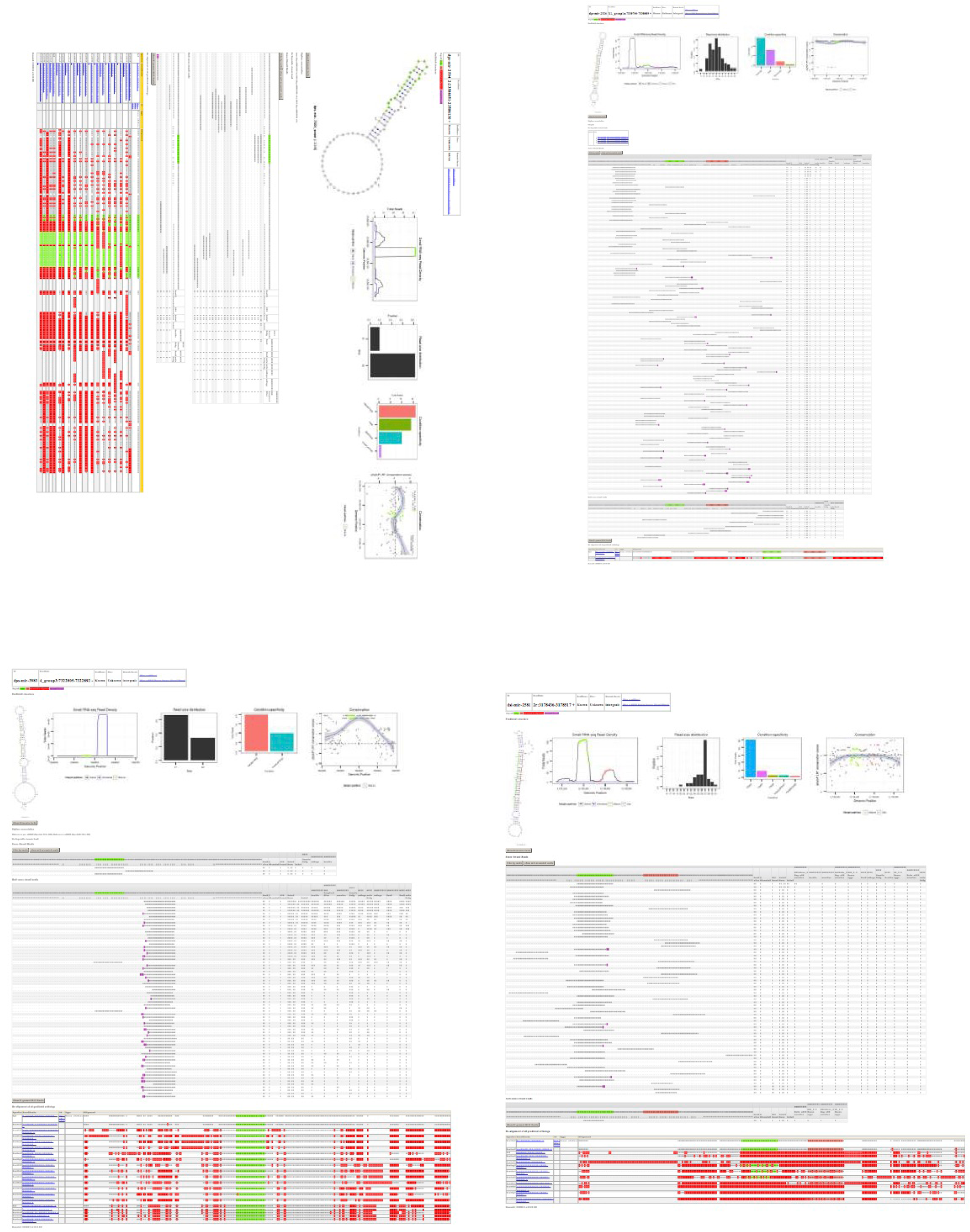
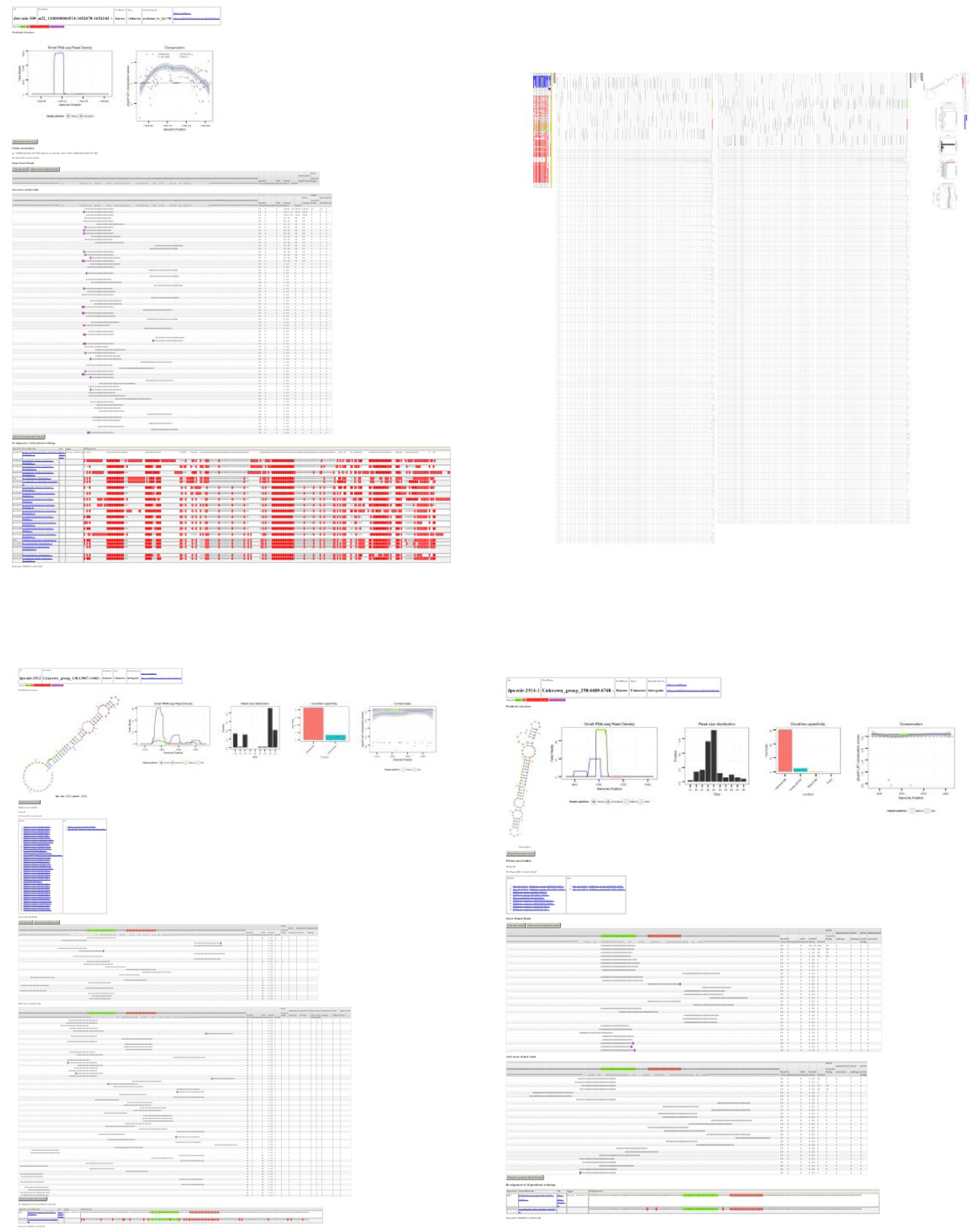
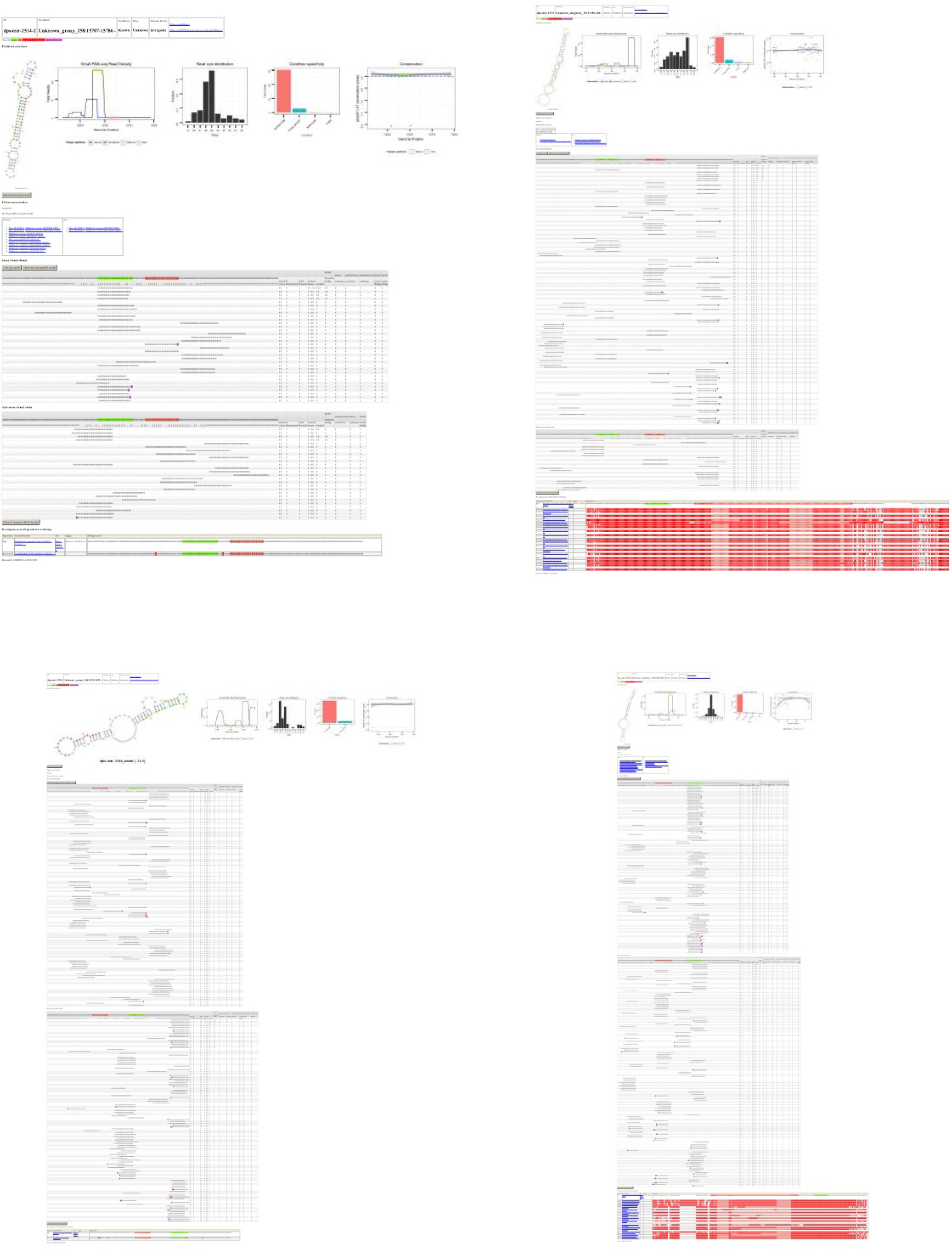
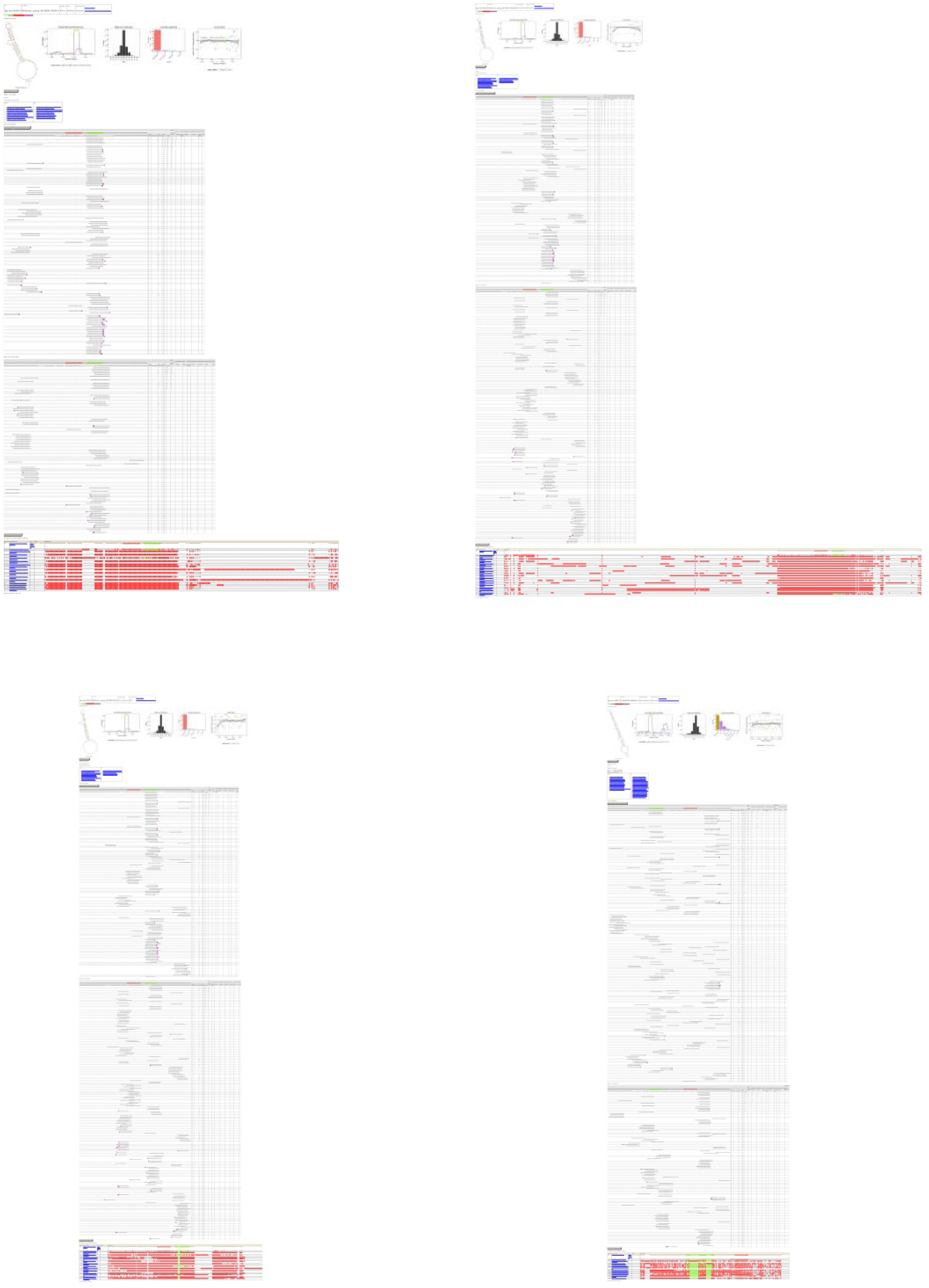
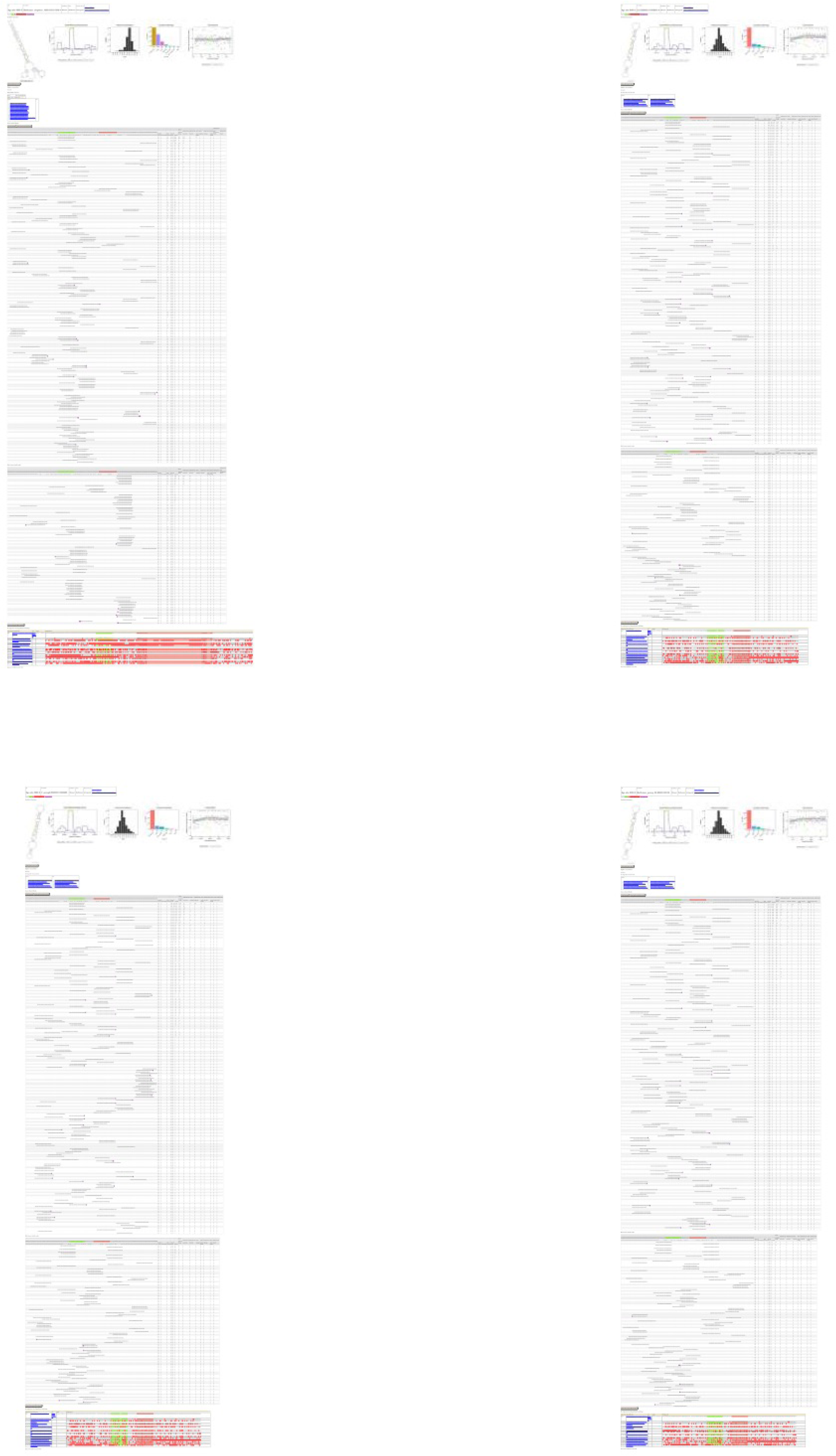
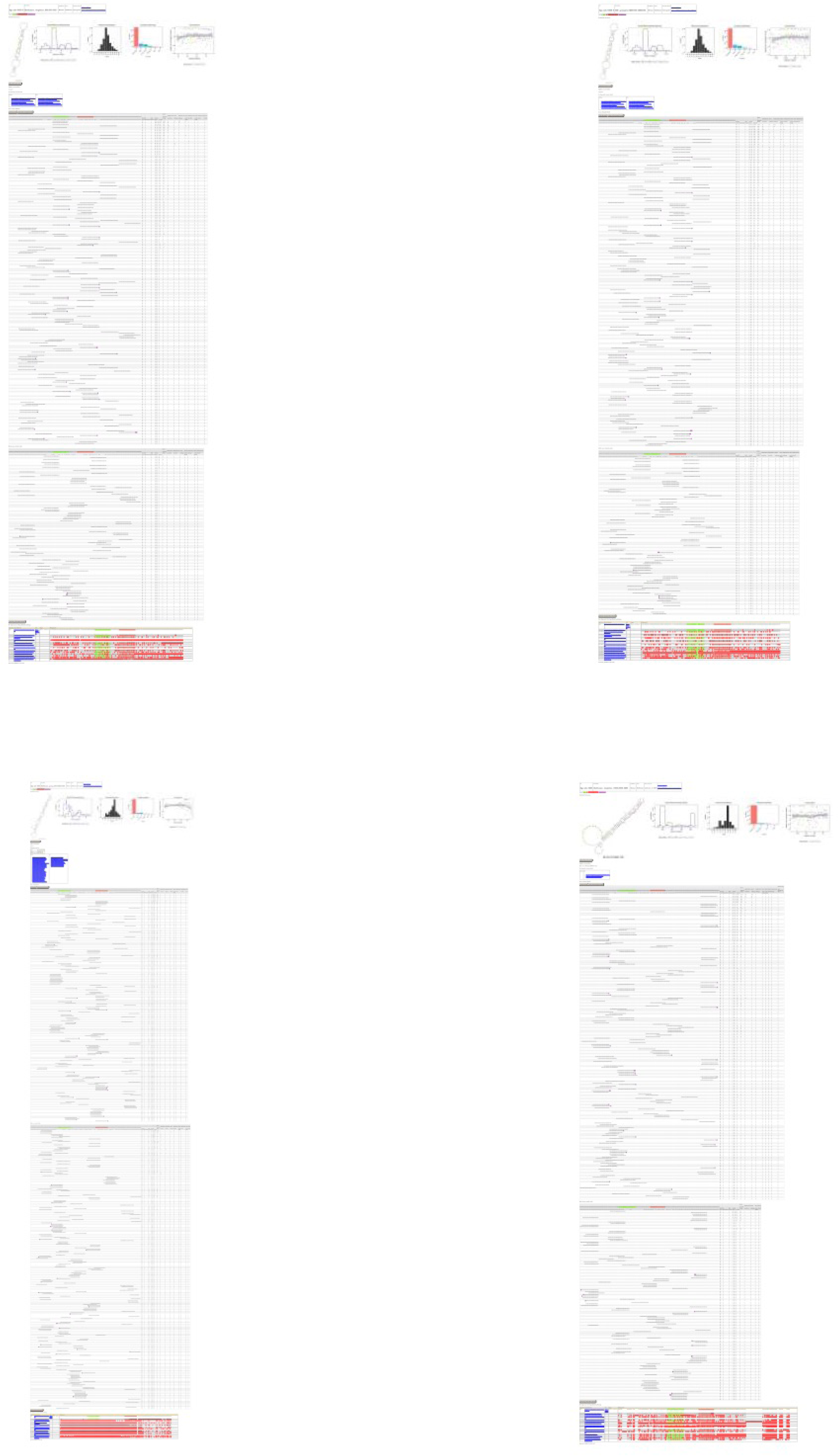
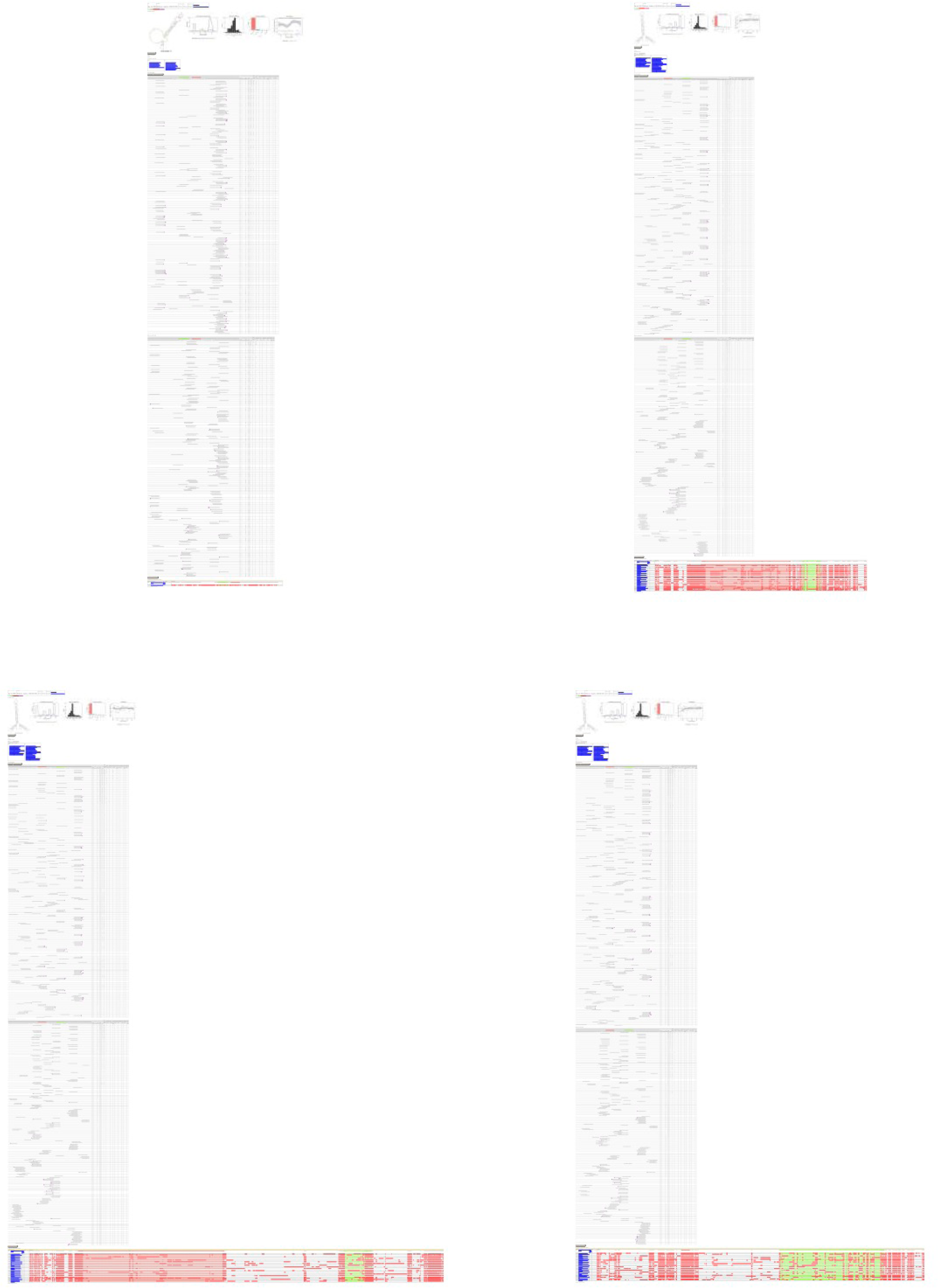
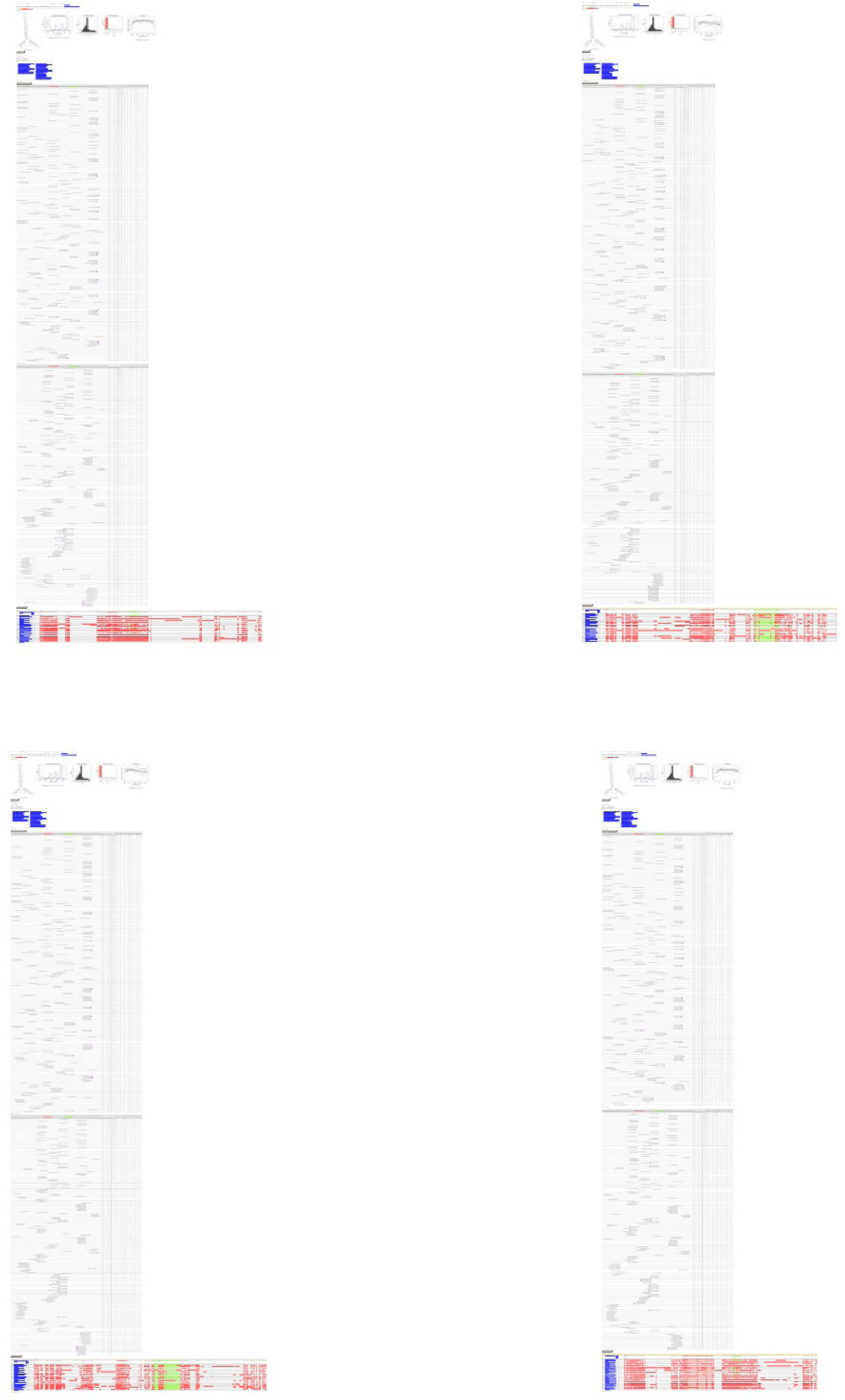
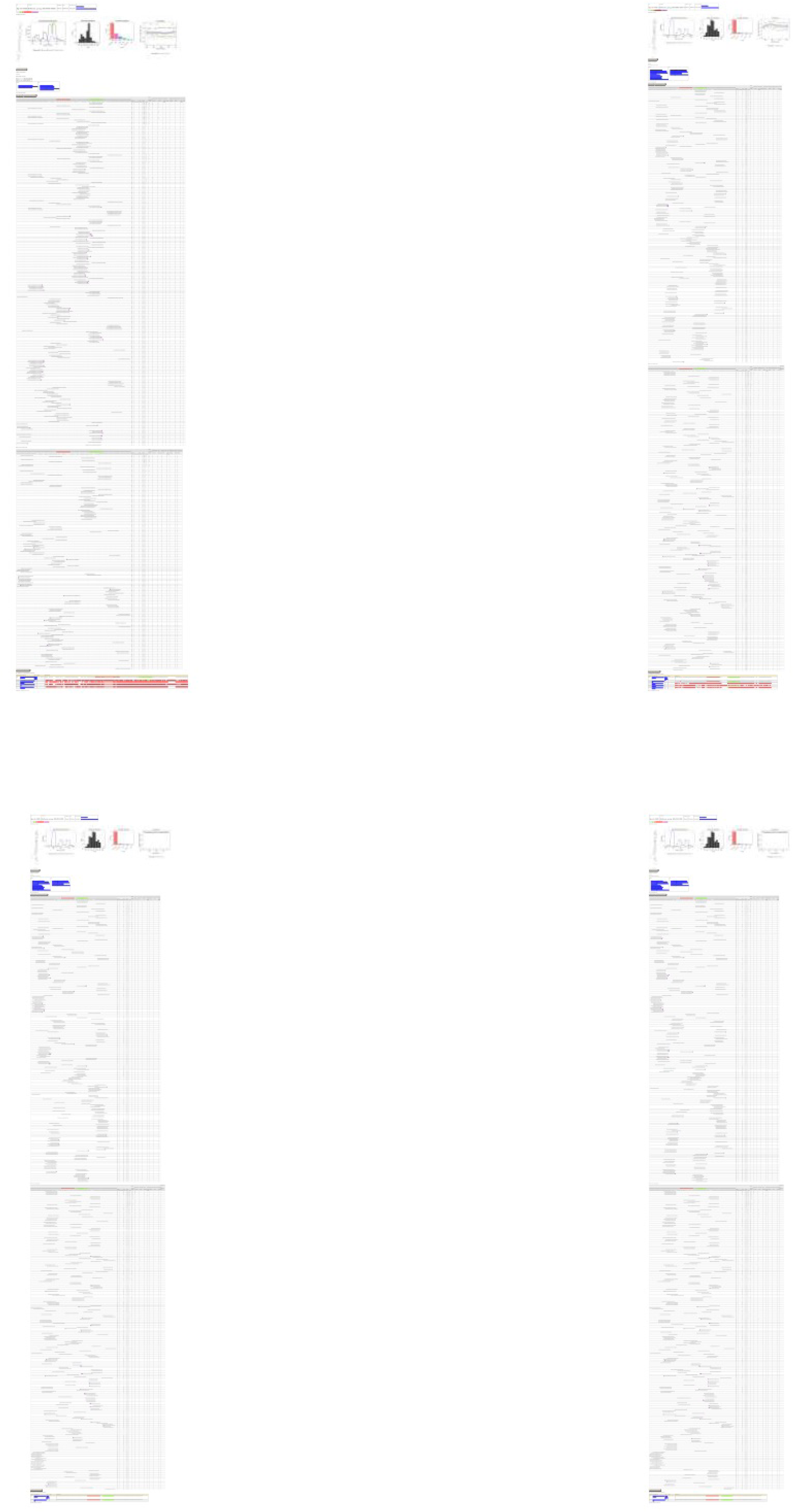
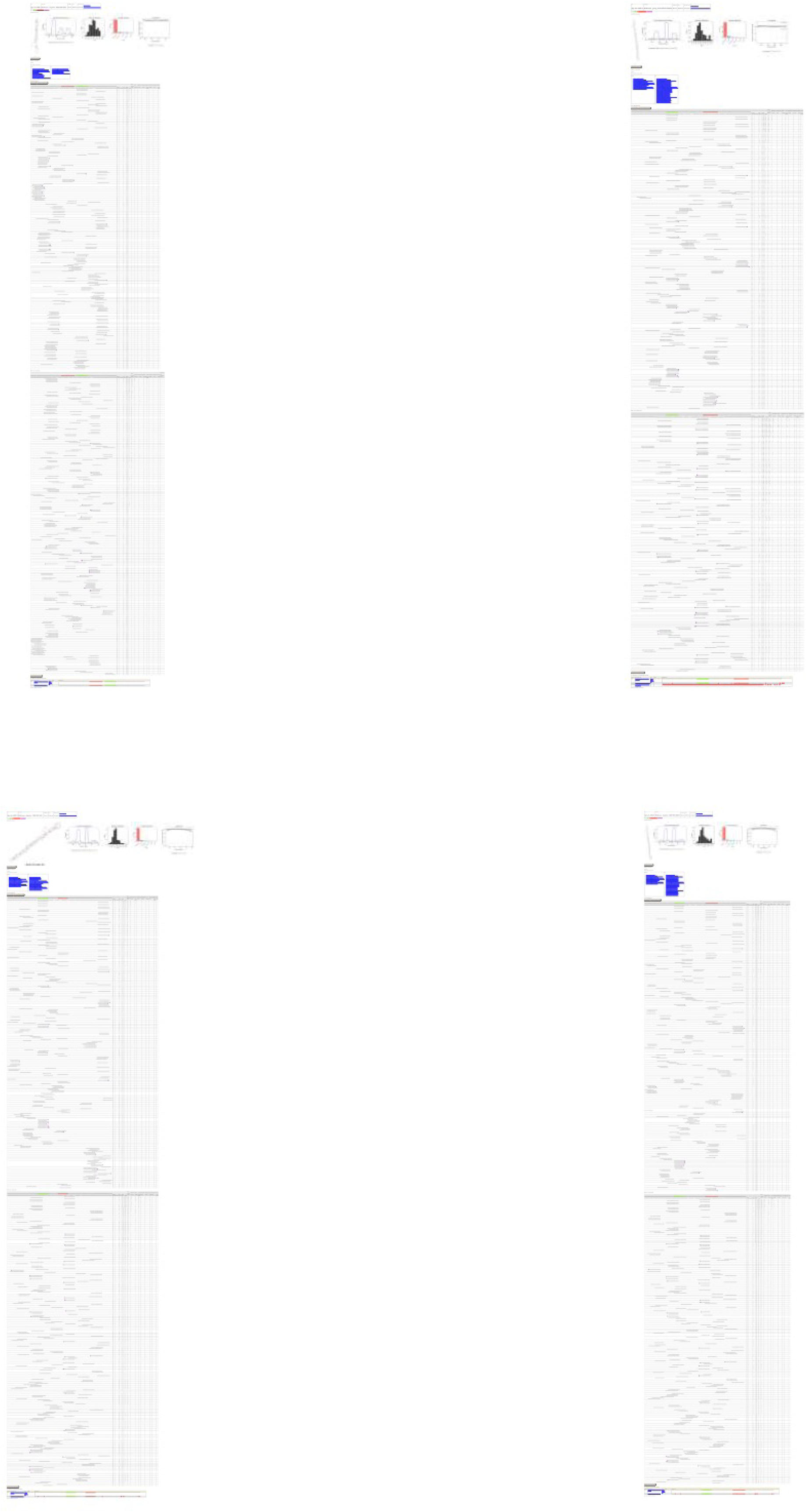
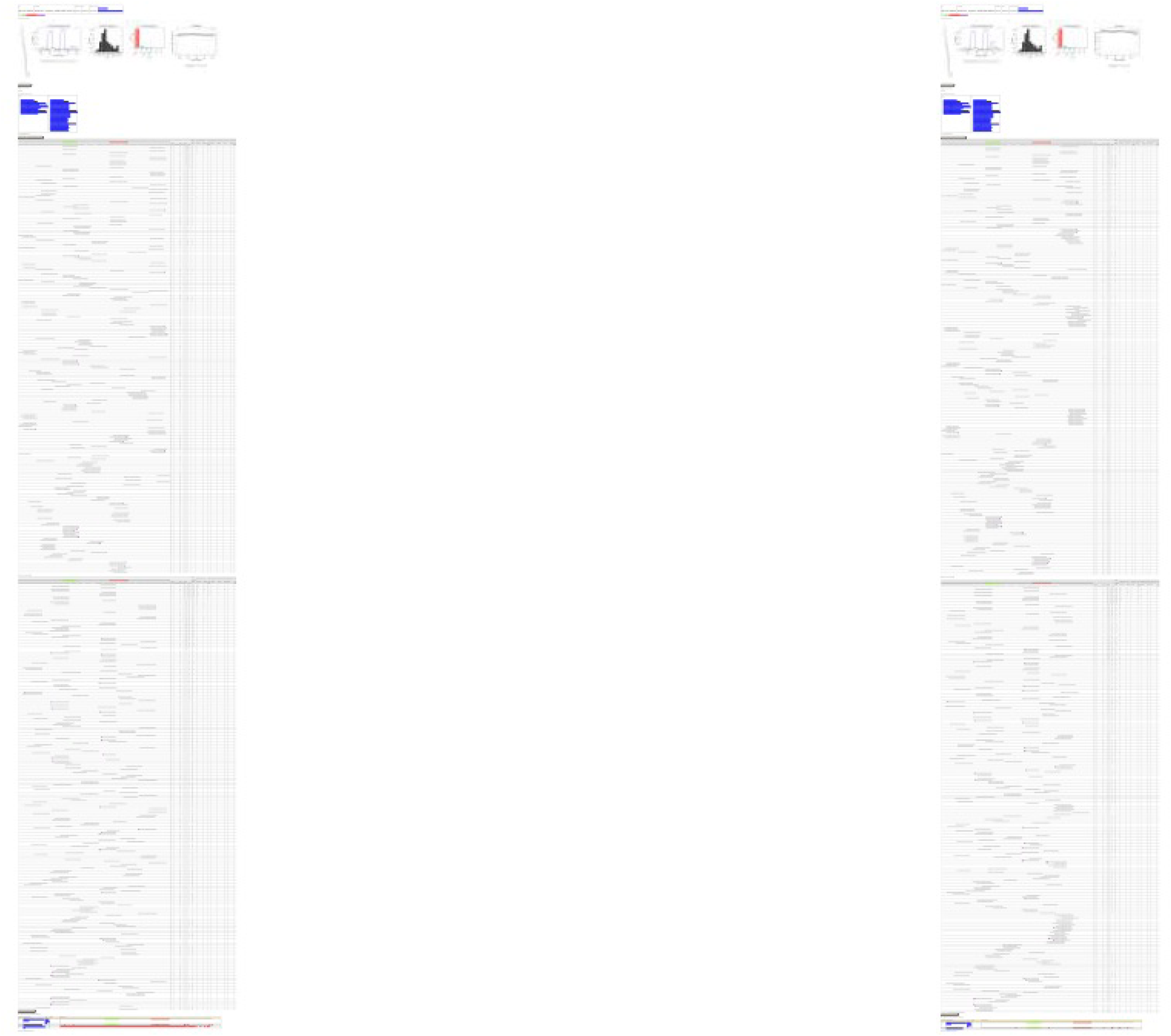
Drosophilid miRbase loci demoted for lack of compelling small RNA evidence. Examples of 47 demoted miRBase (v21) miRNAs and mirtrons. 37 of these represent piRNAs within *D. pseudoobscura* and *D. virilis* previously classified as miRNAs.

**Supplementary Figure S5:**
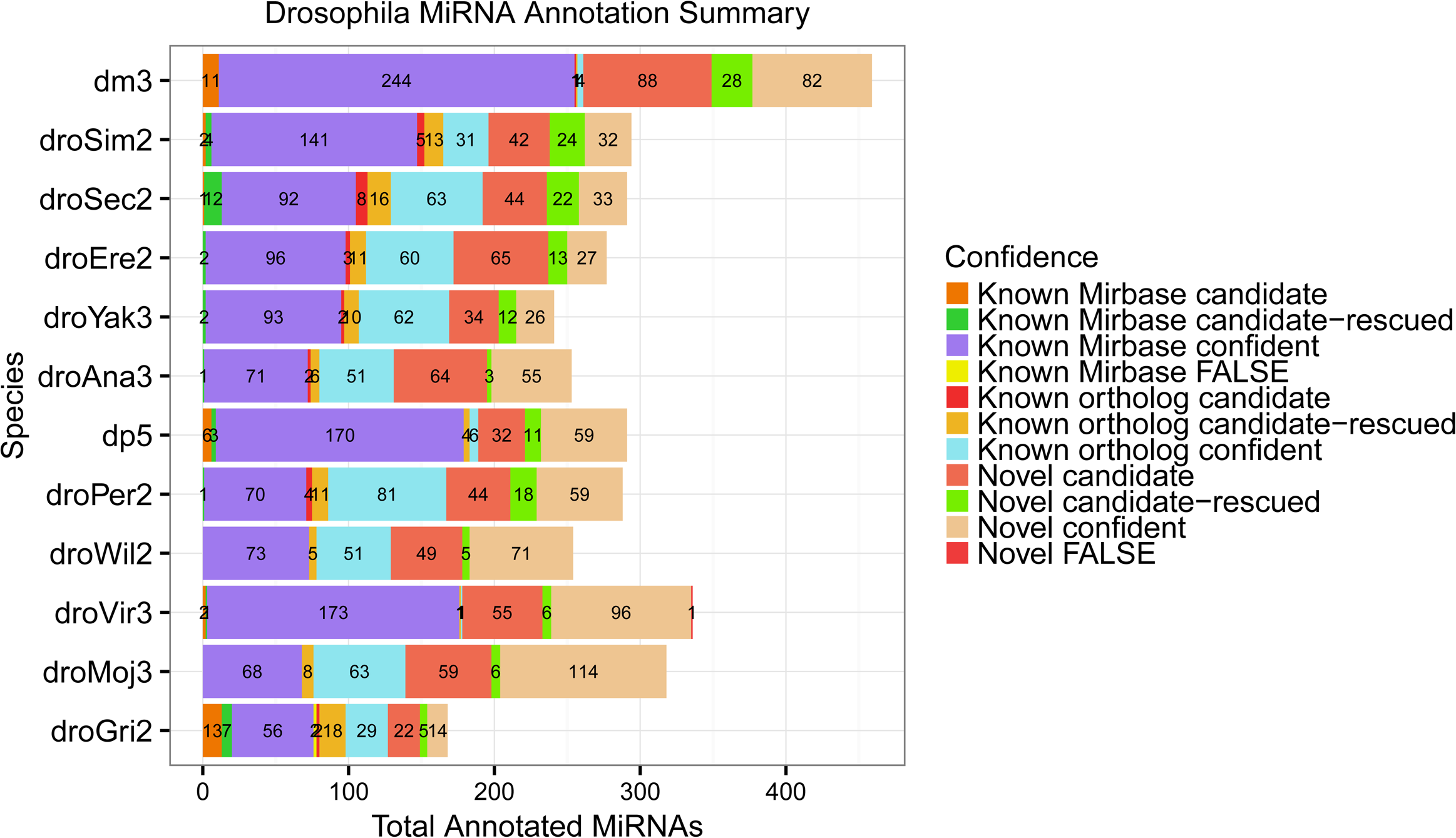
Total miRNA and mirtron annotation count within 12 *Drosophila* species. Annotations are further subdivided within (1) three confidence categories- “confident”, “candidate-rescued, and candidate”, and (2) between known and novel annotations. Note that “candidate” annotations were not utilized for analyses of miRNA flux in this study.

**Supplementary Figure S6:**
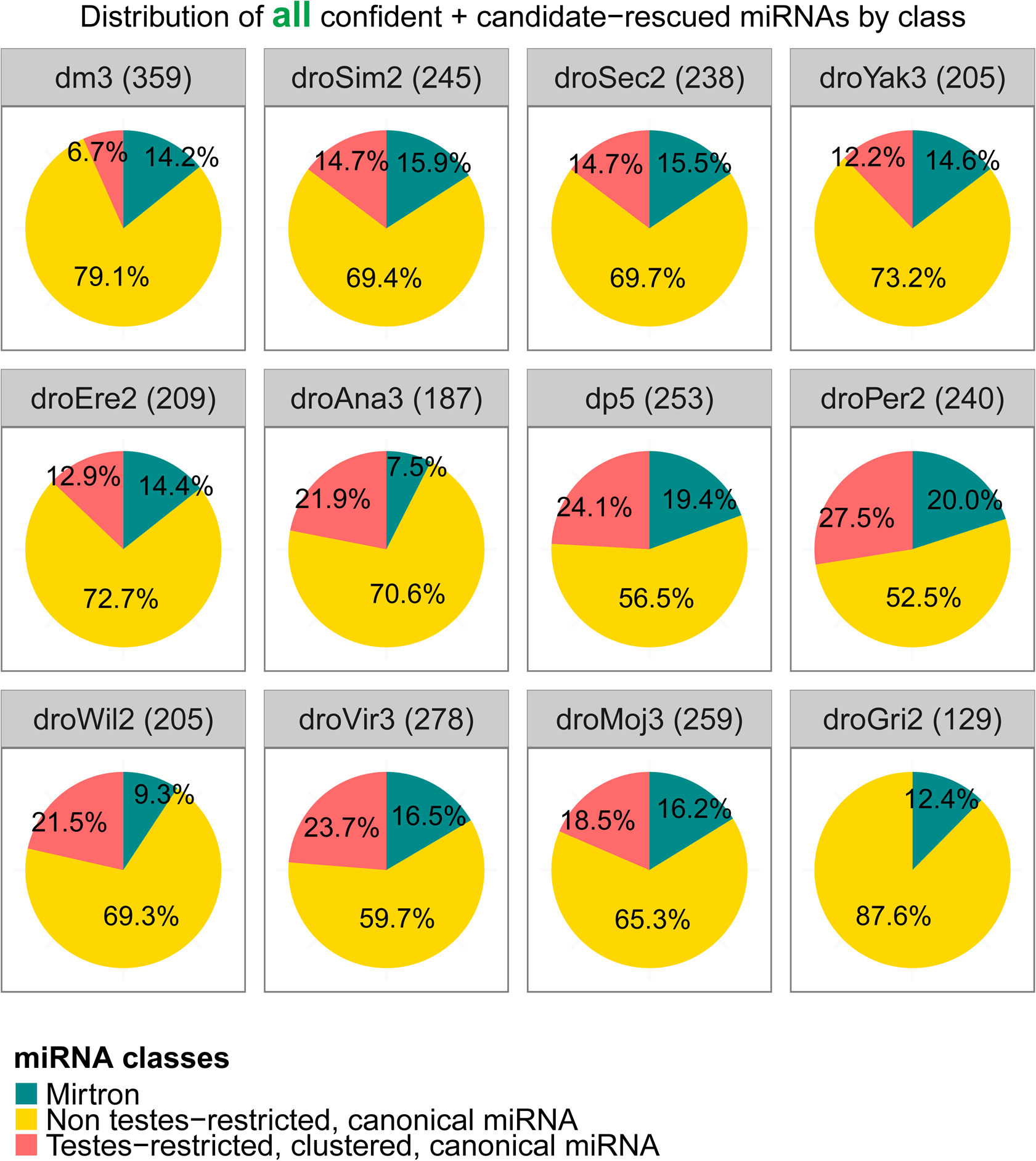
Distribution of miRNAs for three classes of miRNAs within each Drosophila species. These classes are defined by biogenesis pathway and canonical miRNAs are further divided by their testes-restricted expression. Only “confident” and “candidate-rescued” loci are included; loci that are considered “candidate” only and lack further rationale to be rescued based on a confidently processed miRNA ortholog are not included in these pie charts.

**Supplementary Figure S7:**
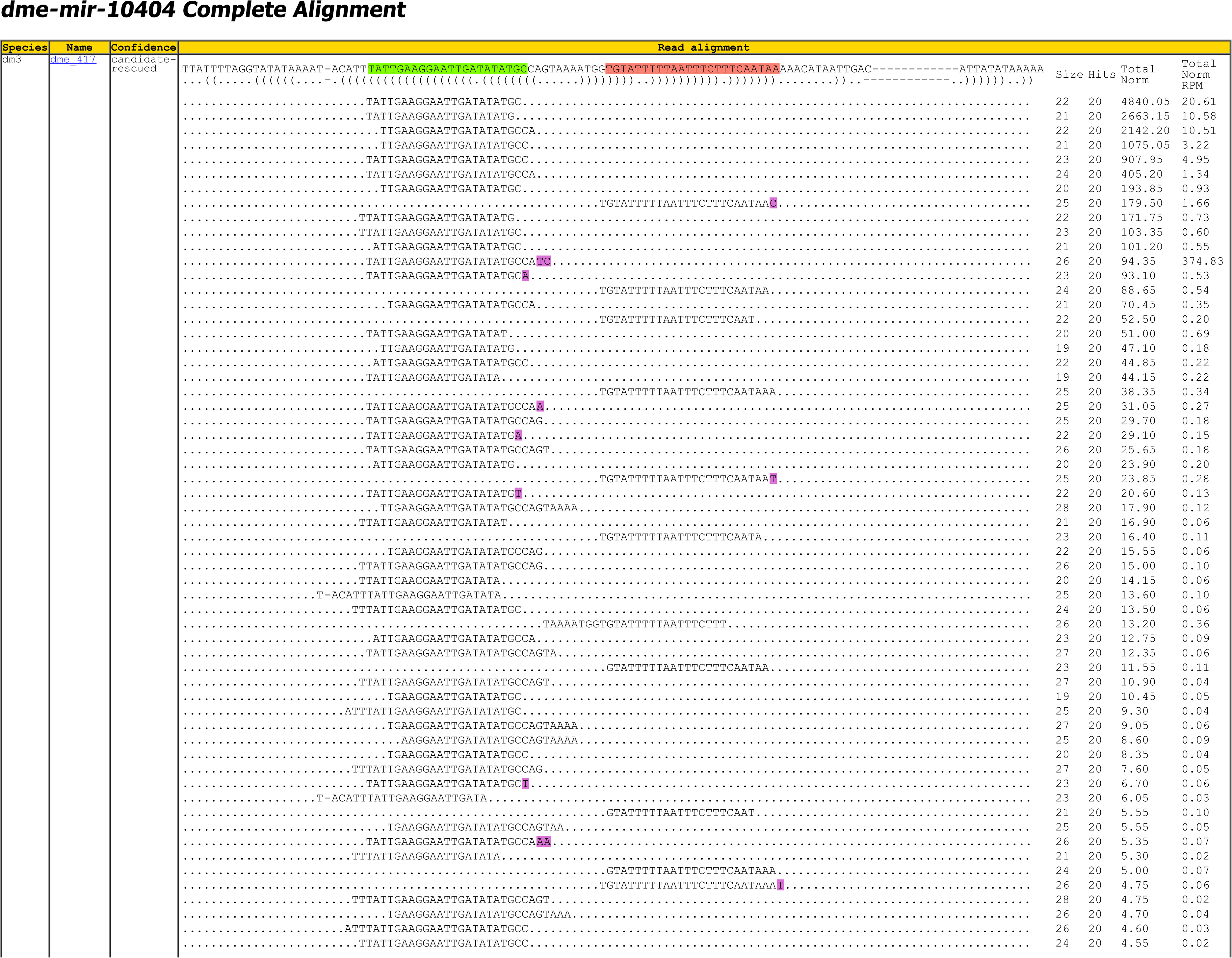
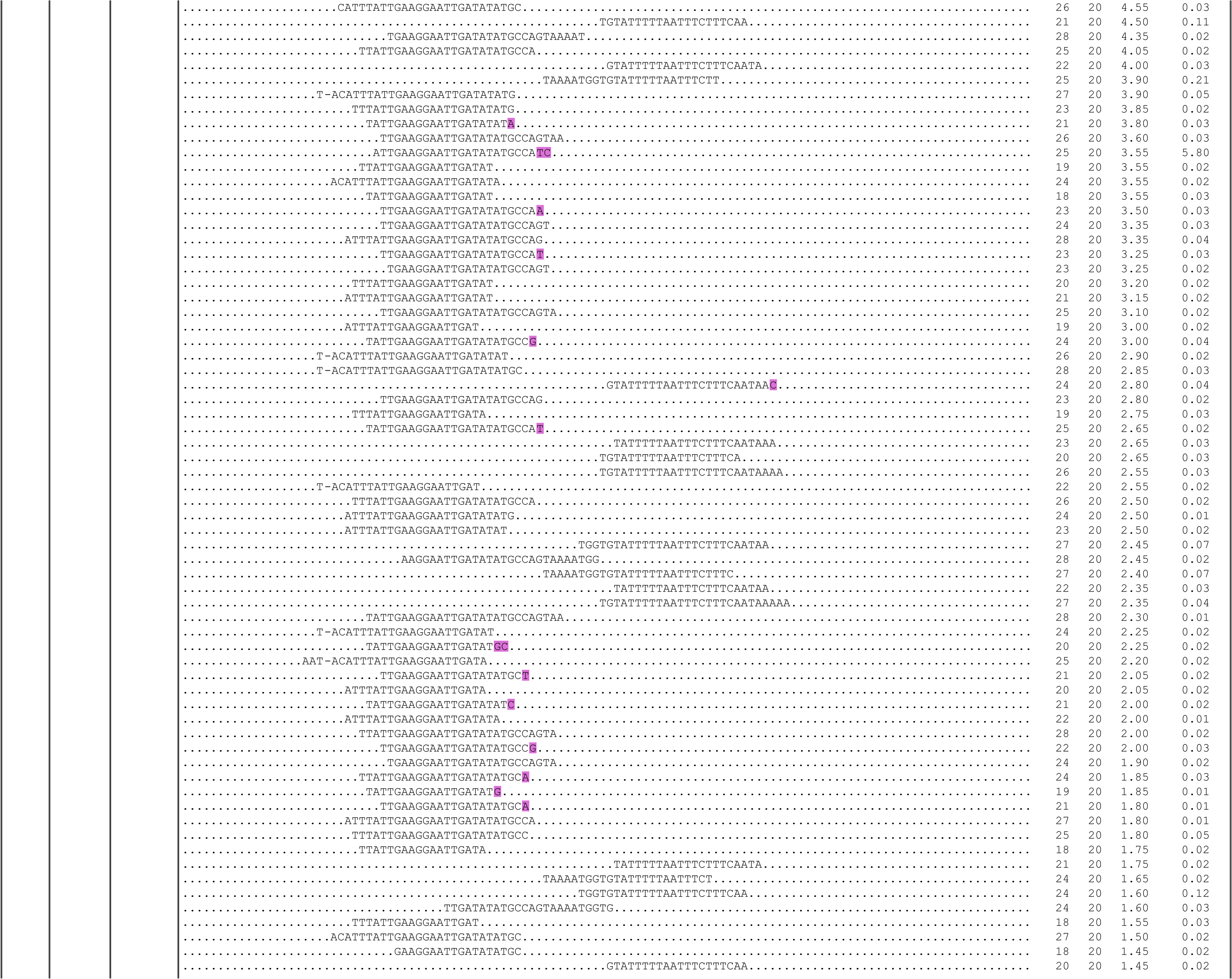
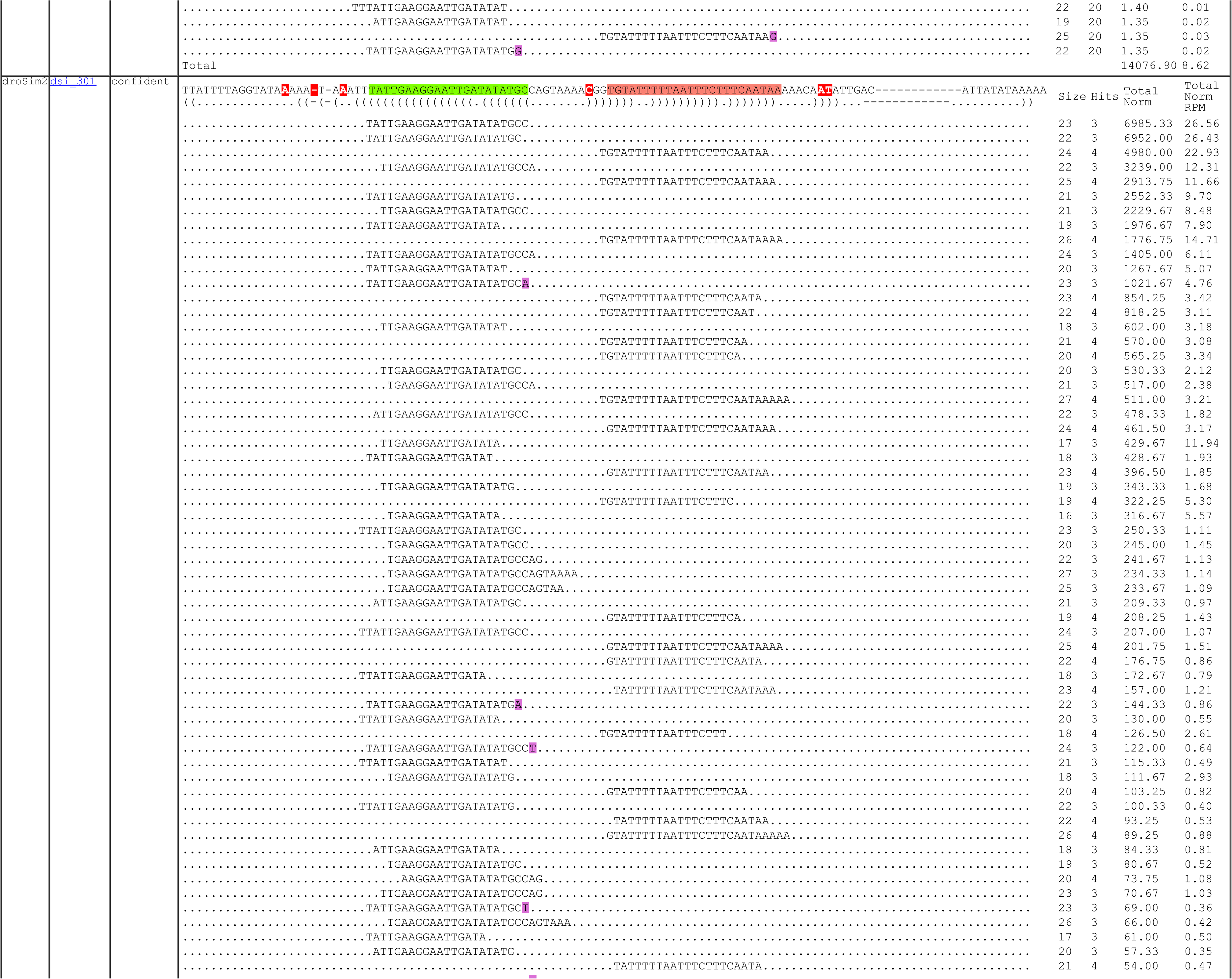
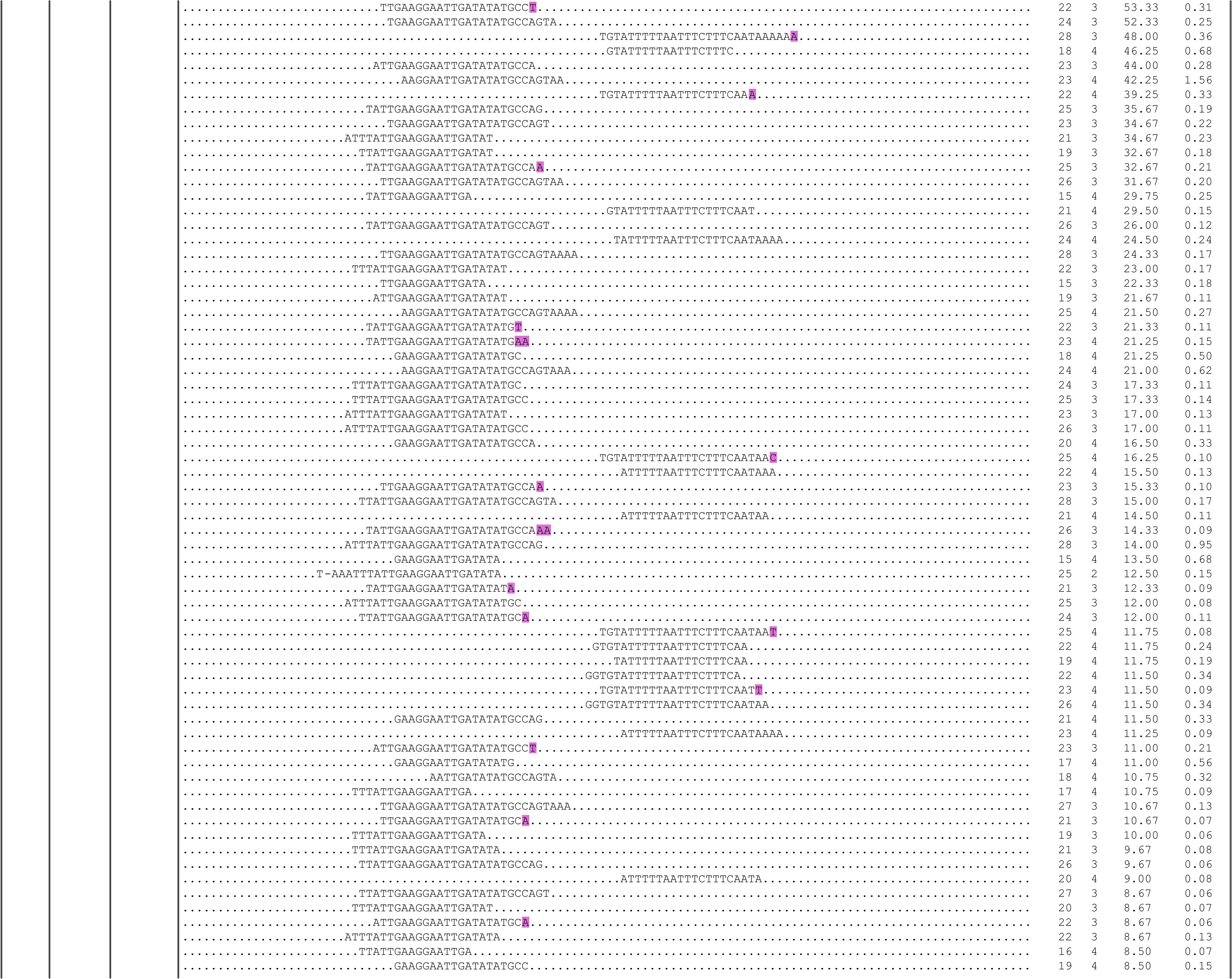
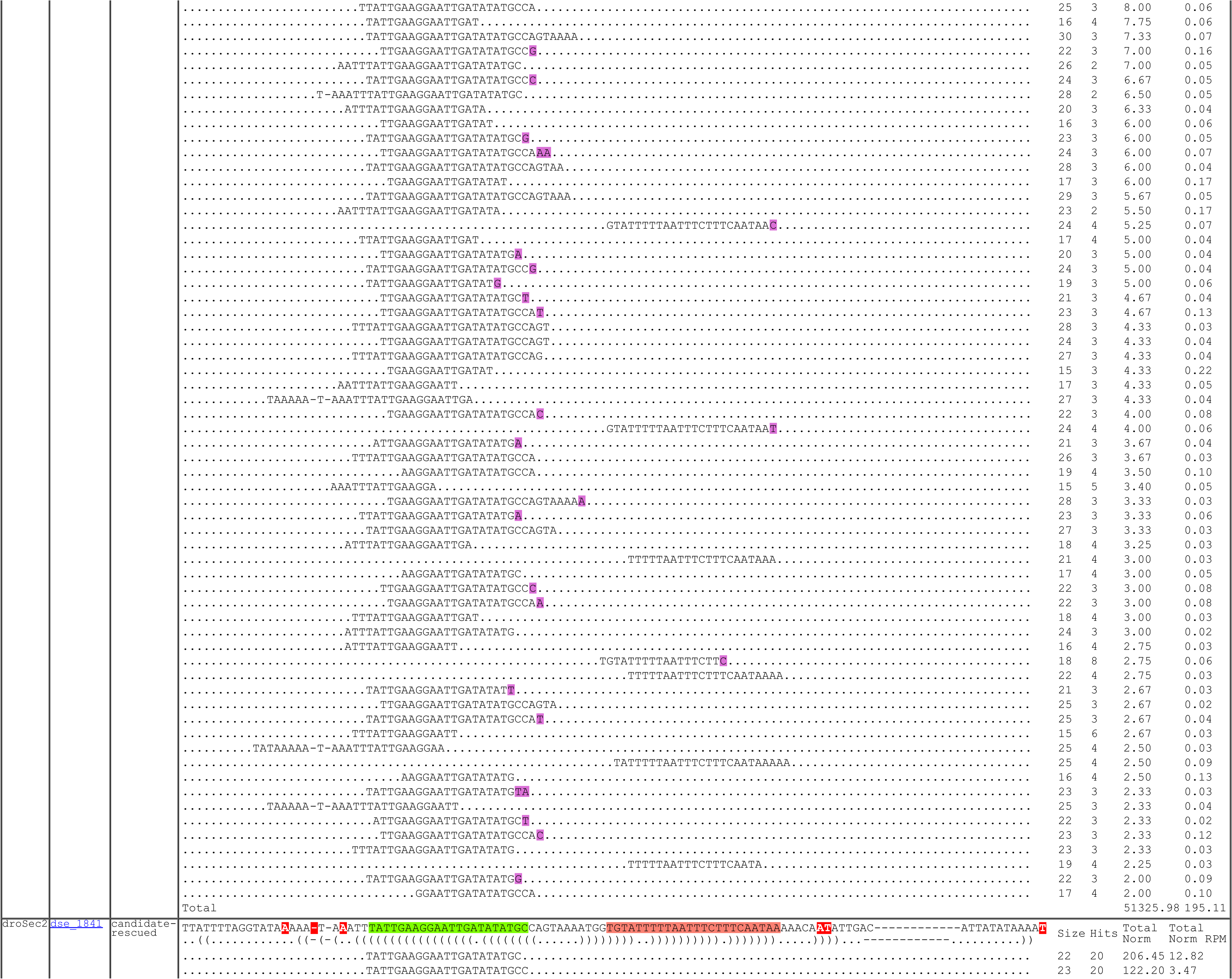
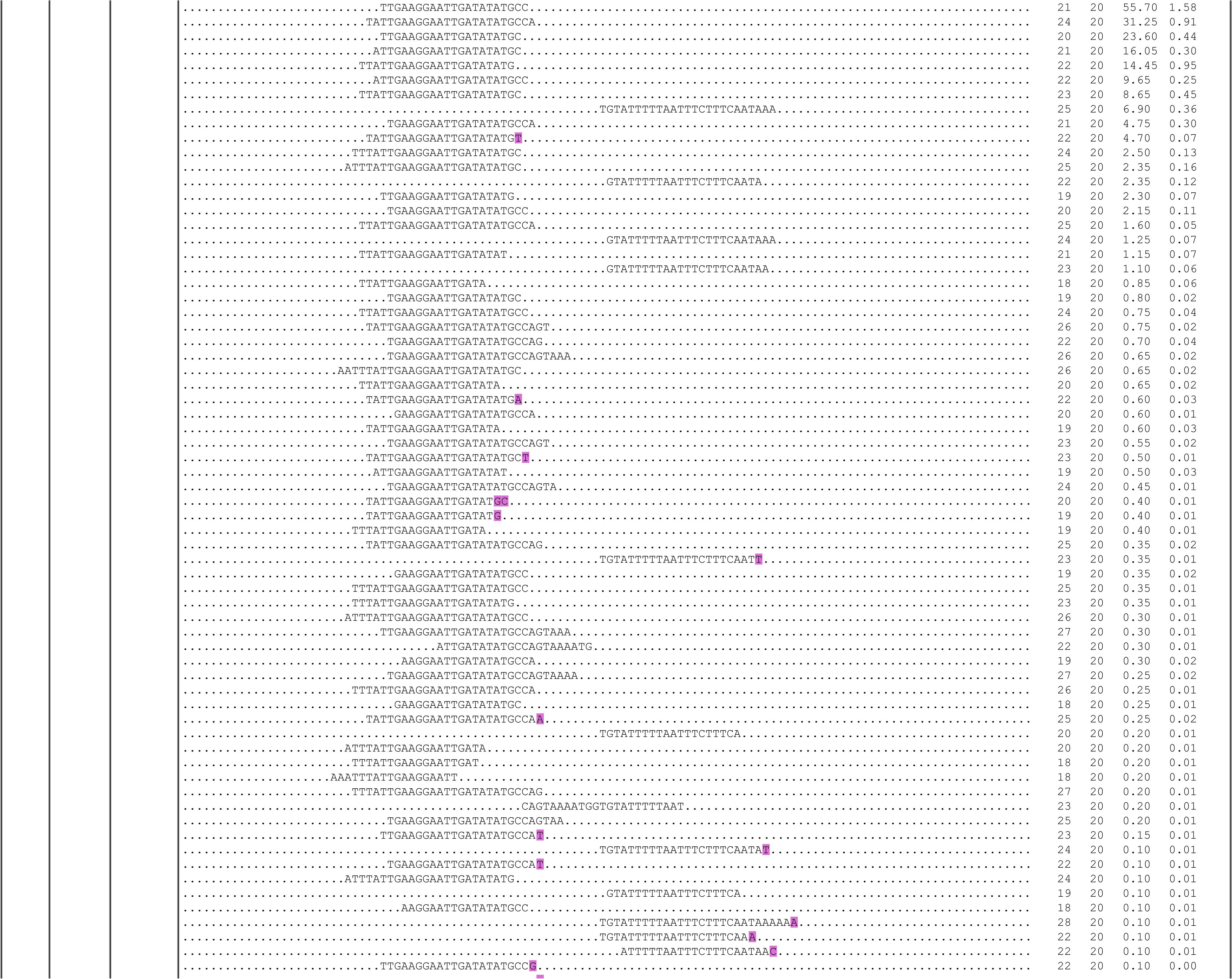
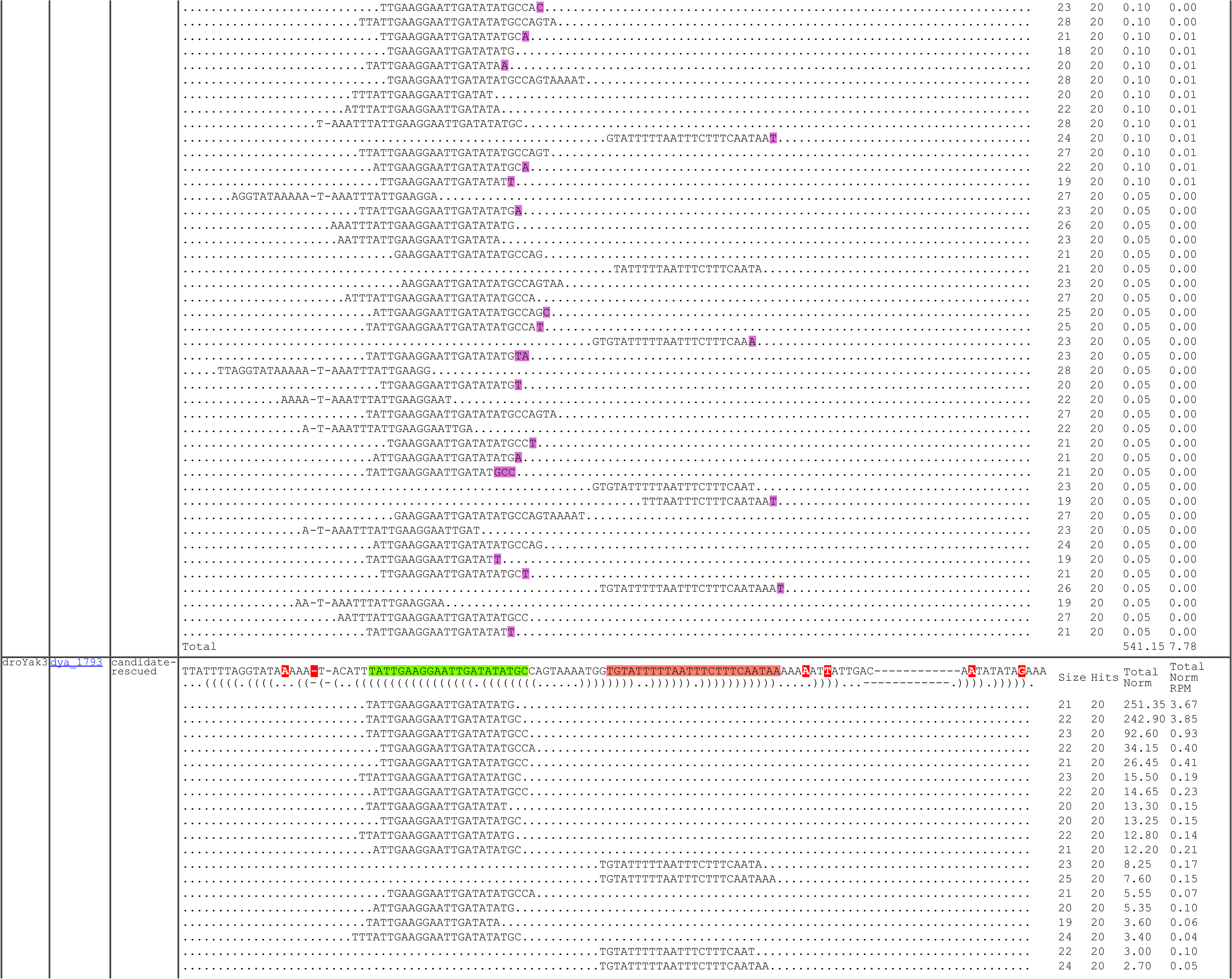
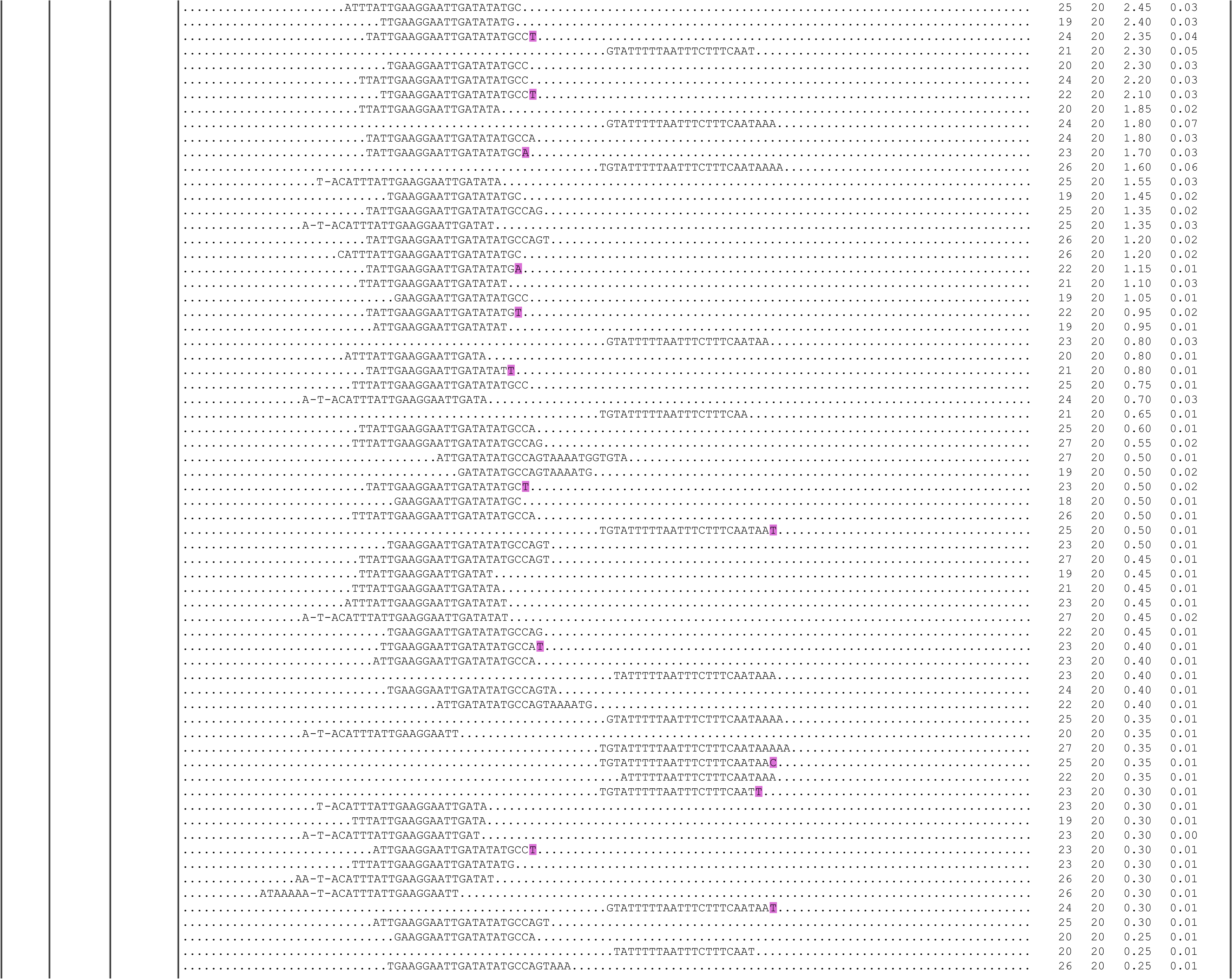
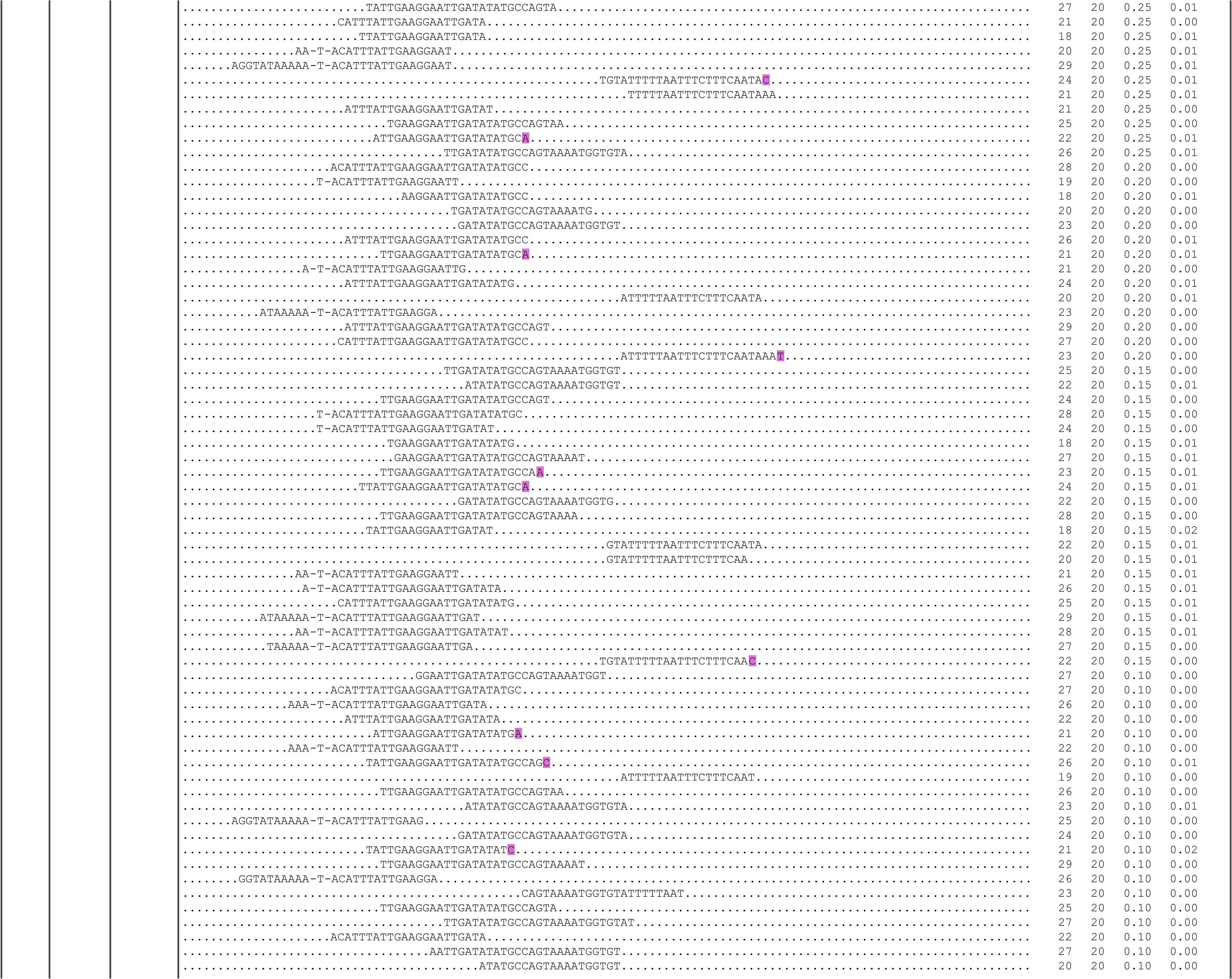
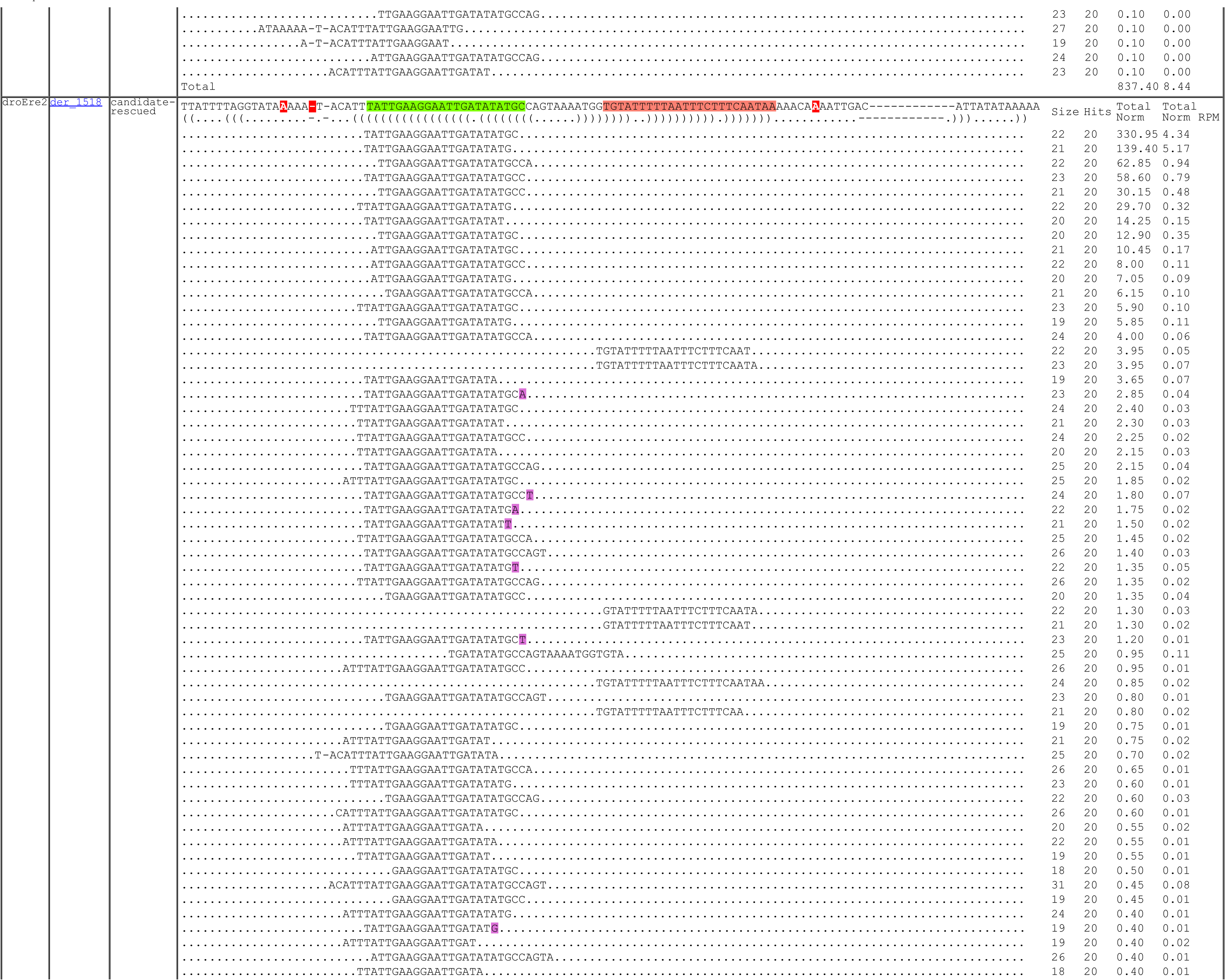
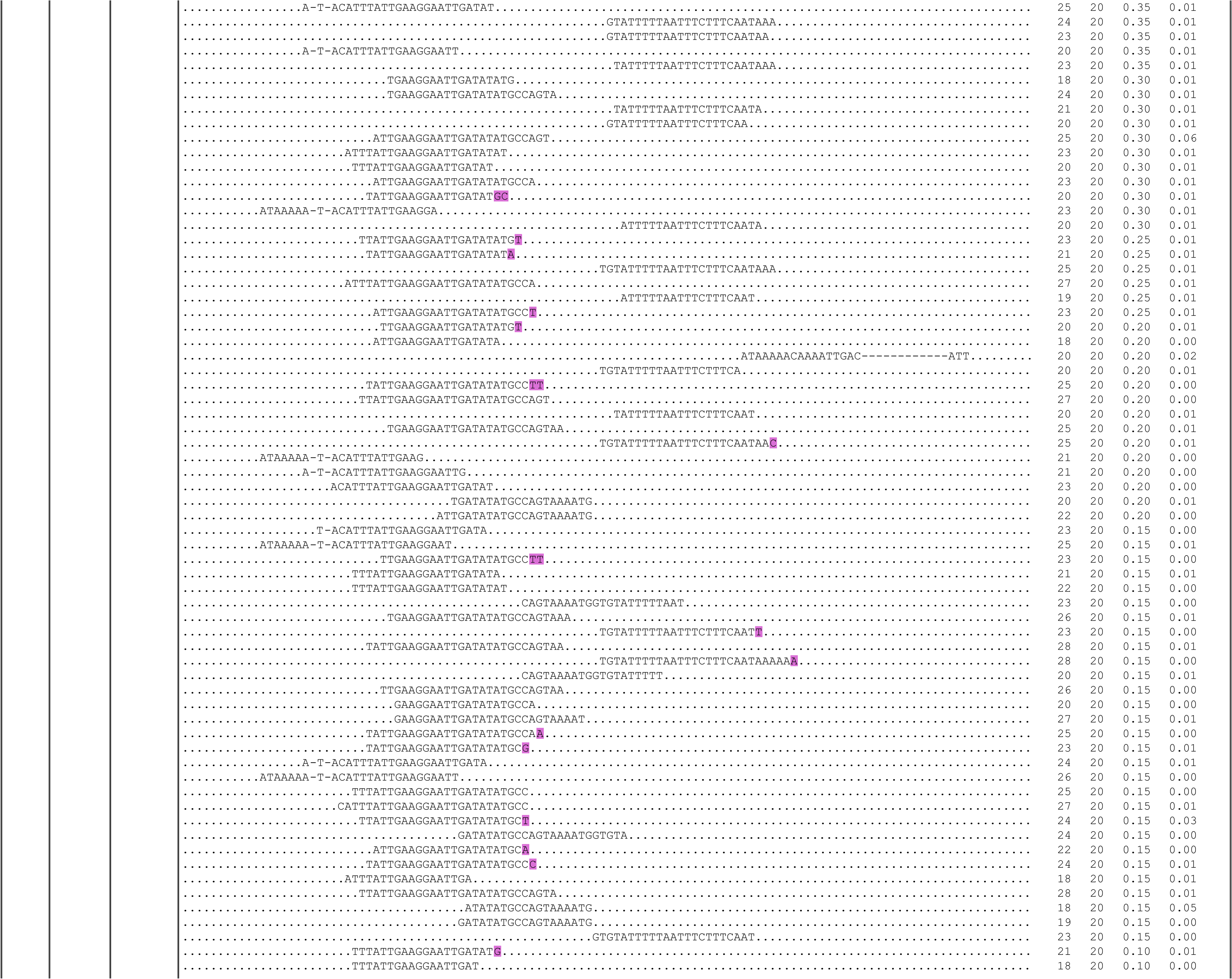
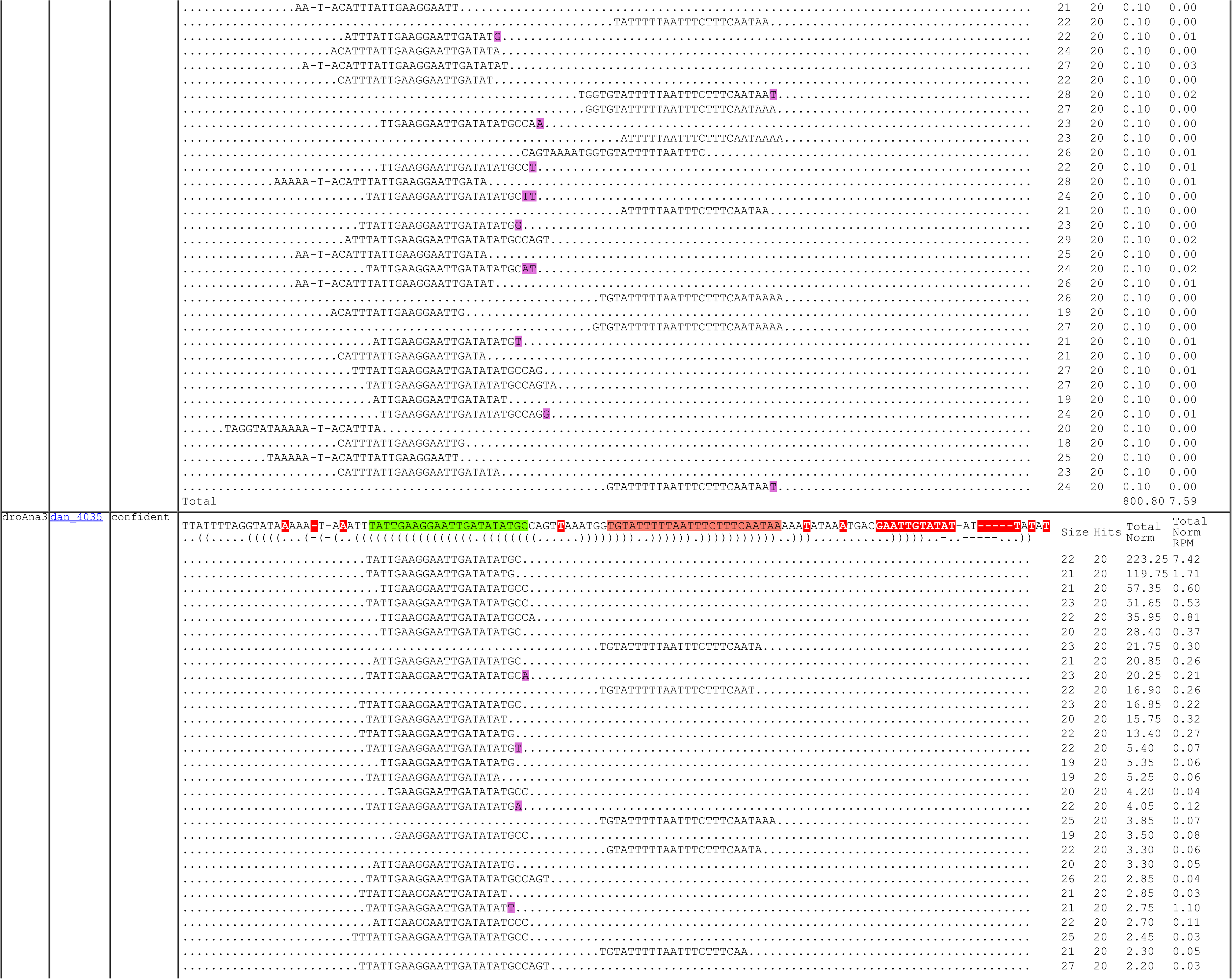
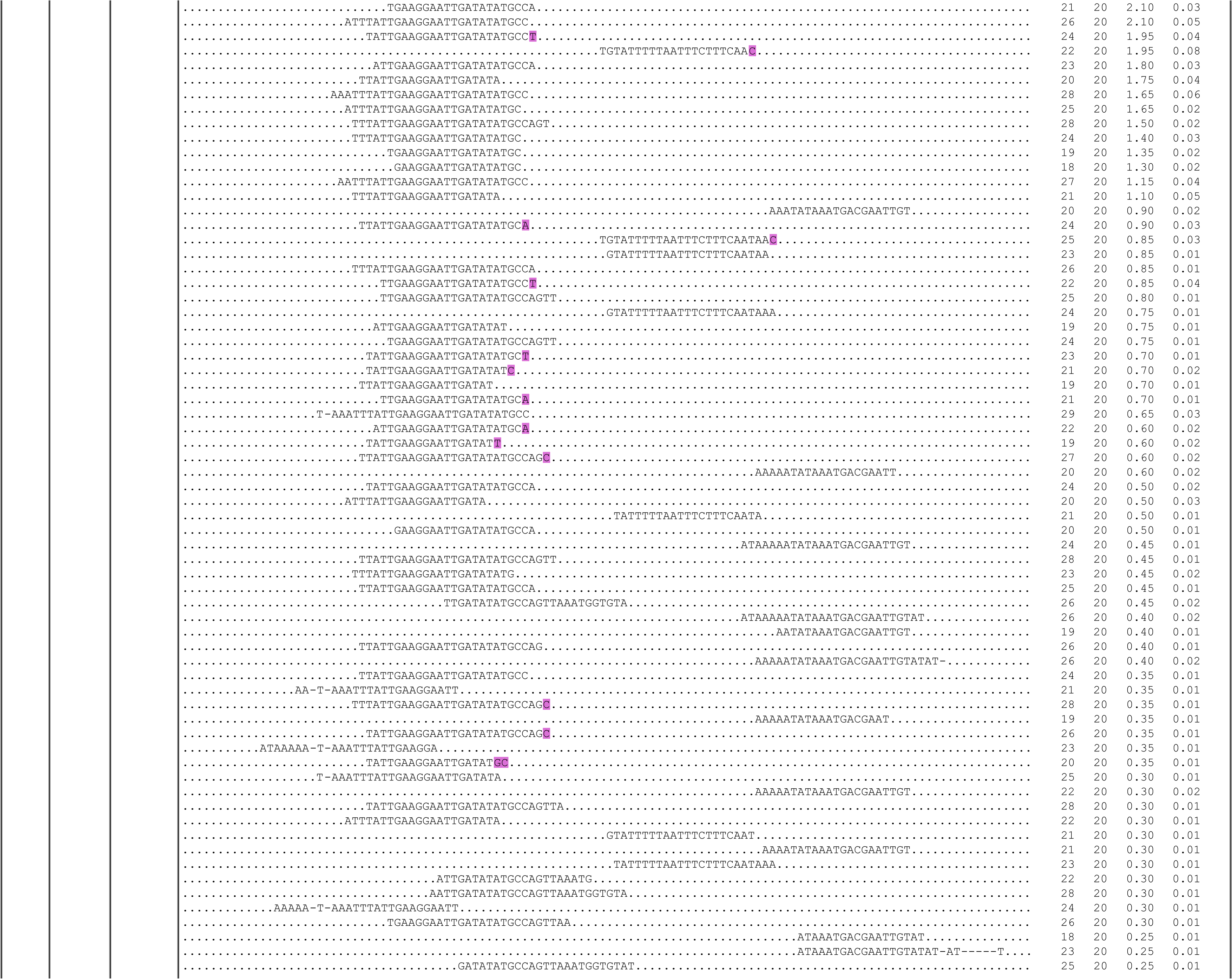
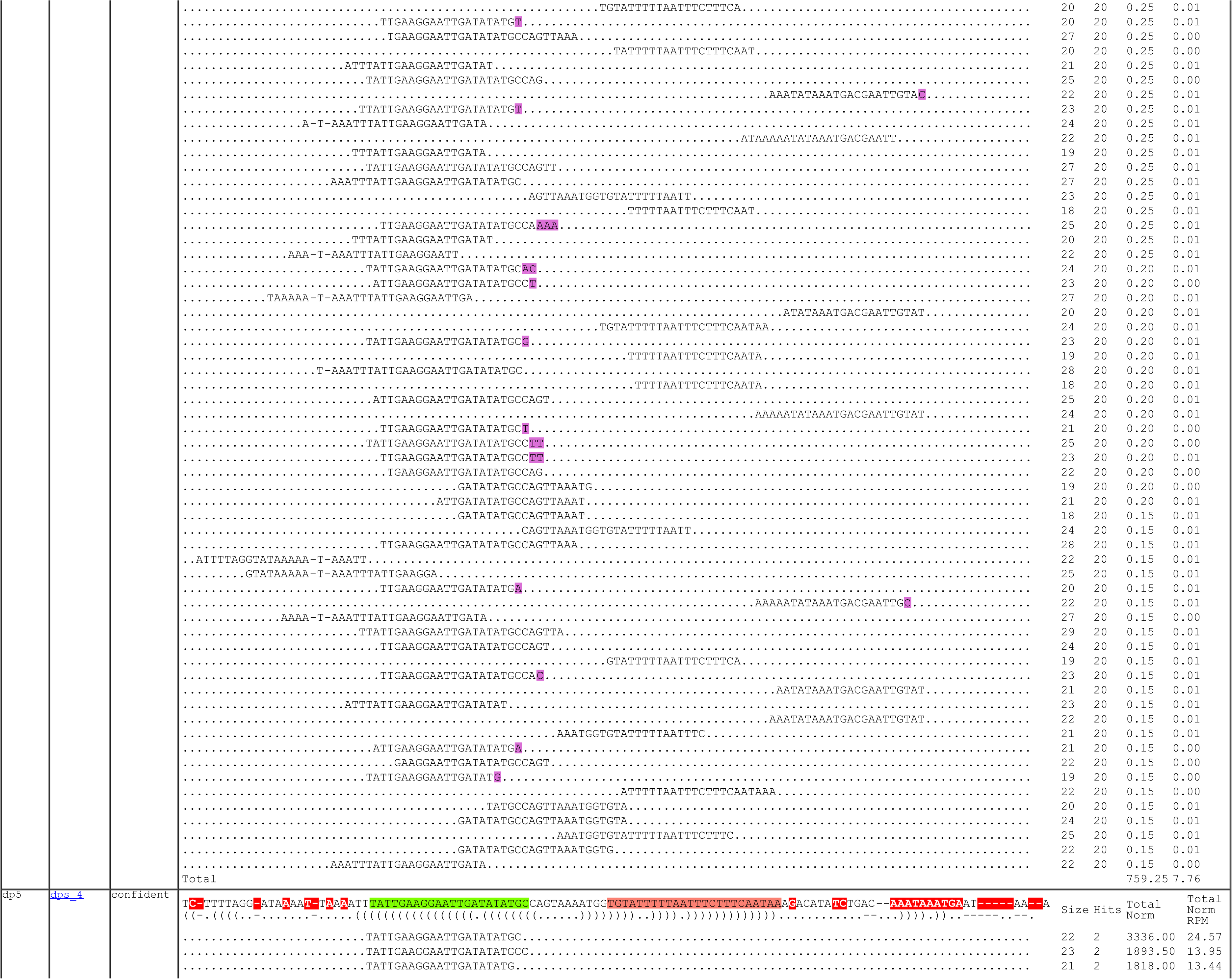
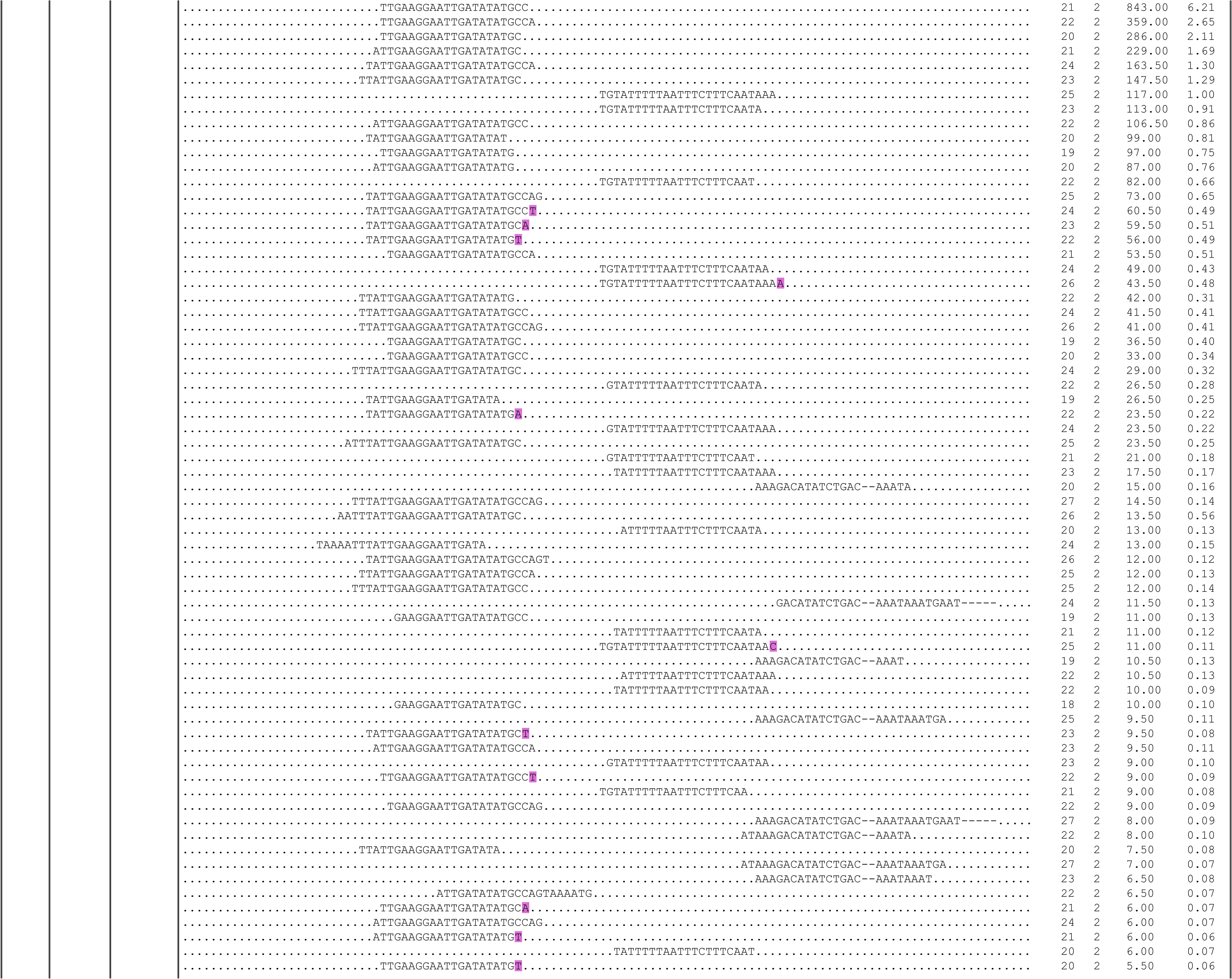
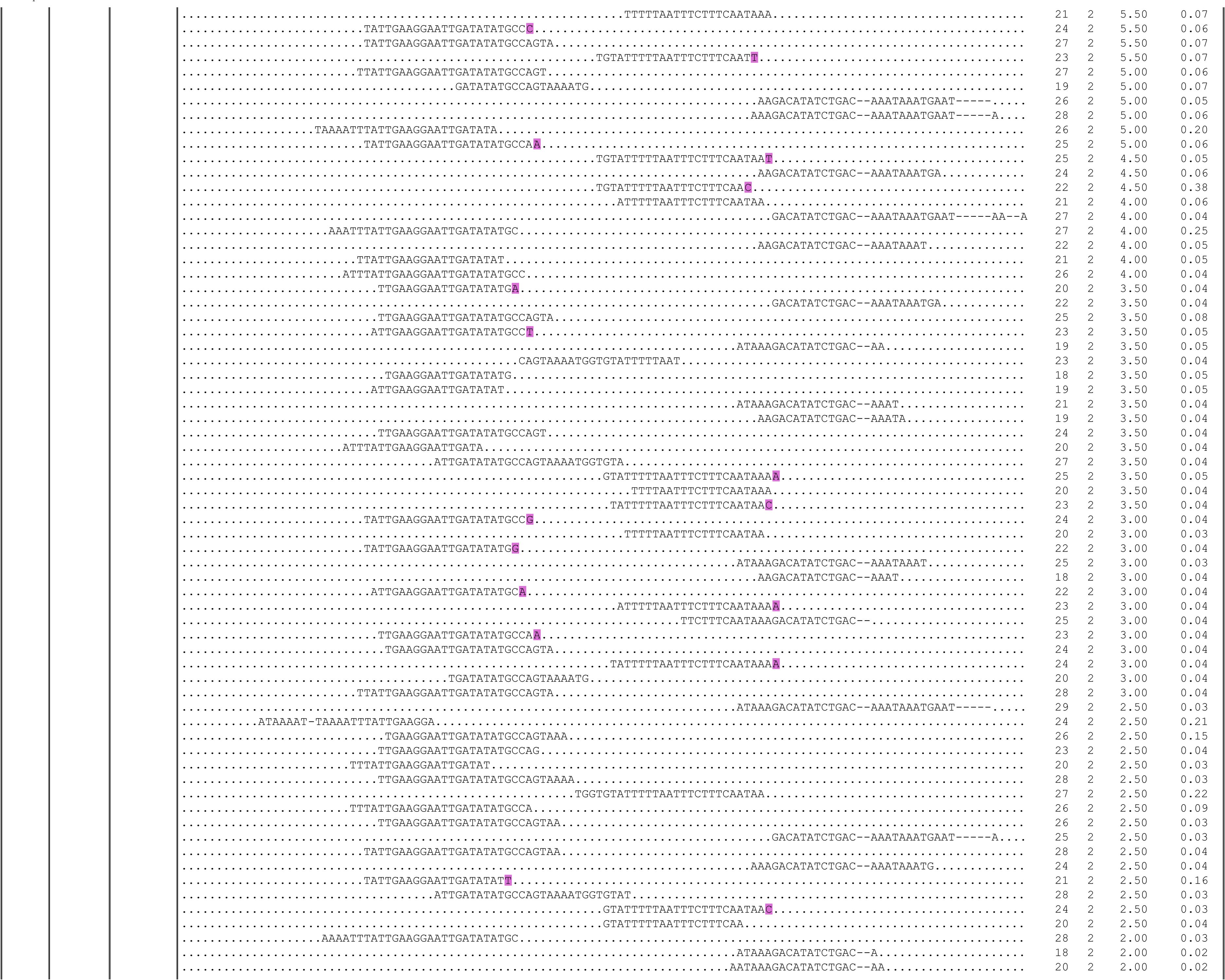
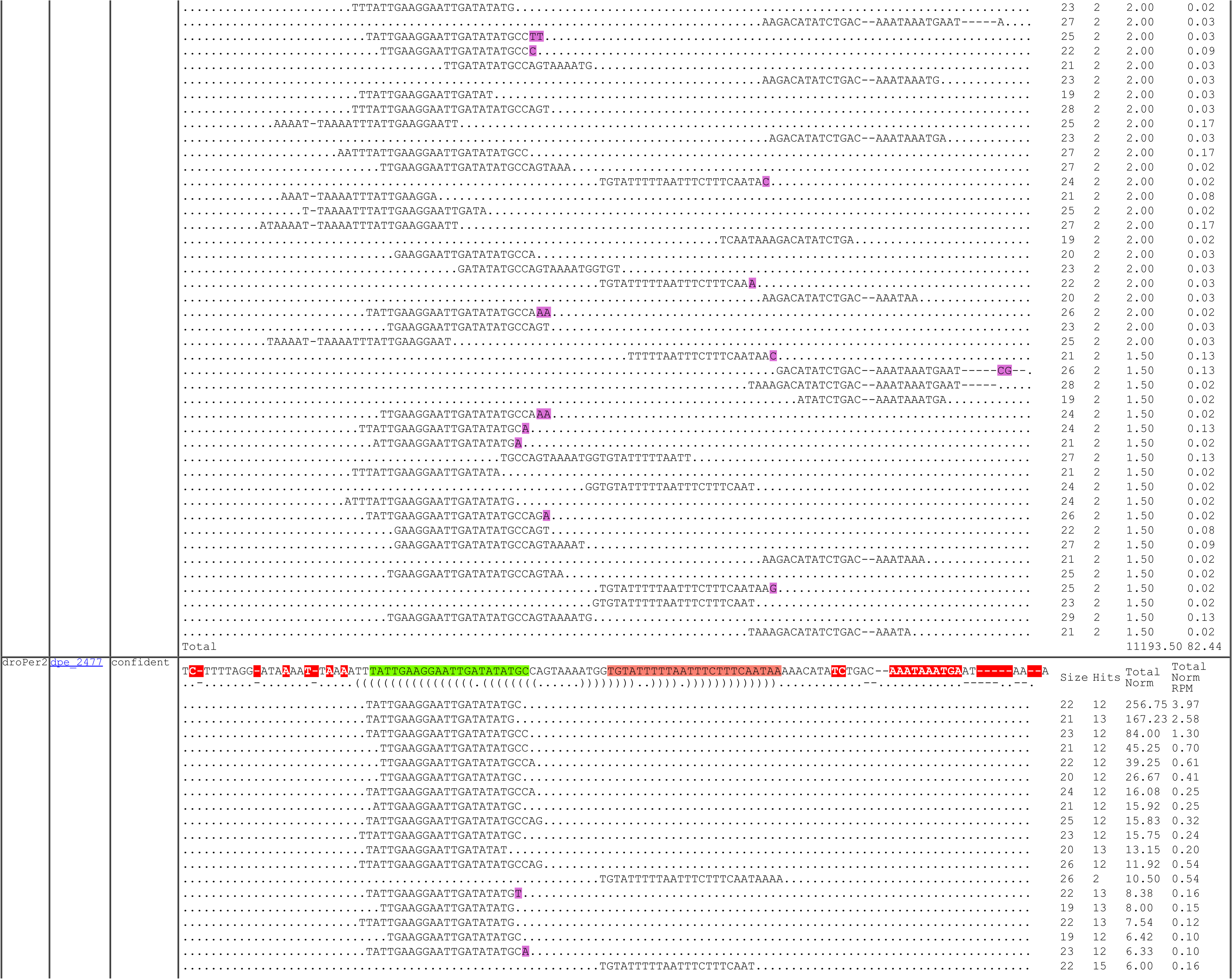
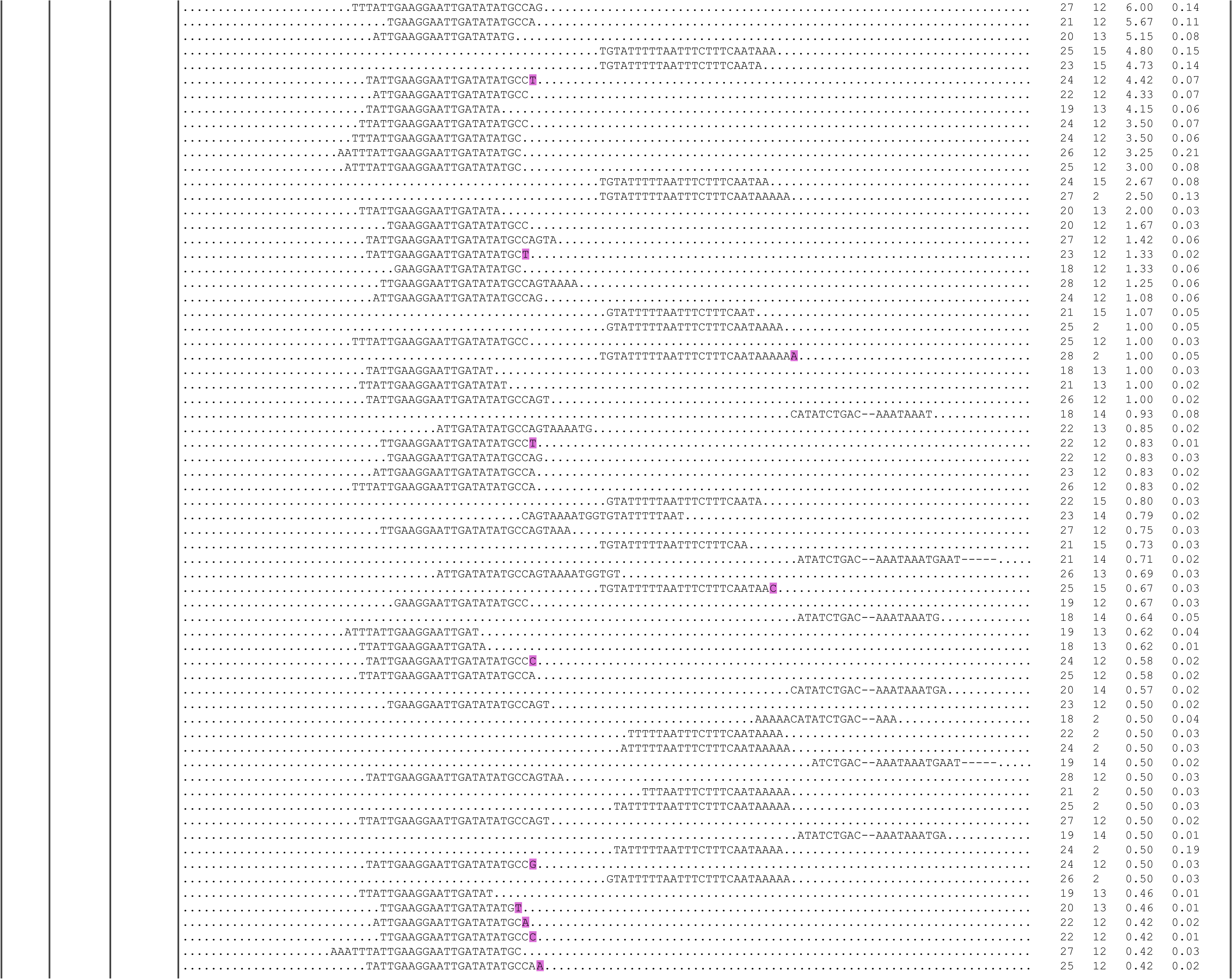
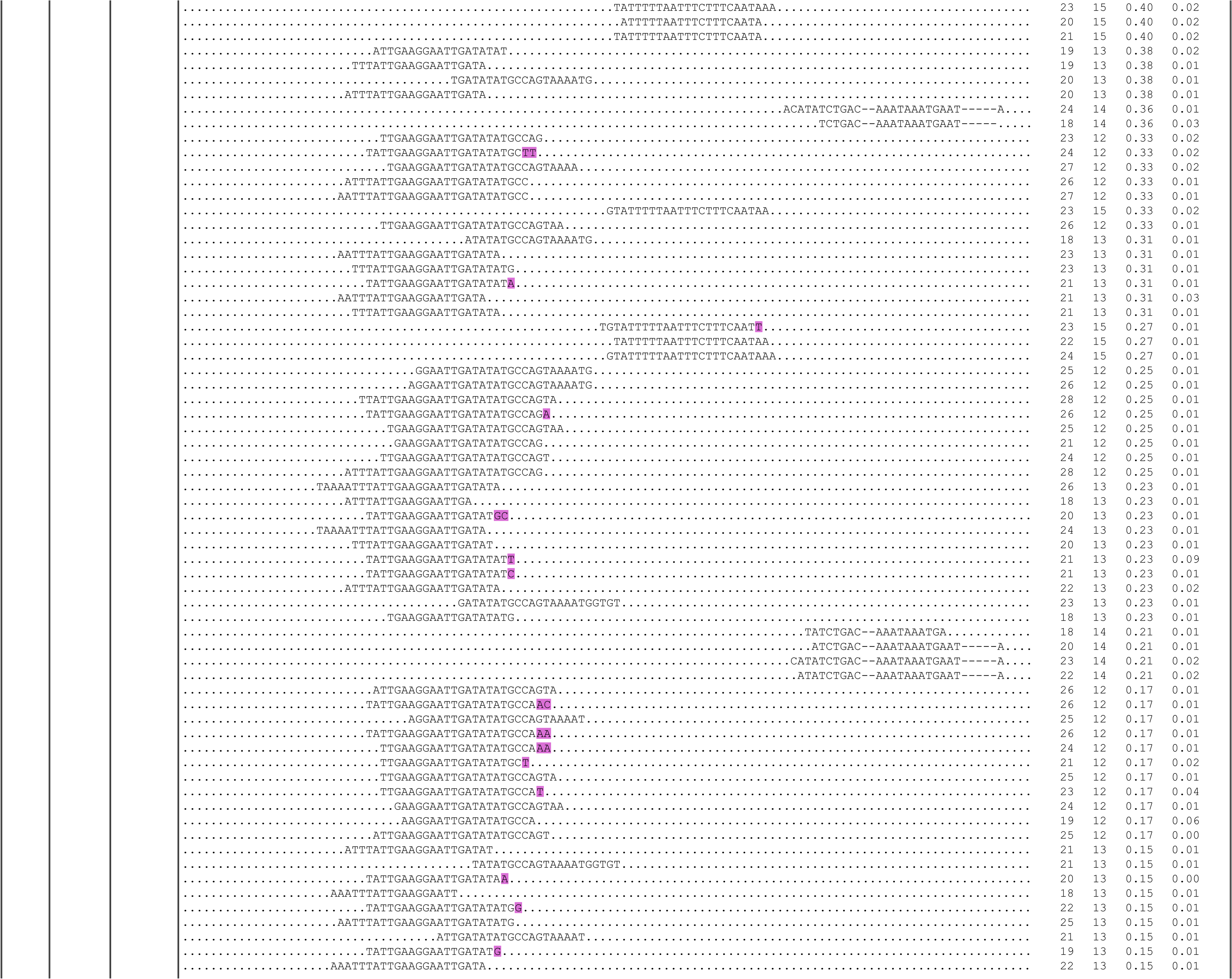
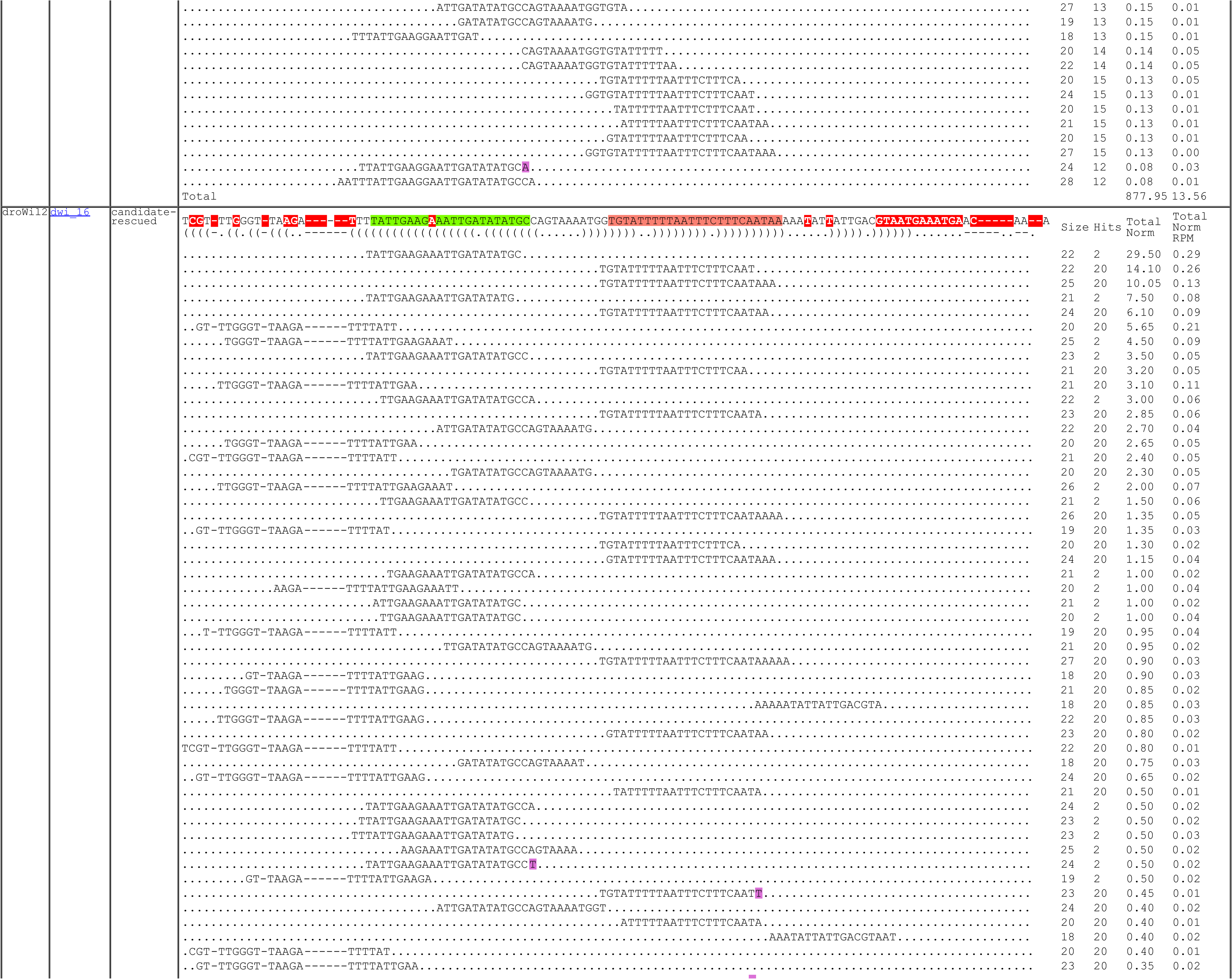
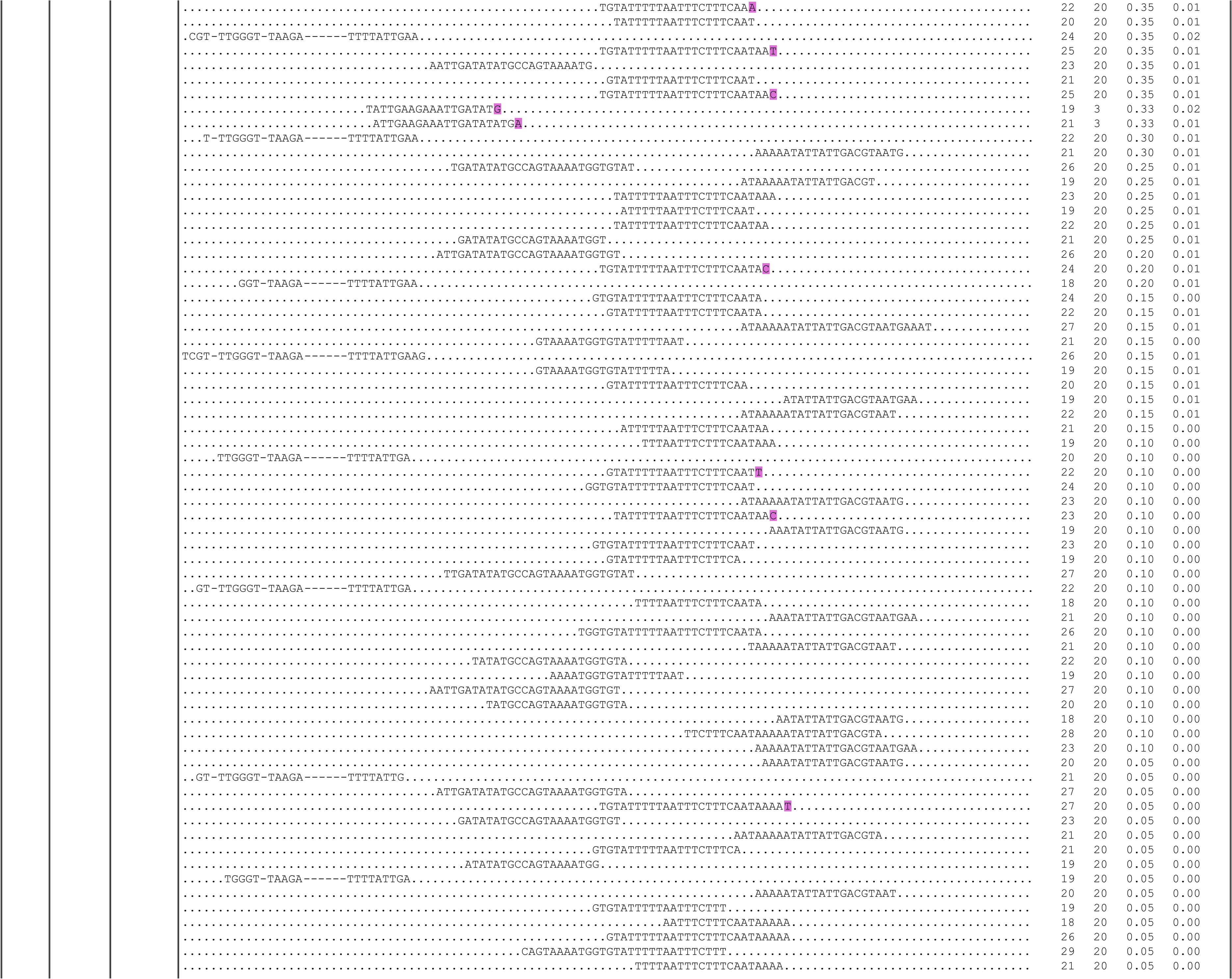
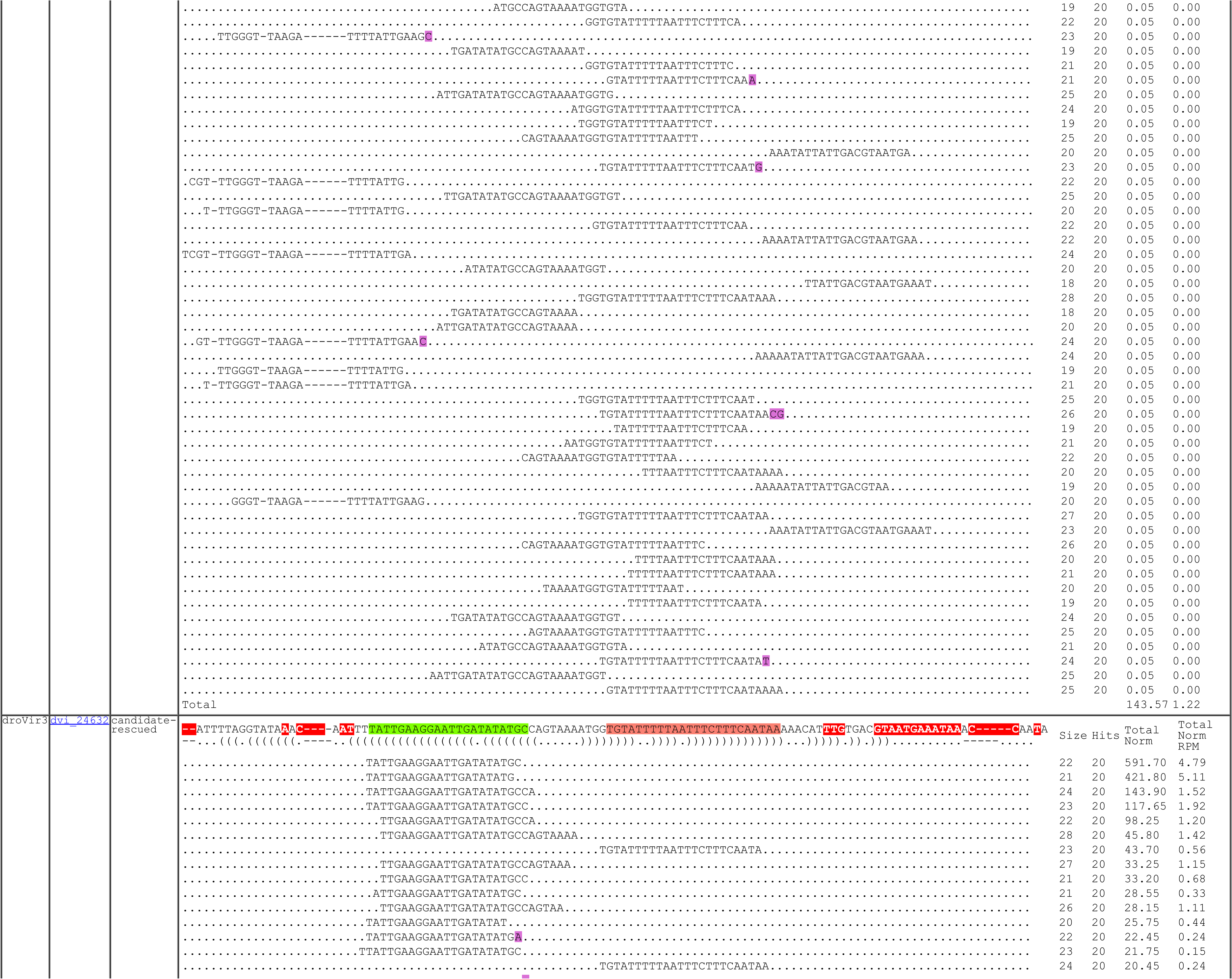
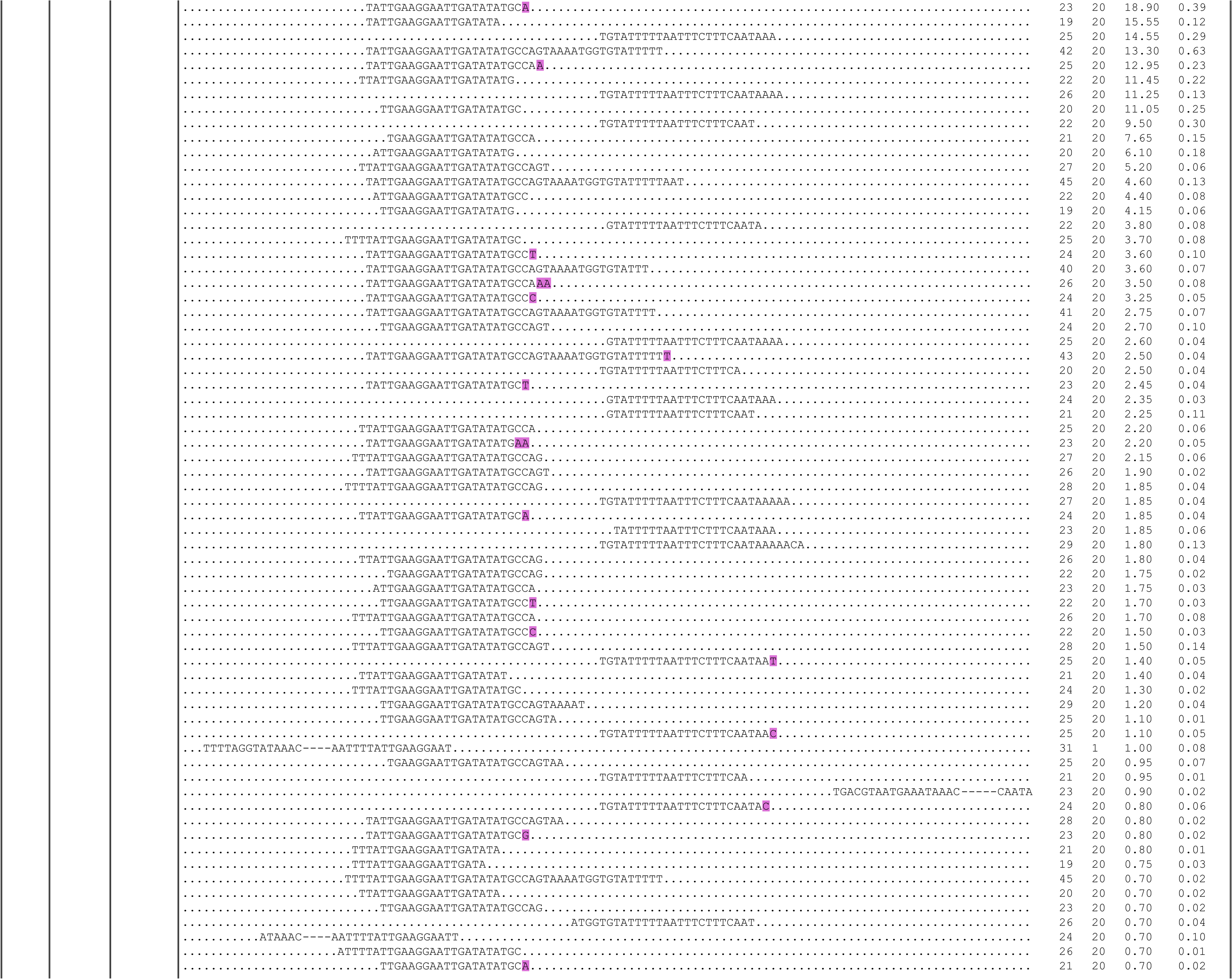
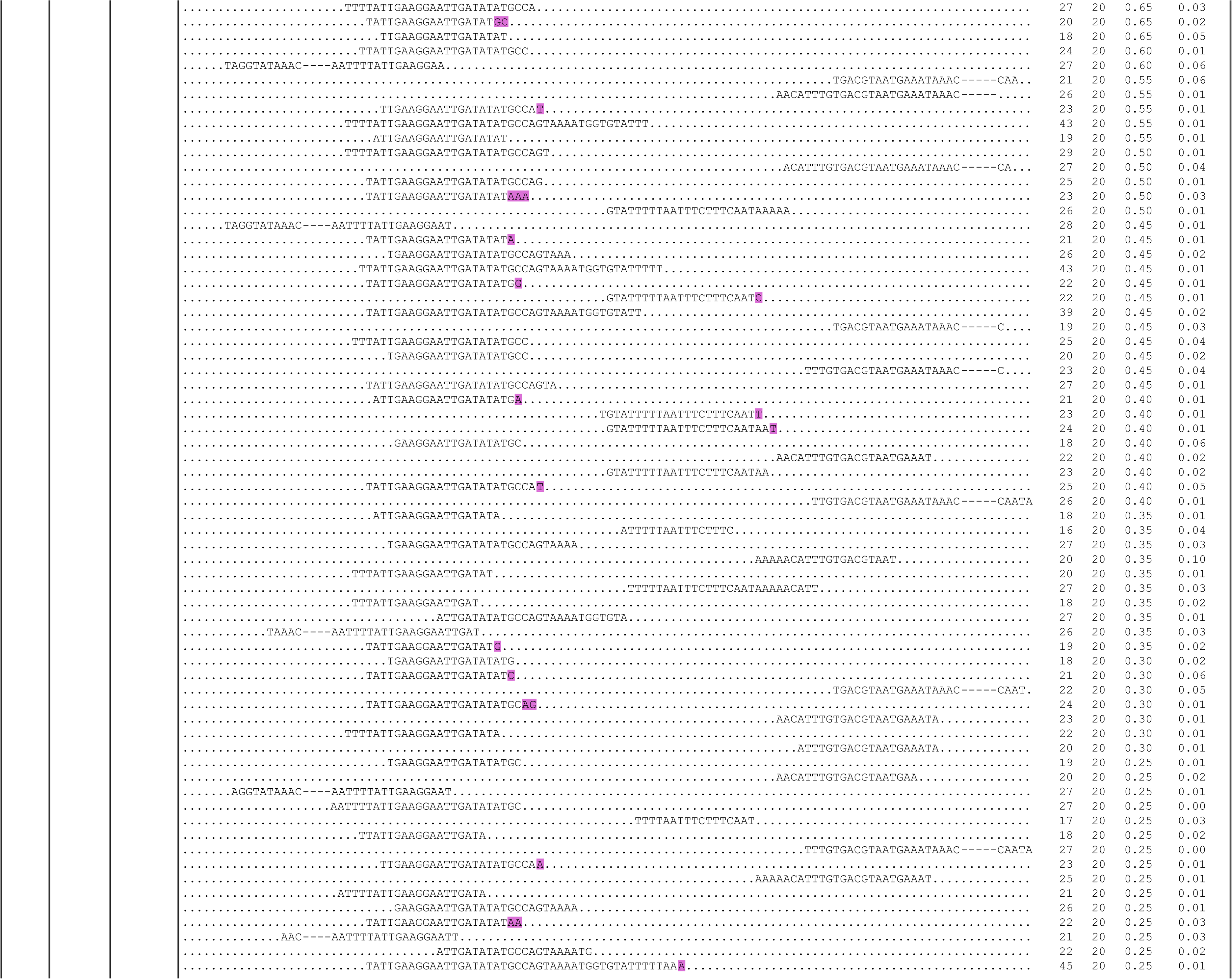
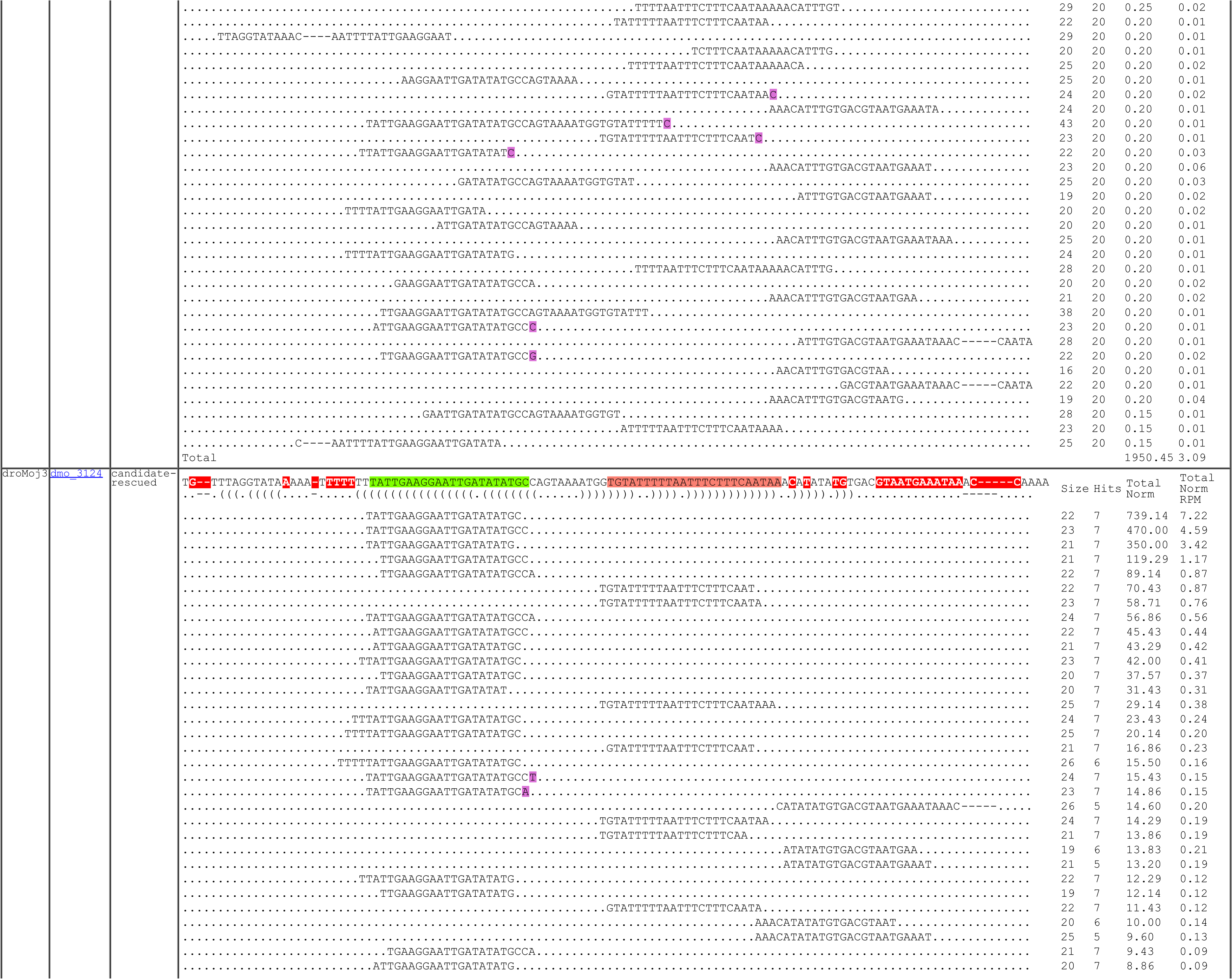
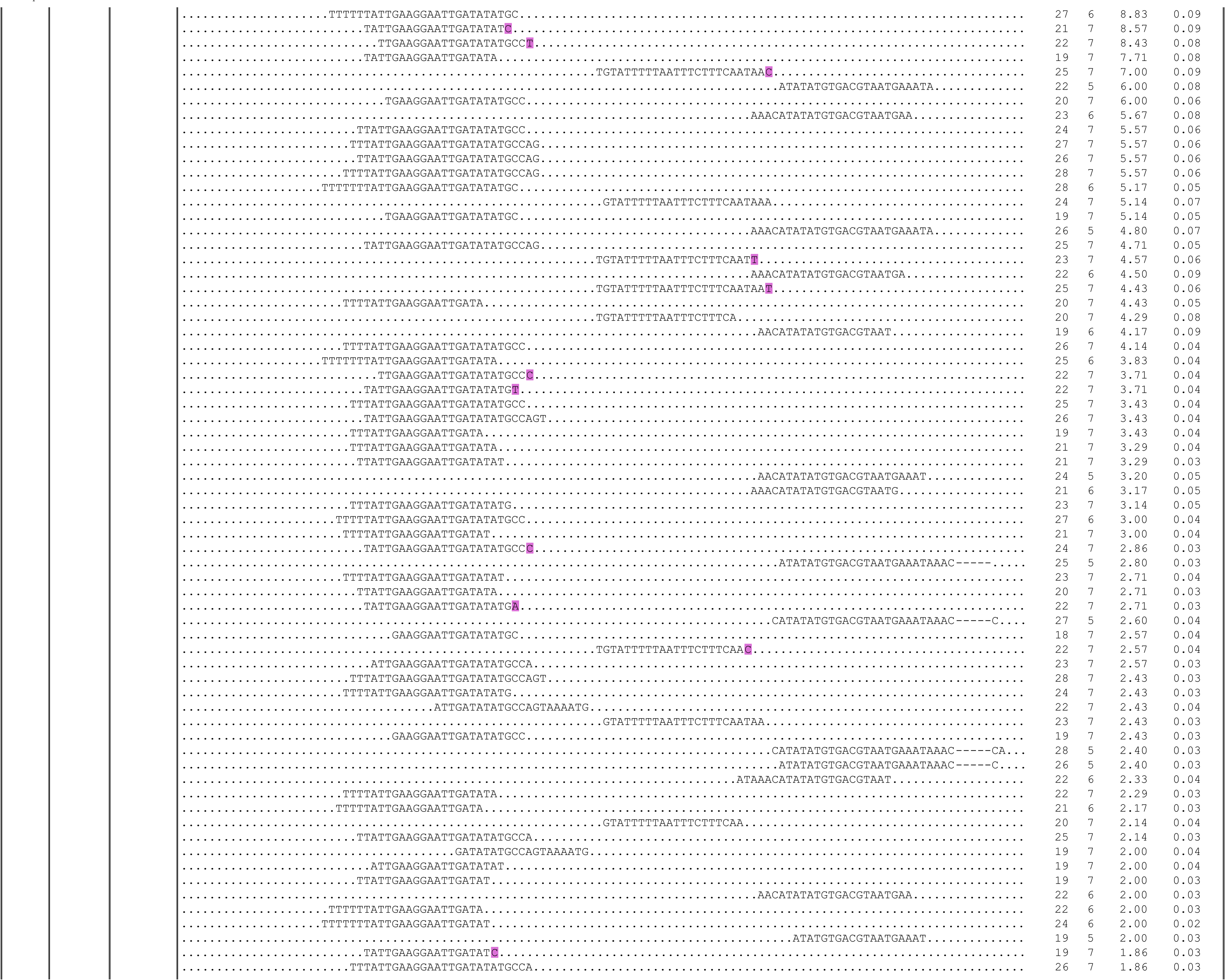
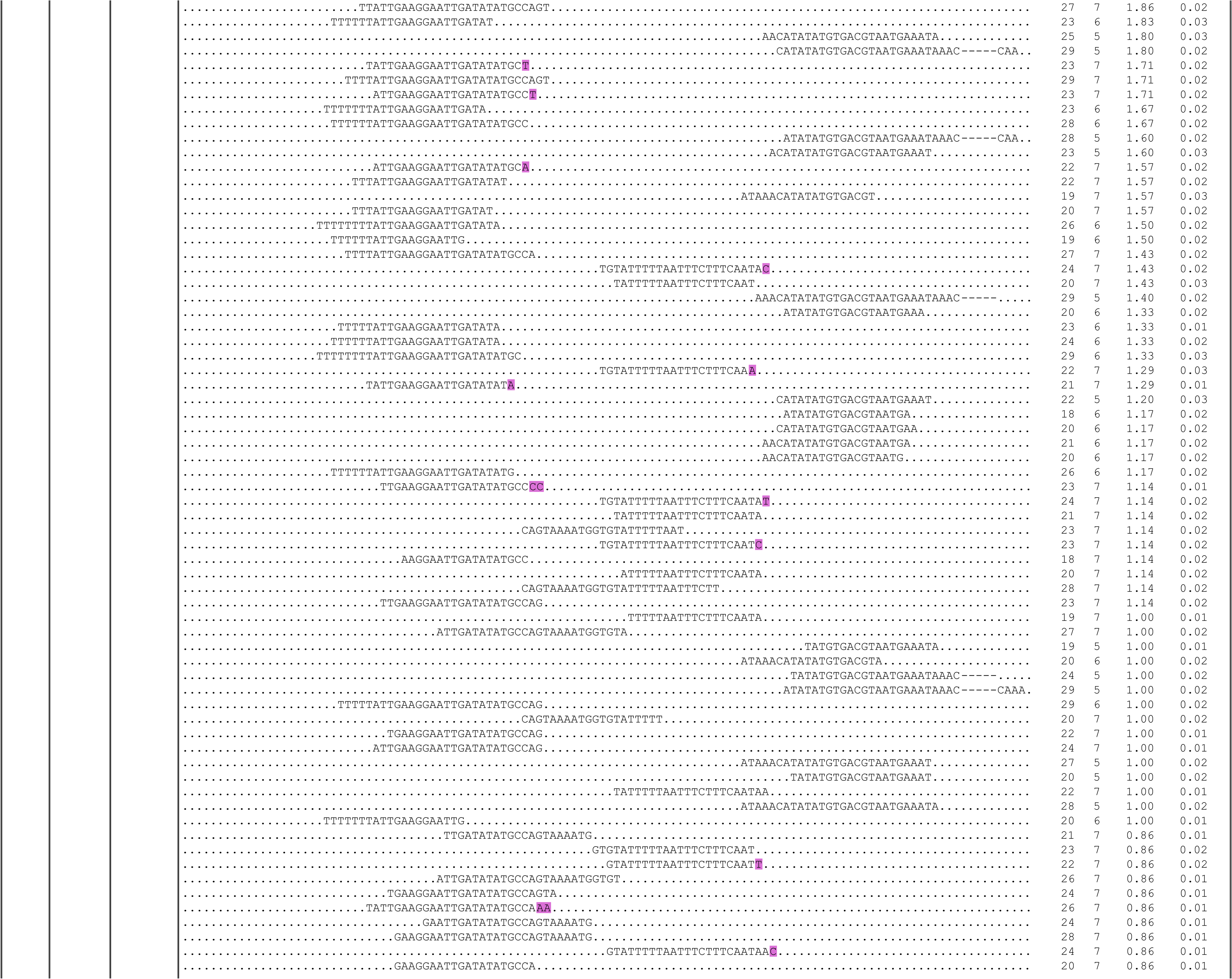
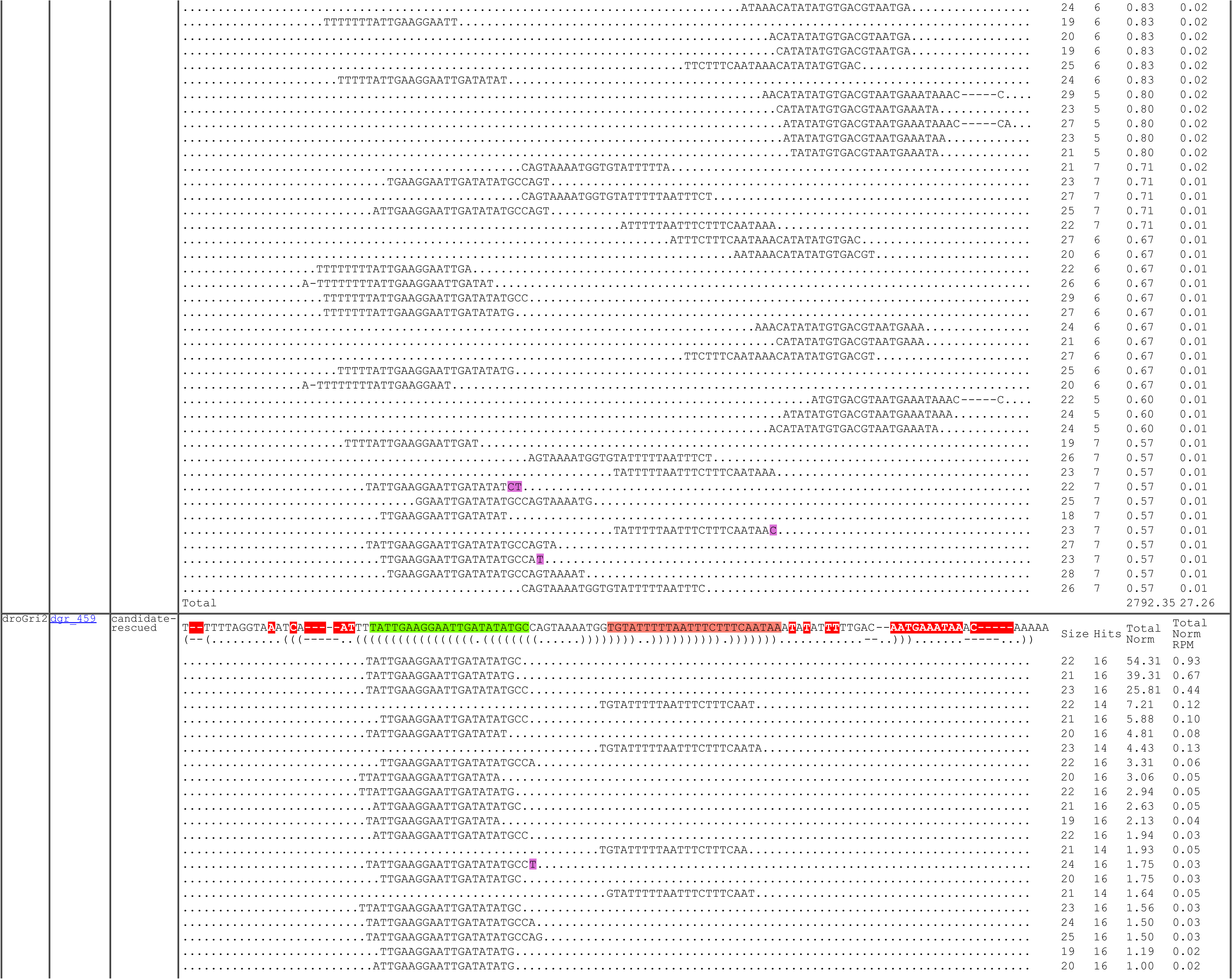
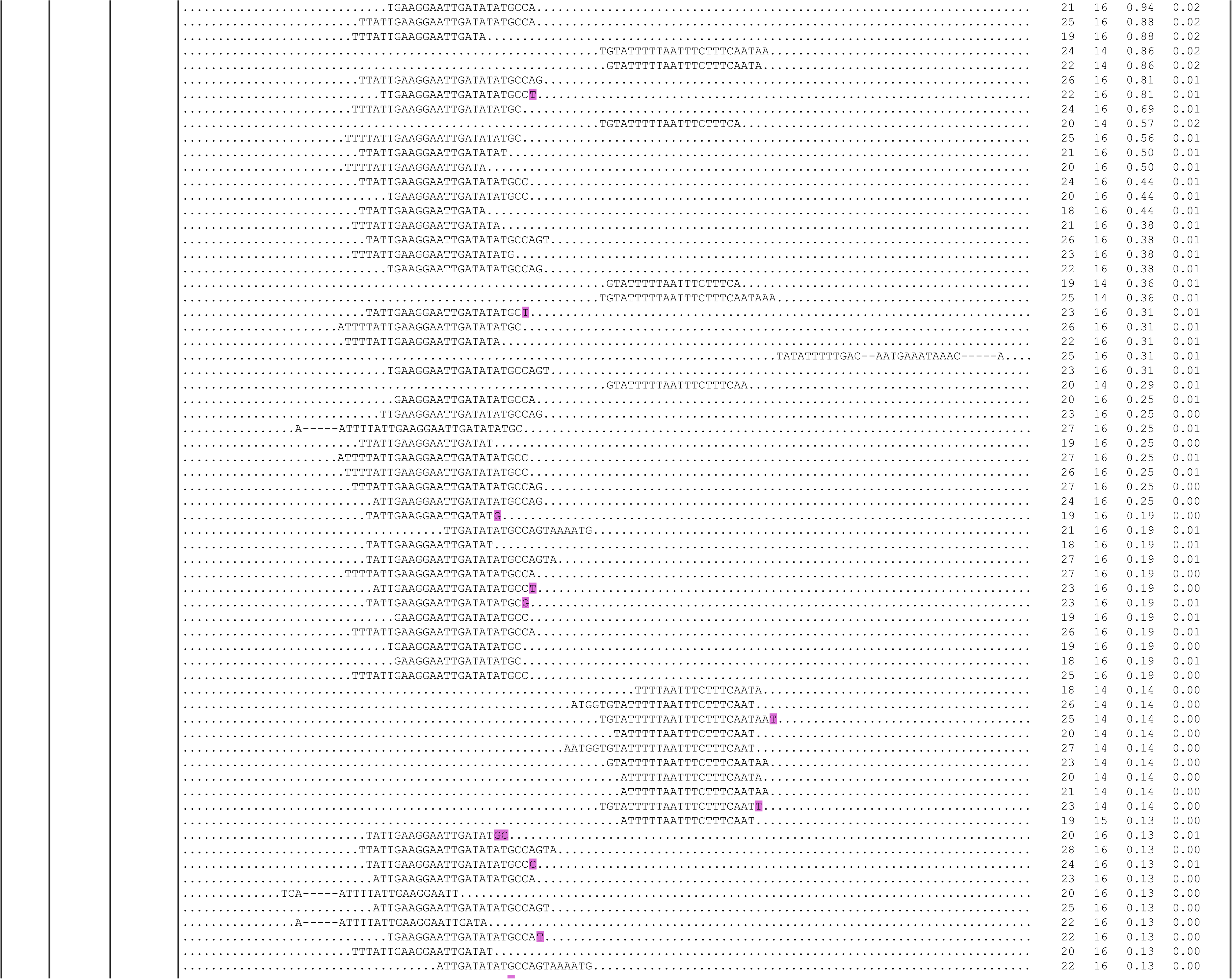
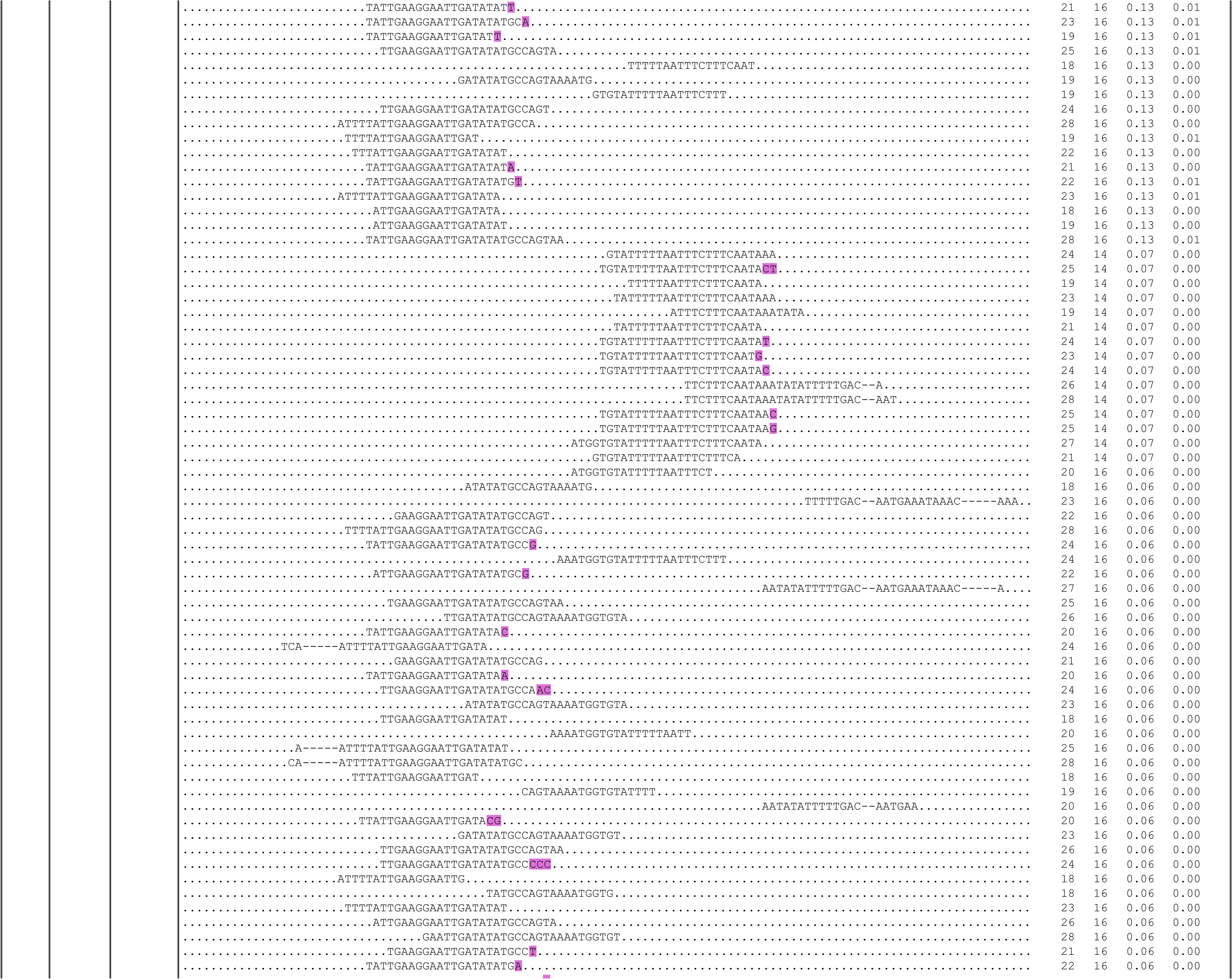
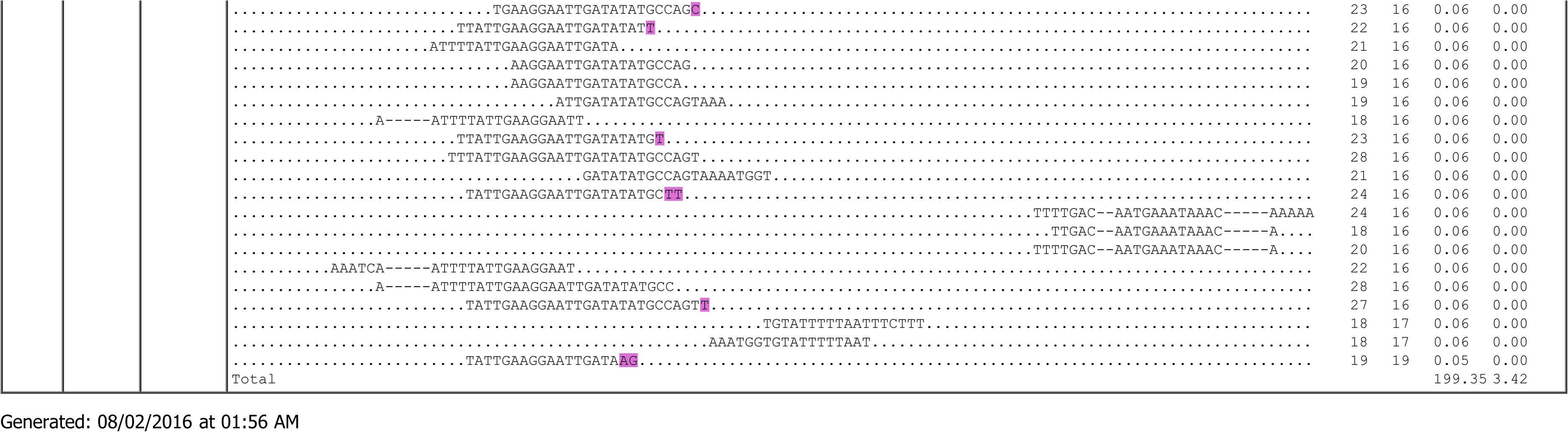
Read alignments for mir-10404, a conserved non-canonical miRNA generated from the ITS1 spacer in ribosomal RNA, across the Drosophilid phylogeny.

**Supplementary Figure S8:**
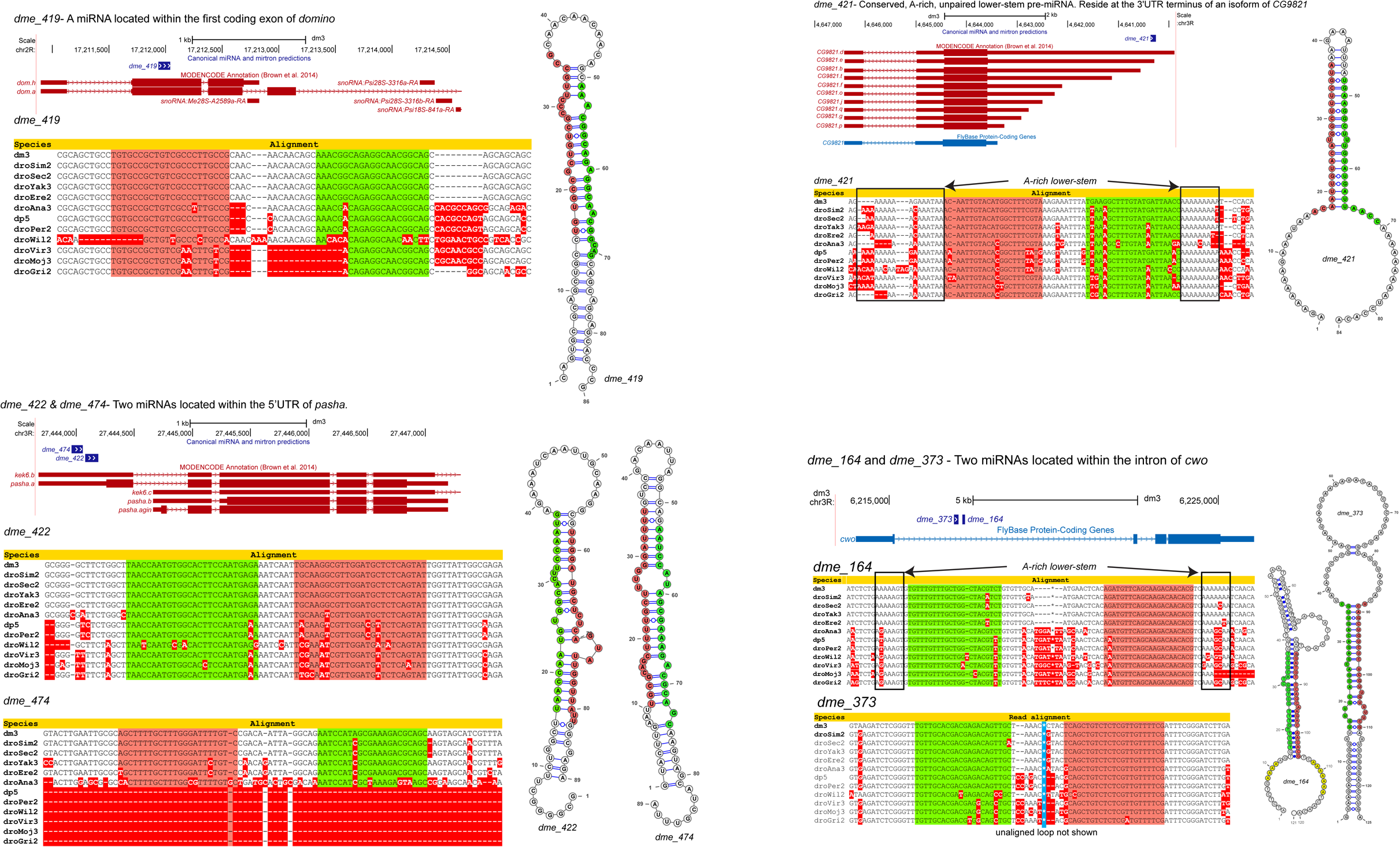
Other well-conserved miRNAs identified within this study. Alignment and representative hairpin structure for each conserved miRNA. Included is dme_474 which is not well-conserved but is one of the Drosha-cleaved hairpins within the pasha 5’ UTR. Note that unlike most other conserved miRNAs, these loci generally accumulate modest amounts of small RNAs and/or have atypical structural features. This might reflect that their processing is atypical and/or regulated, or that their conservation reflects a role other than, or in addition to miRNA-type function. For example, besides the pasha 5‘ UTR hairpins, two of these loci are located in CDS or 3’UTR, and thus cleavage could mediate host mRNA downregulation.

**Supplementary Figure S9:**
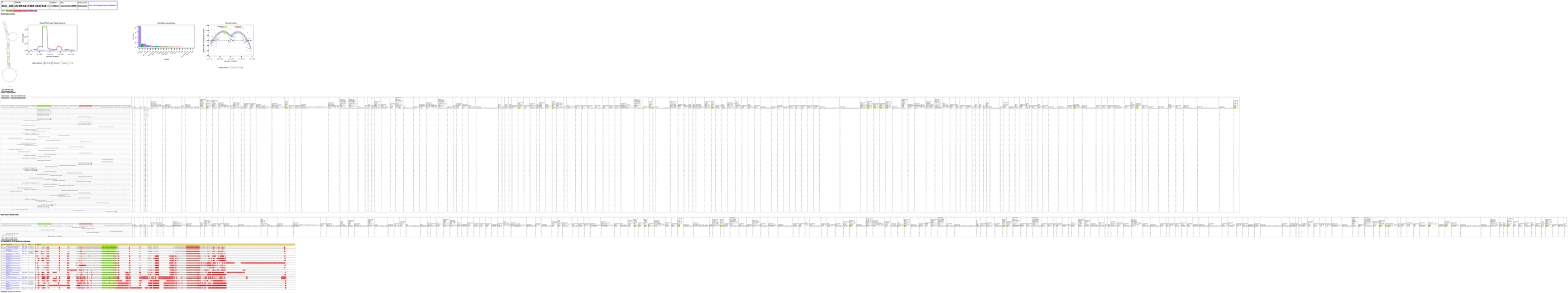
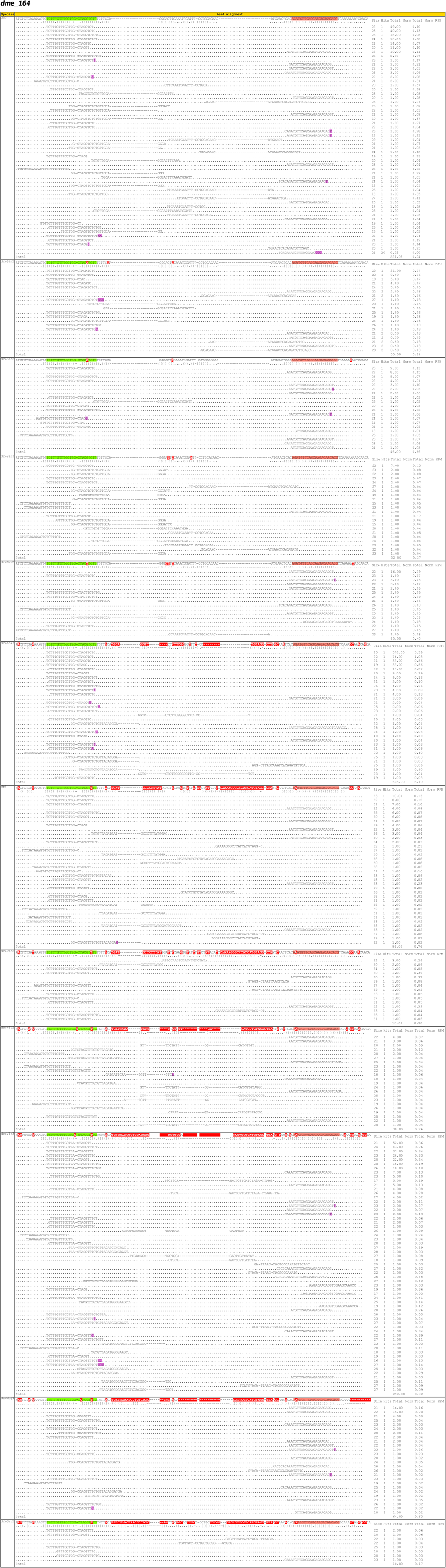
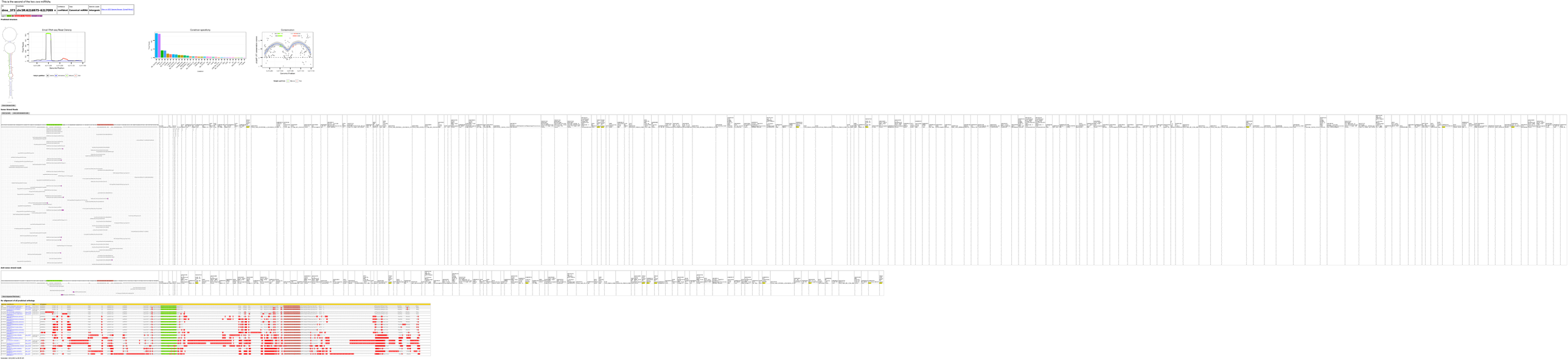
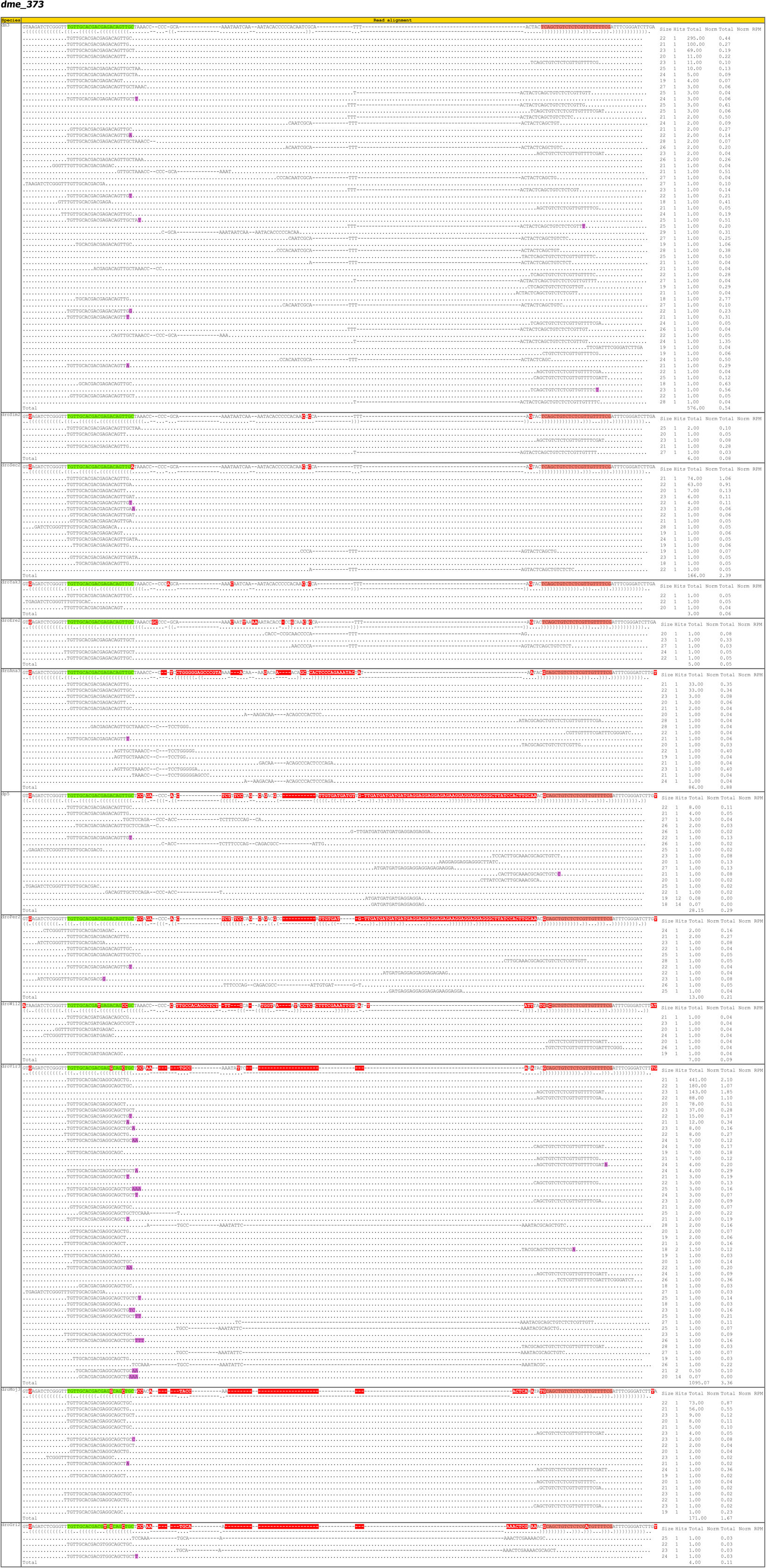
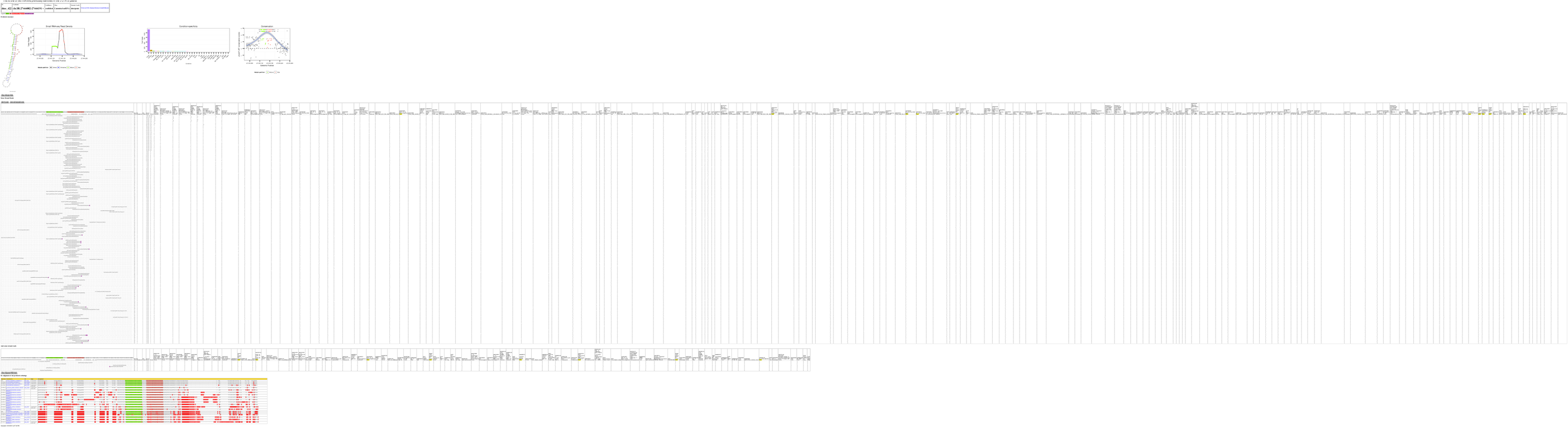
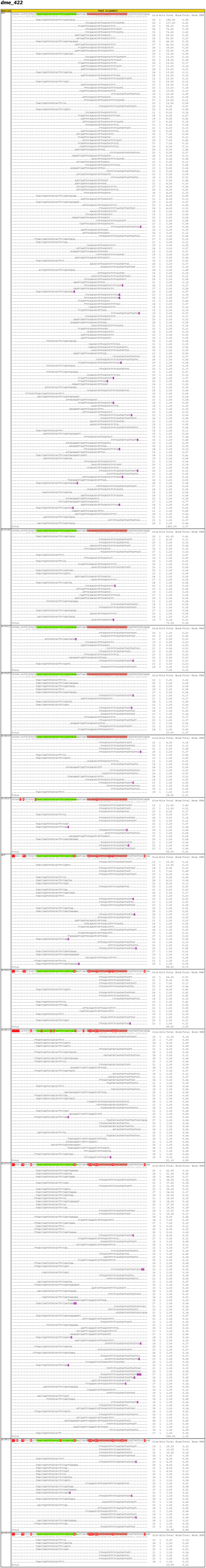
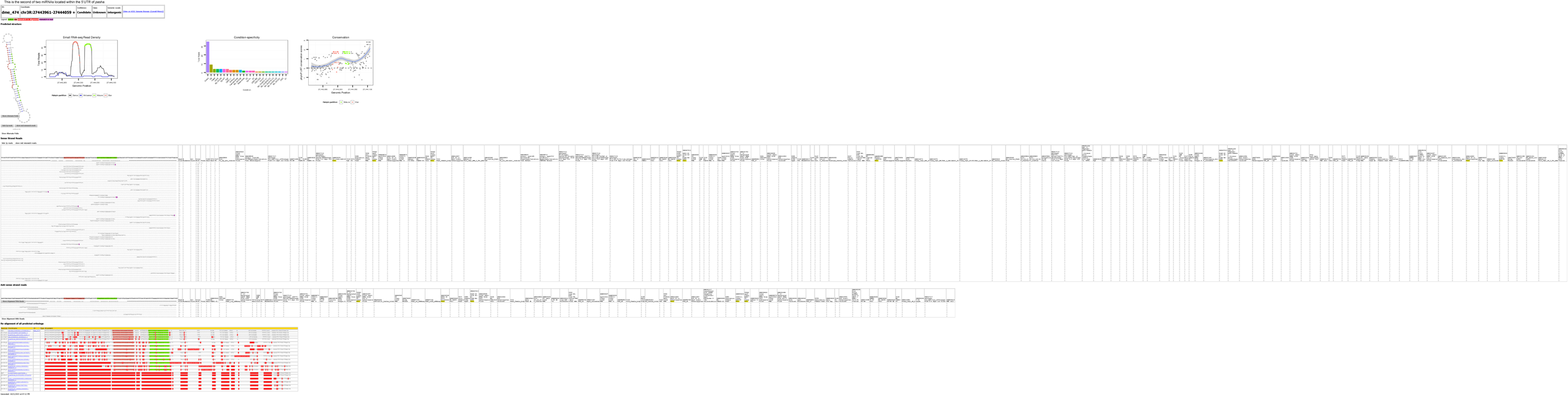
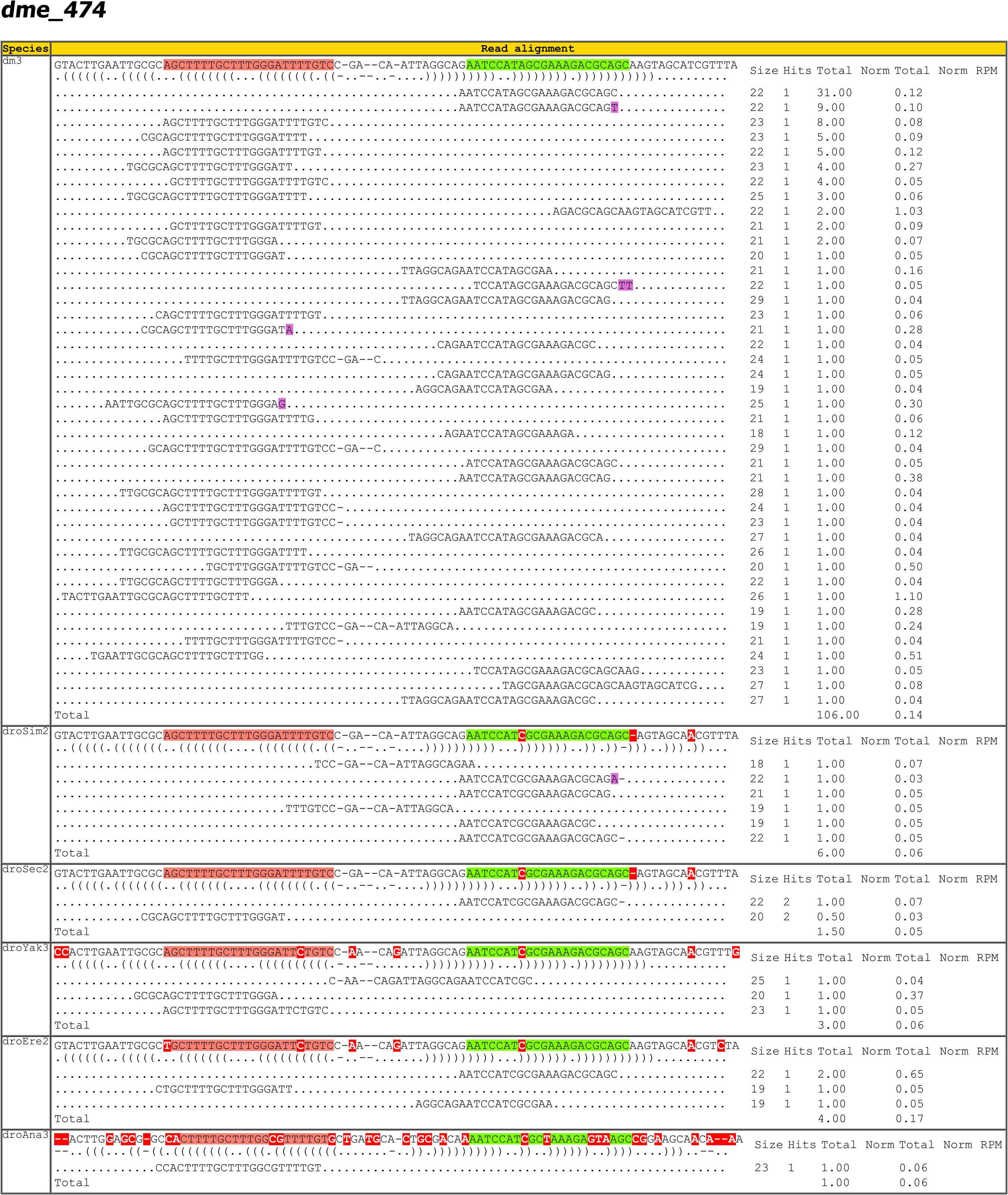
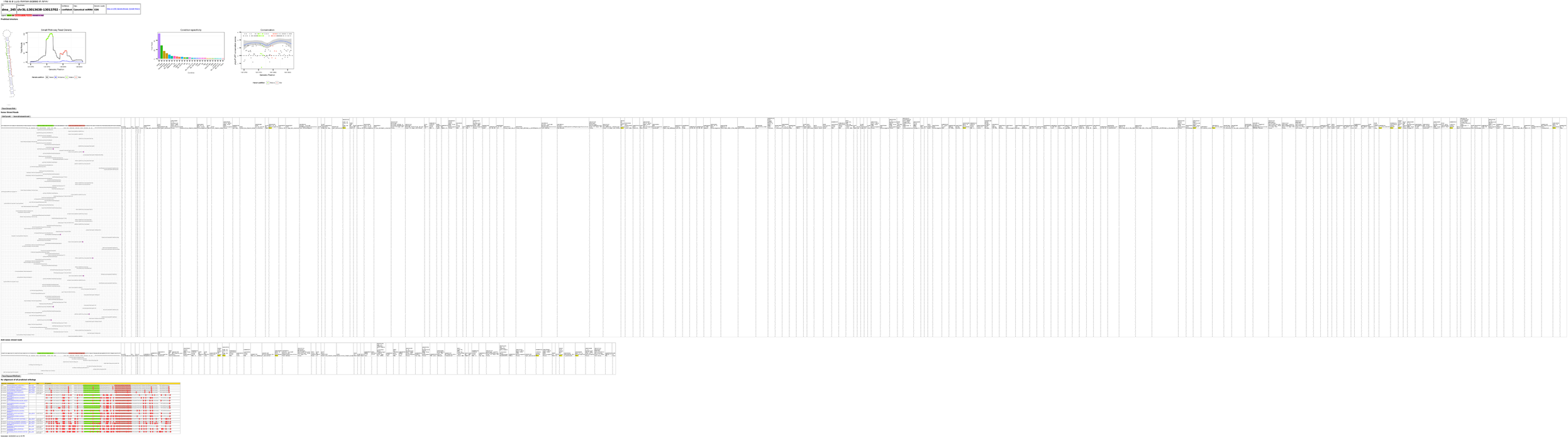
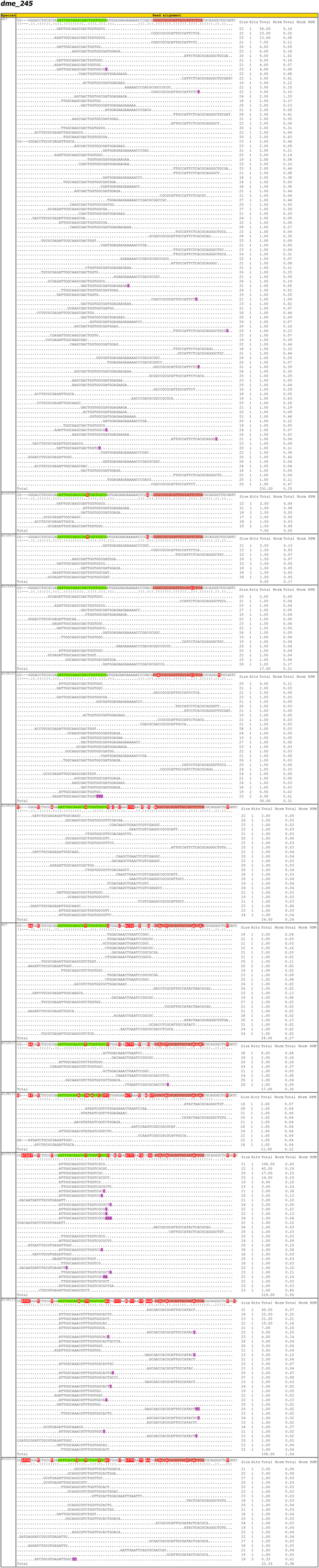
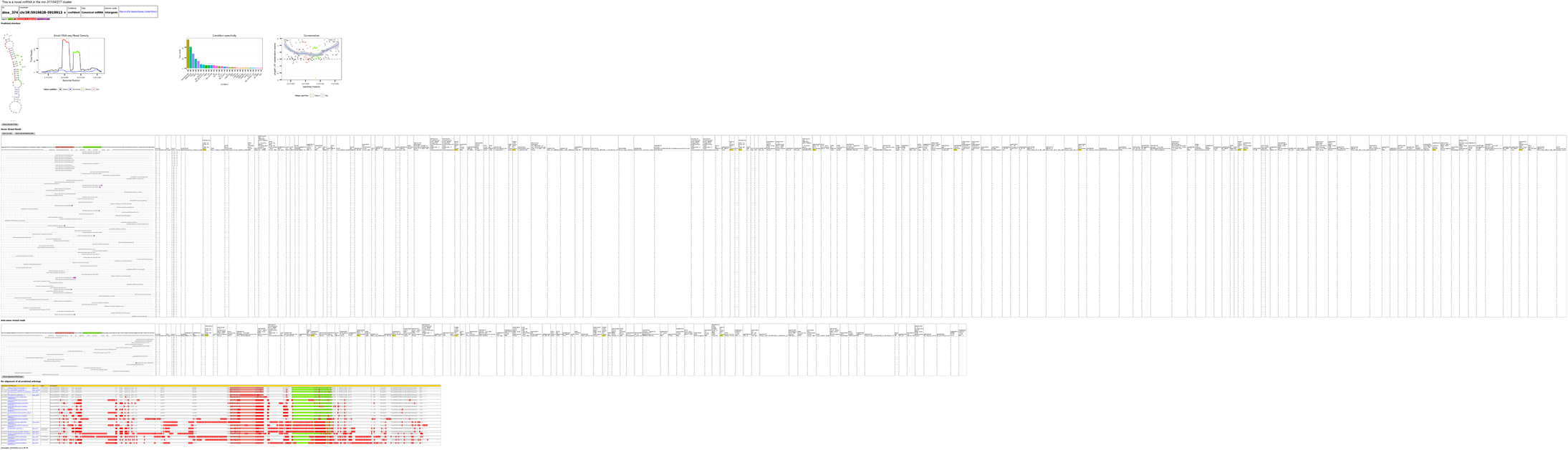
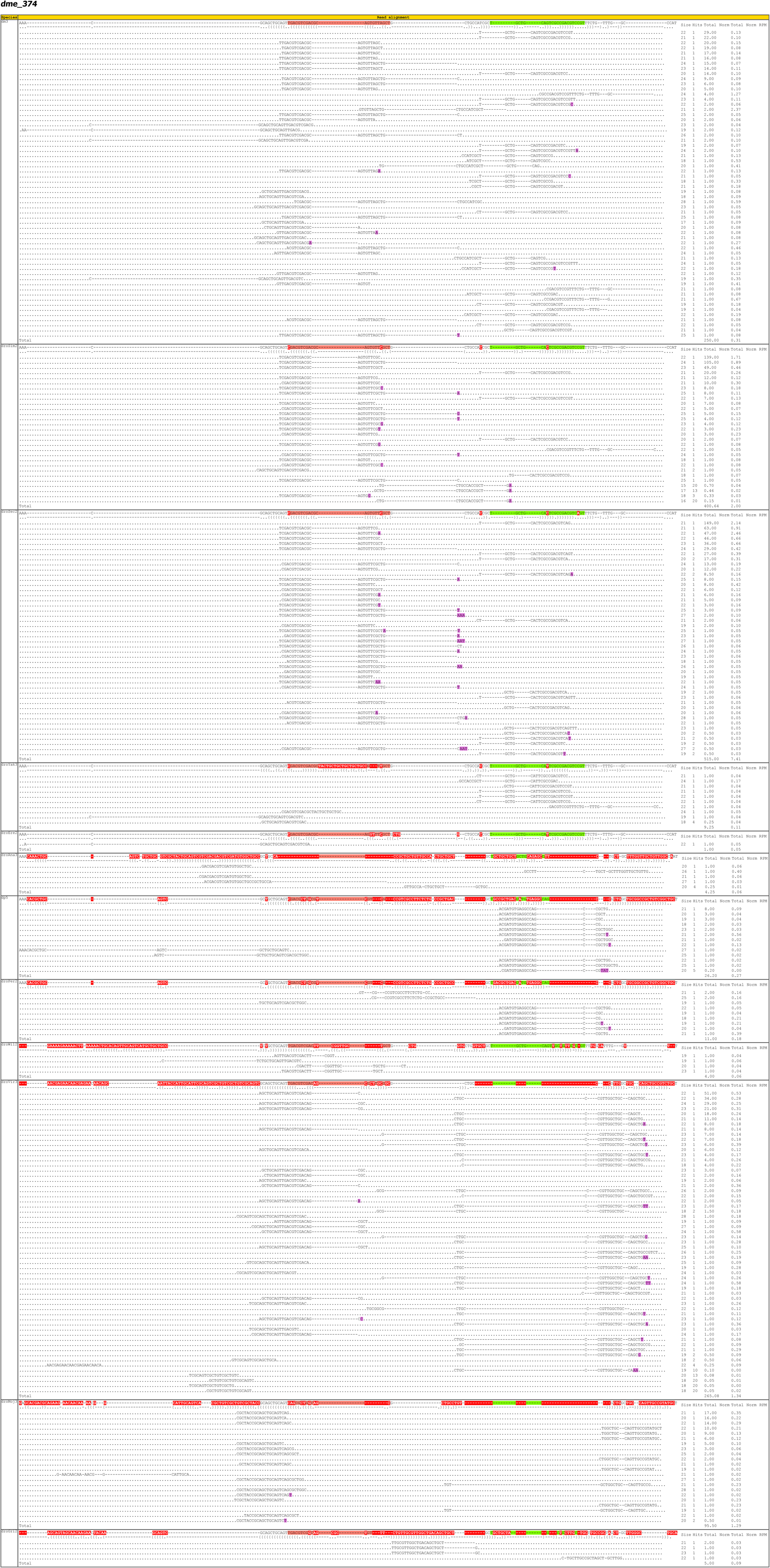
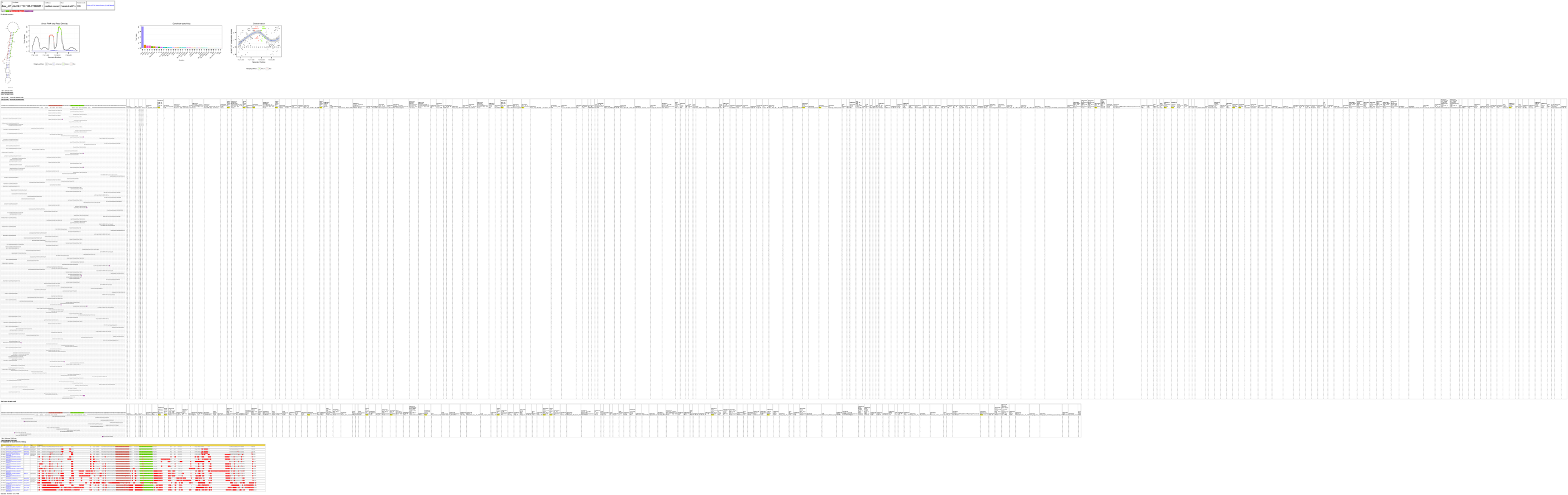
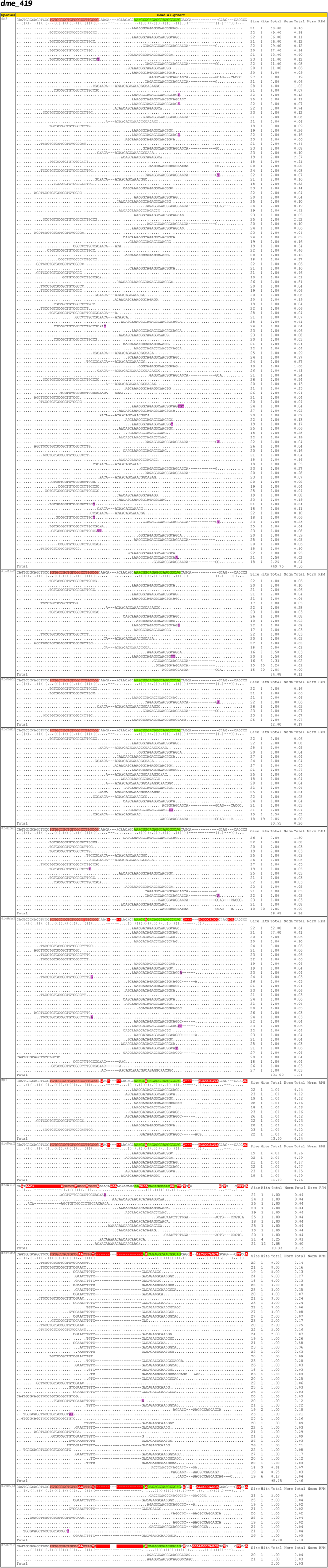
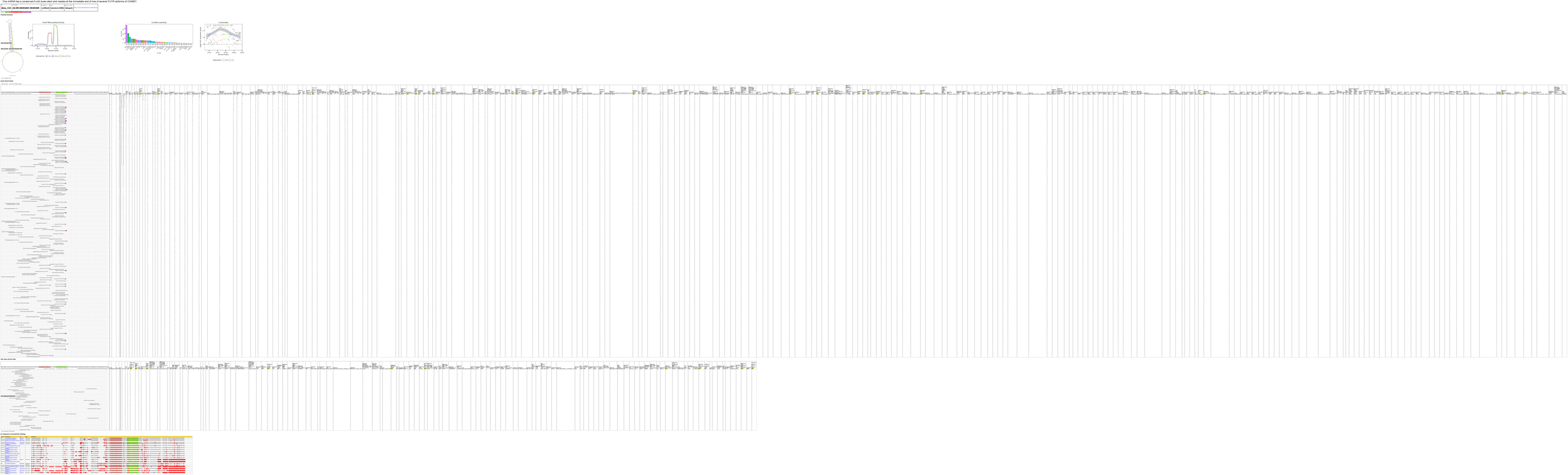
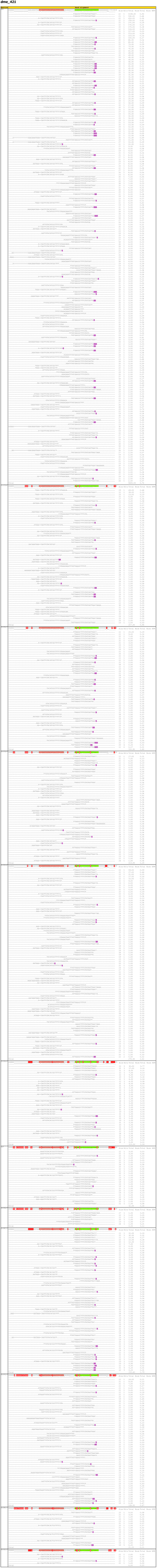
Alignment and small RNA read details for novel conserved miRNAs annotated in this study.

**Supplementary Figure S10:**
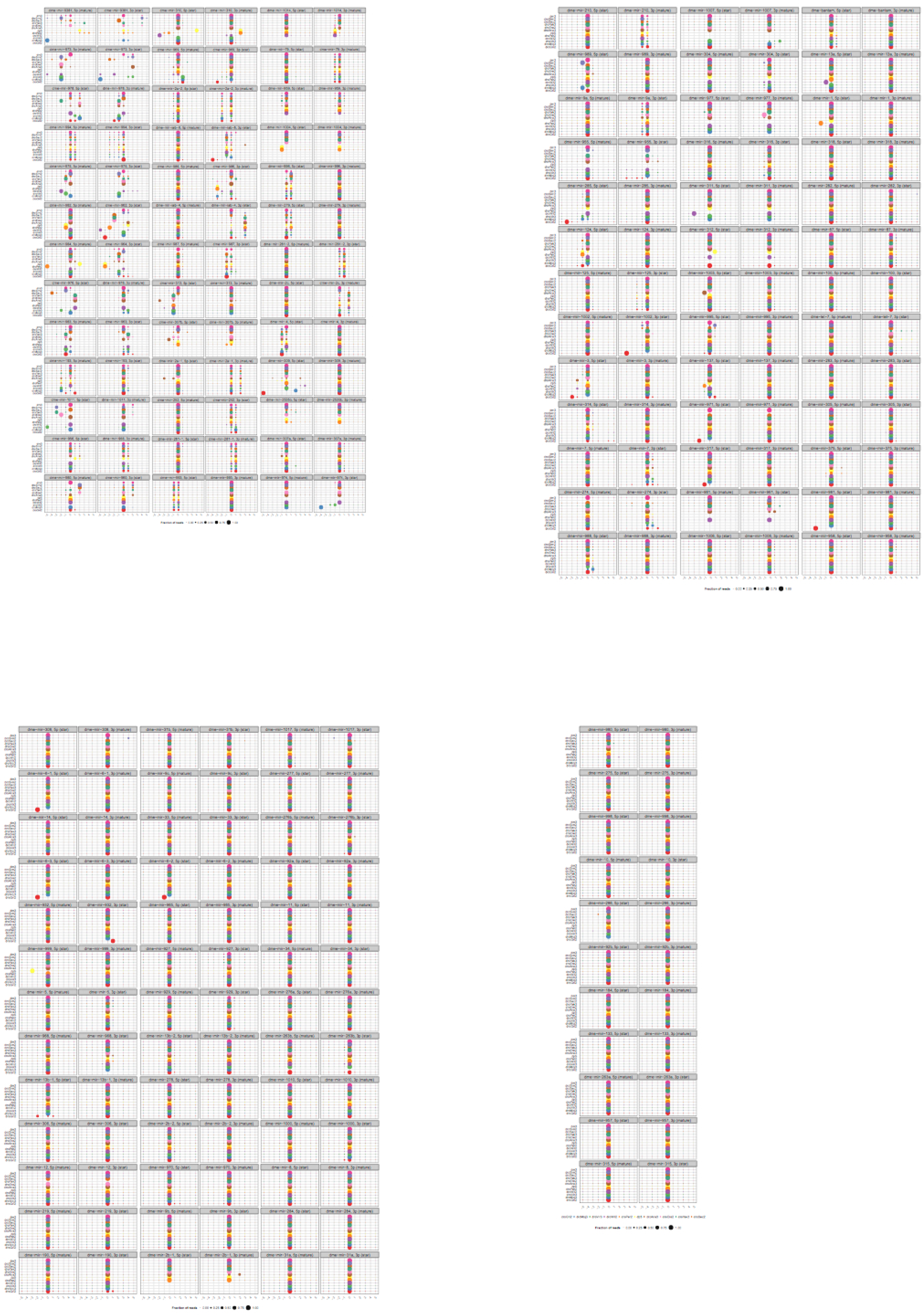
Evolutionary patterns of 5’ end cleavage precision for all conserved D. melanogaster miRNAs.

**Supplementary Figure S11:**
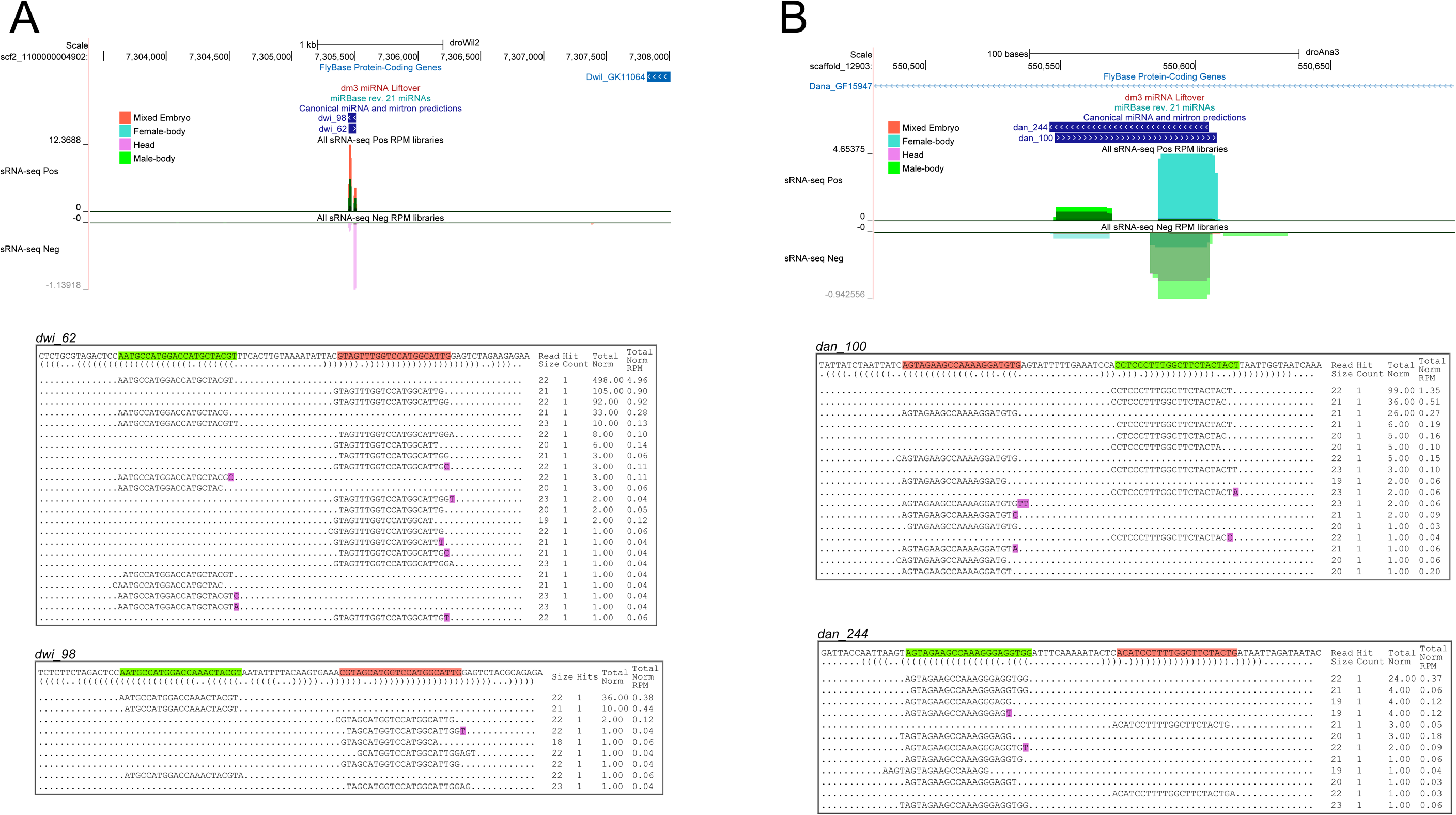
Additional examples of novel sense/antisense miRNA pairs. (A) Example of novel sense/antisense miRNA pair from D. willistoni (dwi_62/dwi-98). (B) Example of novel sense/antisense miRNA pair from D. ananassae (dan_100/dan_244).

**Supplementary Figure S12:**
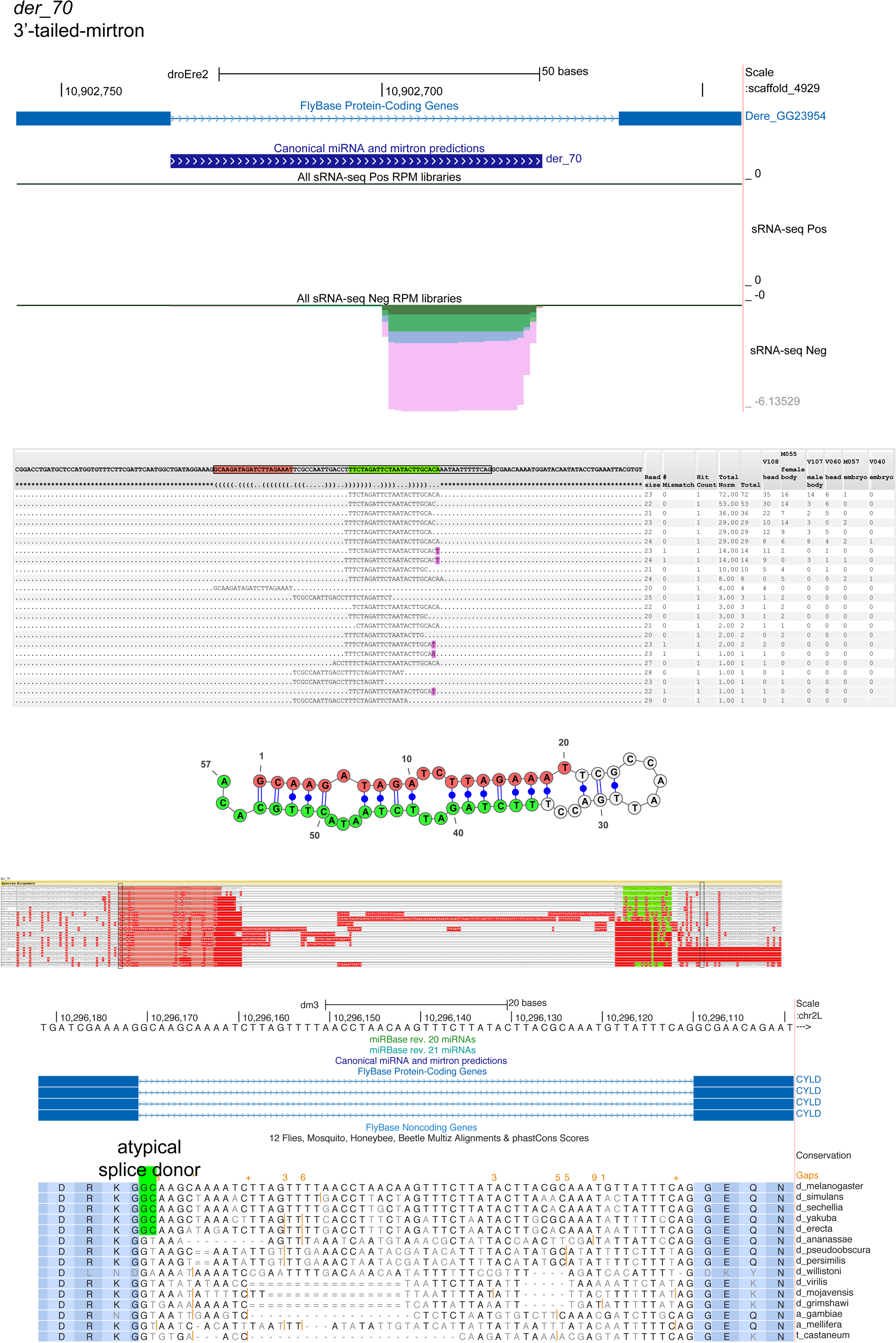
Example of a novel, atypical 3’ tailed mirtron. A 3’ tailed mirtron from D. erecta, Der_70. This locus produces a dominant 3p miRNA, which is trimmed by ∼13 from the splice acceptor site. Note that there is 3’ untemplated uridylation (purple) associated with a subset of Der_70-3p reads. While Der_70-5p reads are modest, there are also apparent partially diced products that include the terminal loop that phase precisely with the Der_70-5p species, supporting this as a Dicer position. Bottom alignment shows that Der_70 tailed mirtron resides in the conserved gene CYLD, and is associated with a non-canonical splice “GC” donor in the five melanogaster group species, which is instead a conventional “GT” splice donor in most other insects. Note D. willistoni seems to have lost both splice sites.

**Supplementary Figure S13:**
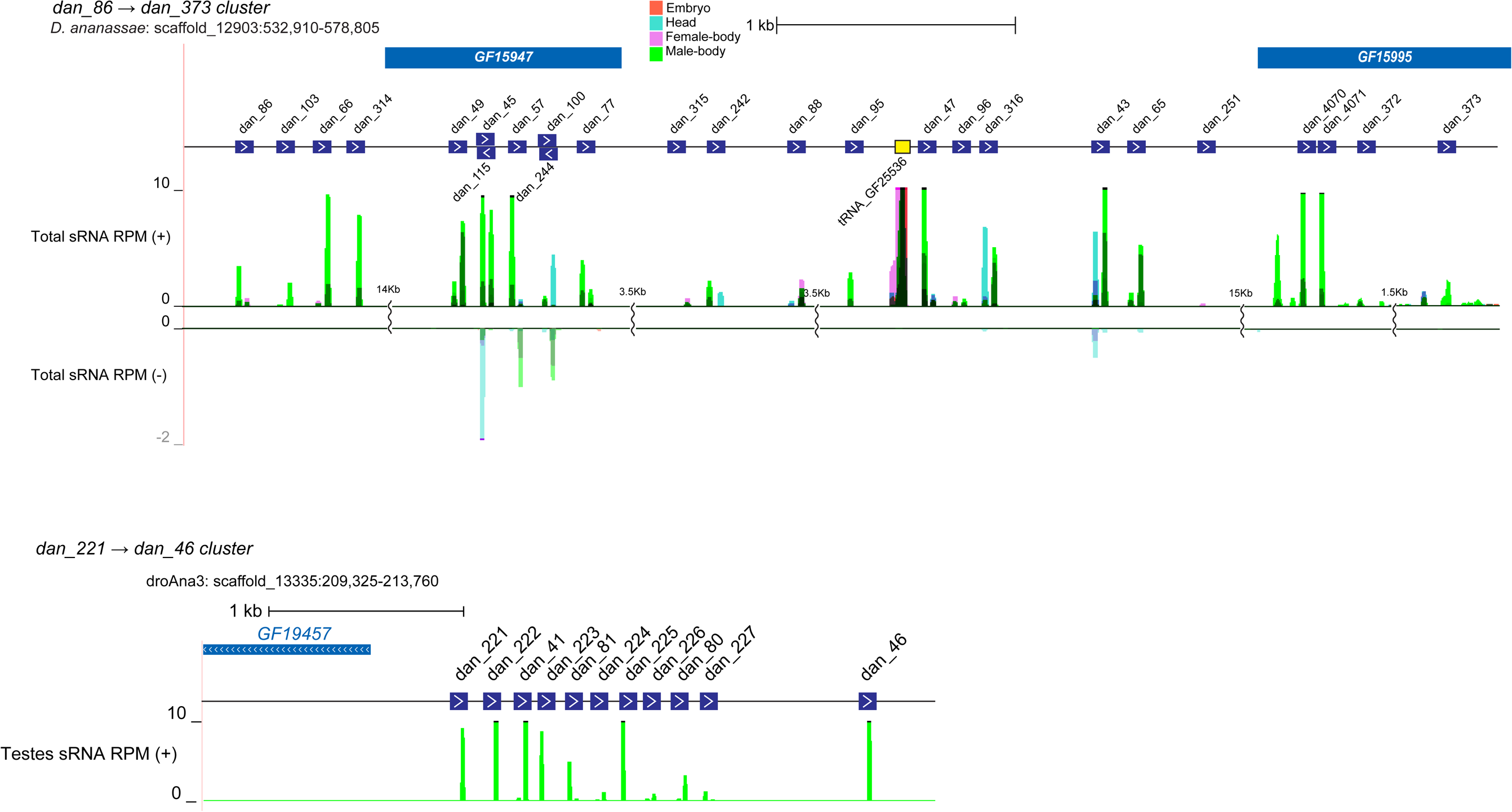
Annotation of novel Testes-restricted, recently-evolved, clustered (TRC) miRNA clusters identified within D. ananassae.

**Supplementary Figure S14:**
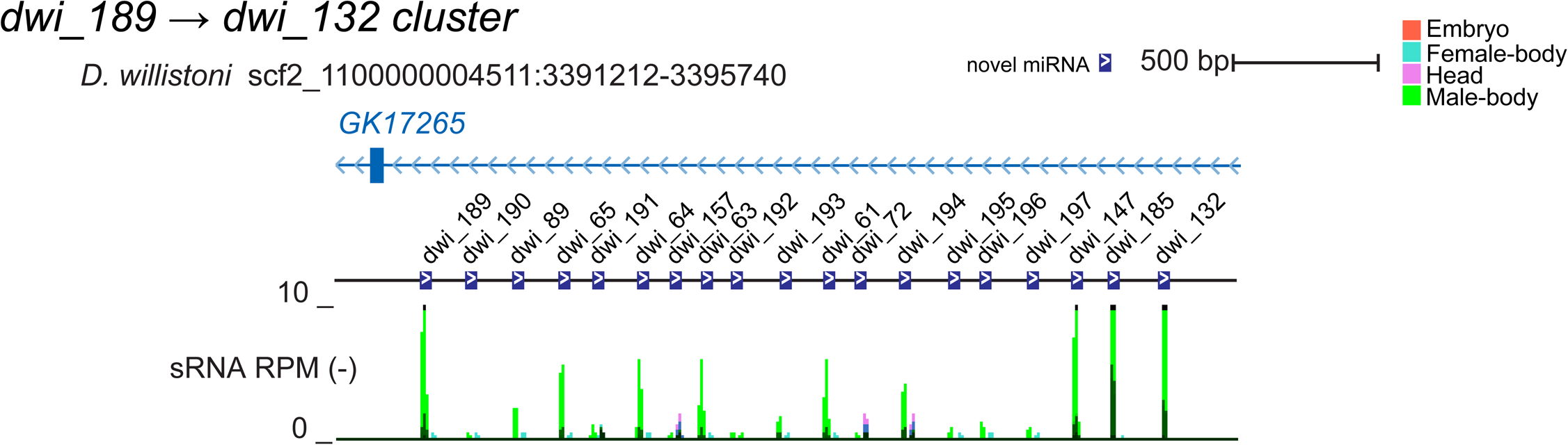
Annotation of a novel Testes-restricted, recently-evolved, clustered (TRC) miRNA cluster identified in D. willistoni. Tissue code indicates the miRNAs are all highest expressed in male body libraries.

**Supplementary Figure S15:**
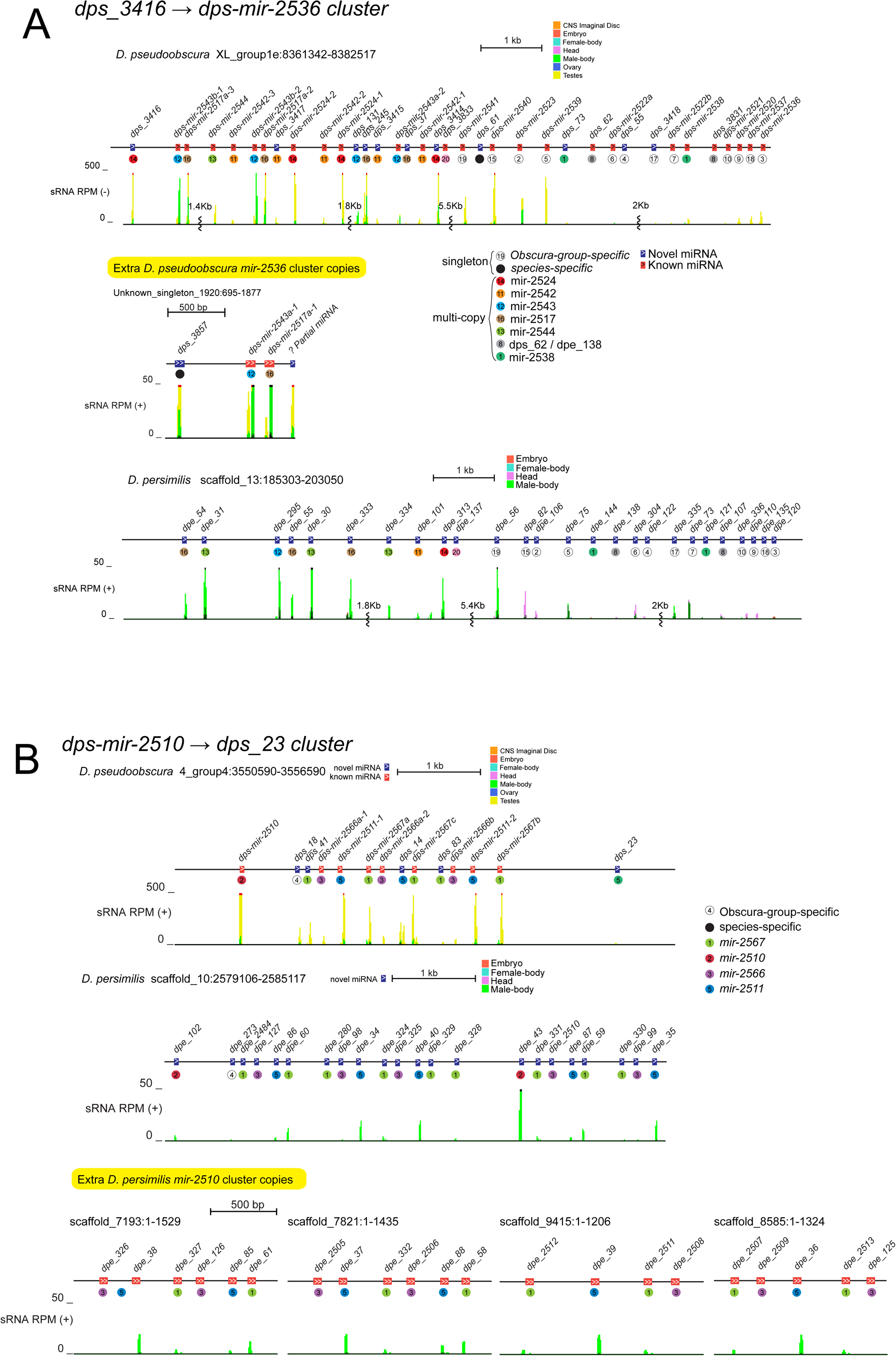
Annotation of novel Testes-restricted, recently-evolved, clustered (TRC) miRNA clusters identified in obscura group species, D. pseudoobscura and D. persimilis. (A) Genomic organization and small RNA read density of orthologous *dps_3416 → dps-mir-2536* TRC clusters in D. pseudoobscura and D. persimilis. (B) Genomic organization and small RNA read density of orthologous *dps-mir-2510 → dps_23* TRC clusters in D. pseudoobscura and D. persimilis. Tissue code indicates the miRNAs are all highest expressed in male body/testis libraries. Note that there are also additional copies of some of these TRC loci located outside of these clusters.

**Supplementary Figure S16:**
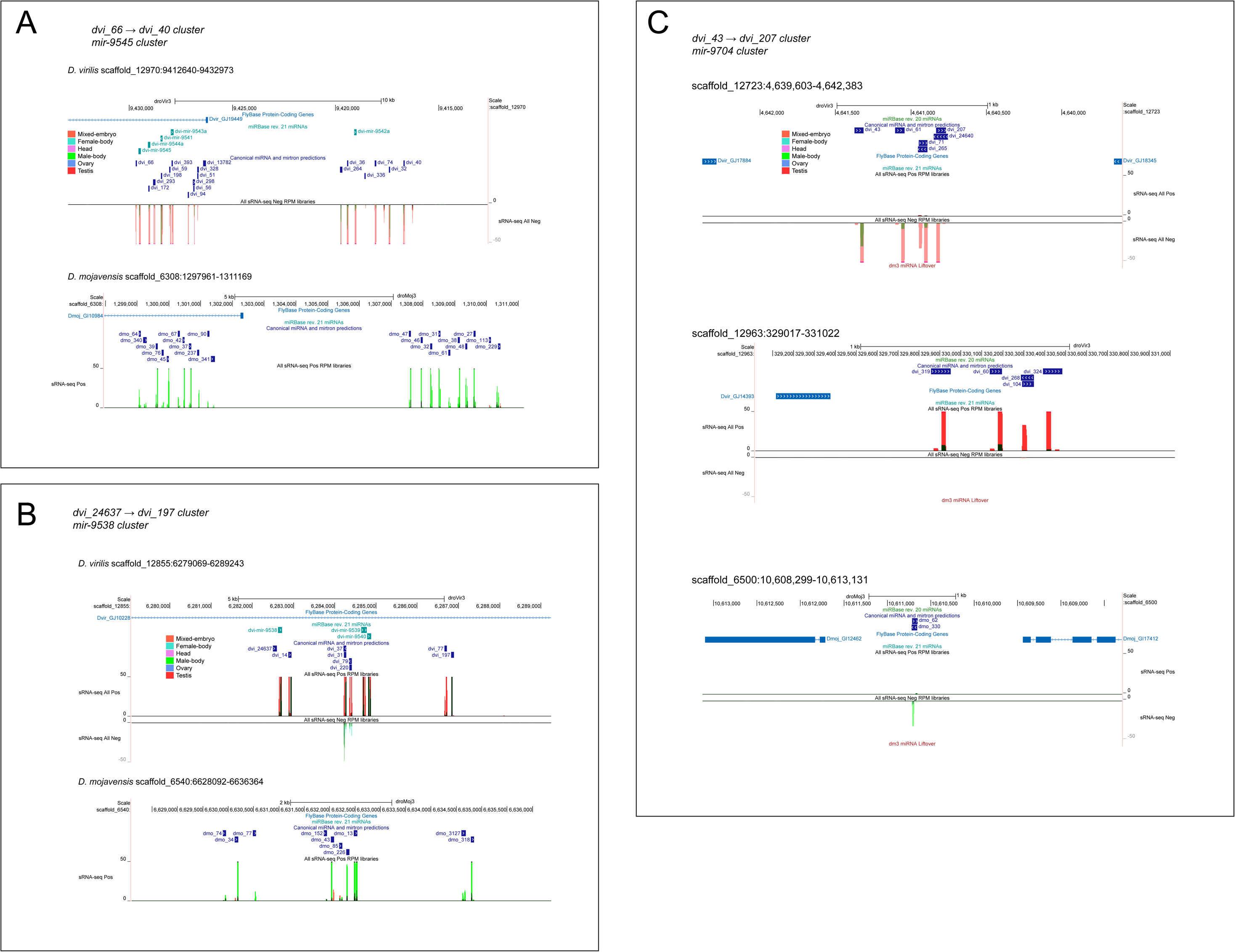
Annotation of novel Testes-restricted, recently-evolved, clustered (TRC) miRNA clusters identified in virilis clade species. (A) The dvi_66 → dvi_40 cluster. This cluster has a clear homologs in mojavensis. The small RNA mappings to their respective genomic loci are shown. (B) The dvi_24637 → dvi_197 cluster has clear homologs in D. mojavensis. The small RNA mappings to their respective genomic loci are shown. (C) Annotation of three additional novel Testes-restricted, recently-evolved, clustered (TRC) miRNA clusters identified in virilis clade species. The dvi_43 → dvi_207 cluster has two copies in D. virilis (i.e. roughly similar members can be found on scaffold_12723 and scaffold_12963). The other cluster is D. mojavensis dmo_62 and dmo_330.

**Supplementary Figure S17:**
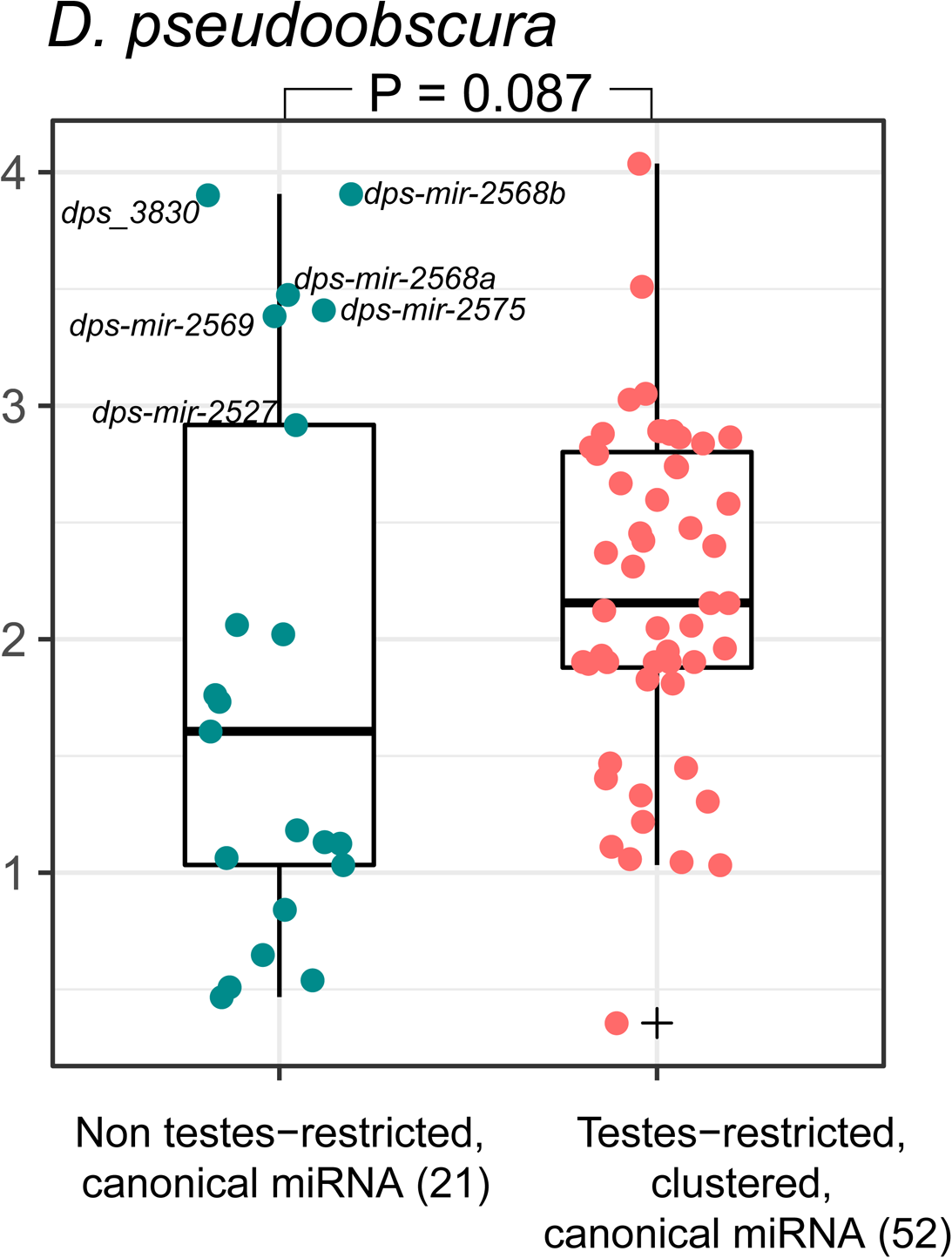
Expression difference between D. pseudoobscura-specific or obscura-group-specific TRC and solo canonical miRNAs. Points reflect the maximum expression per locus assessed over all D. pseudoobscura libraries. P-value computed from twotailed Wilcoxon Rank Sum Test.

**Supplementary Figure S18:**
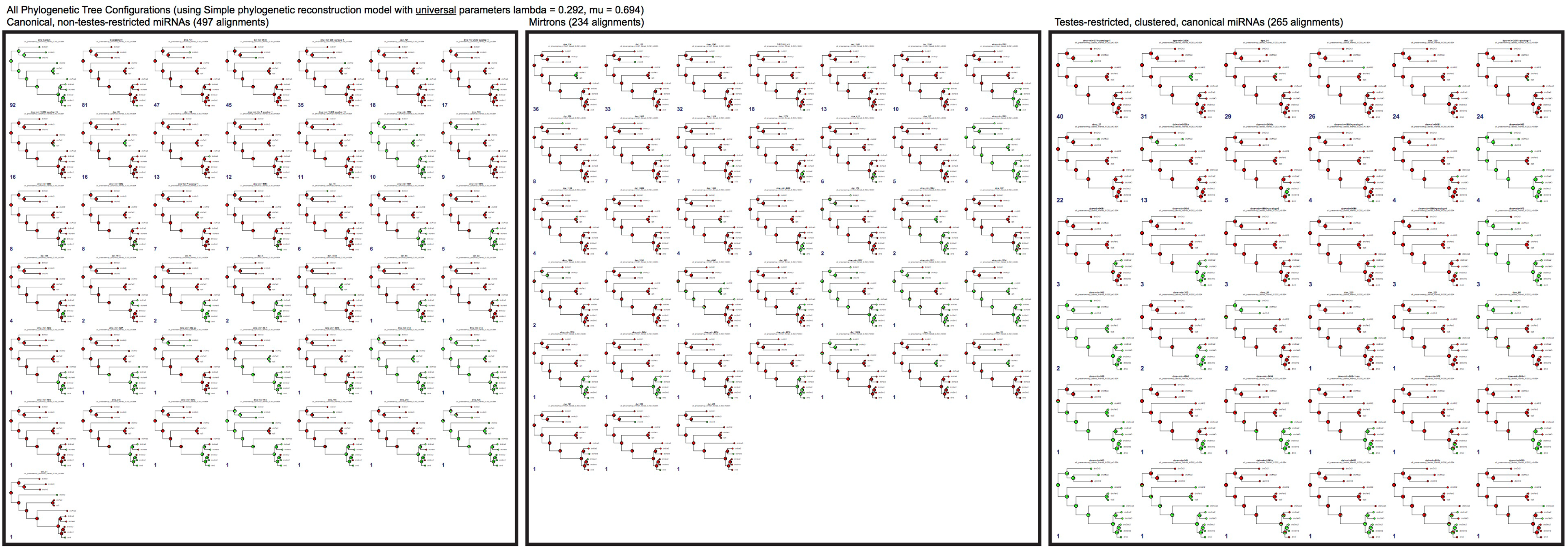
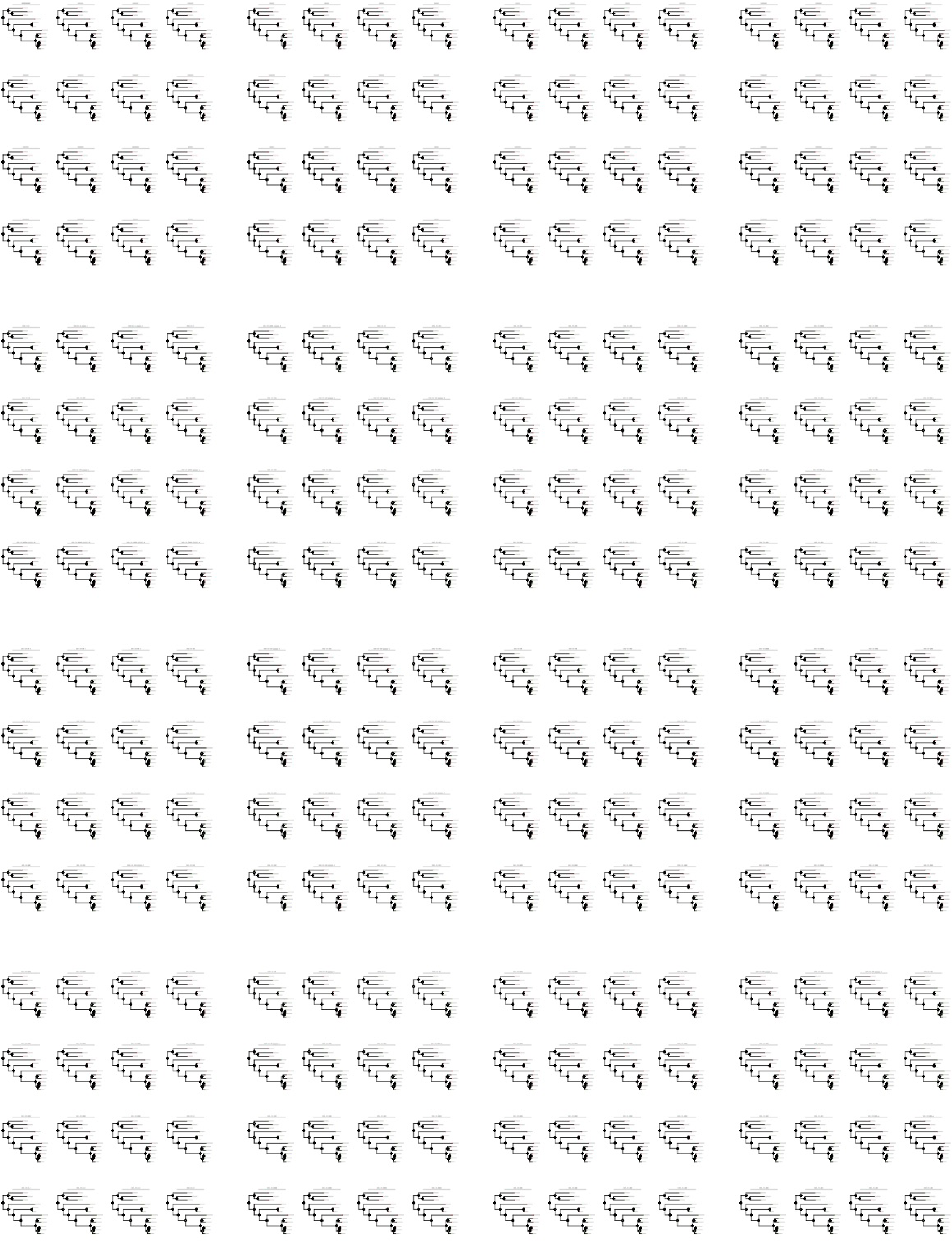
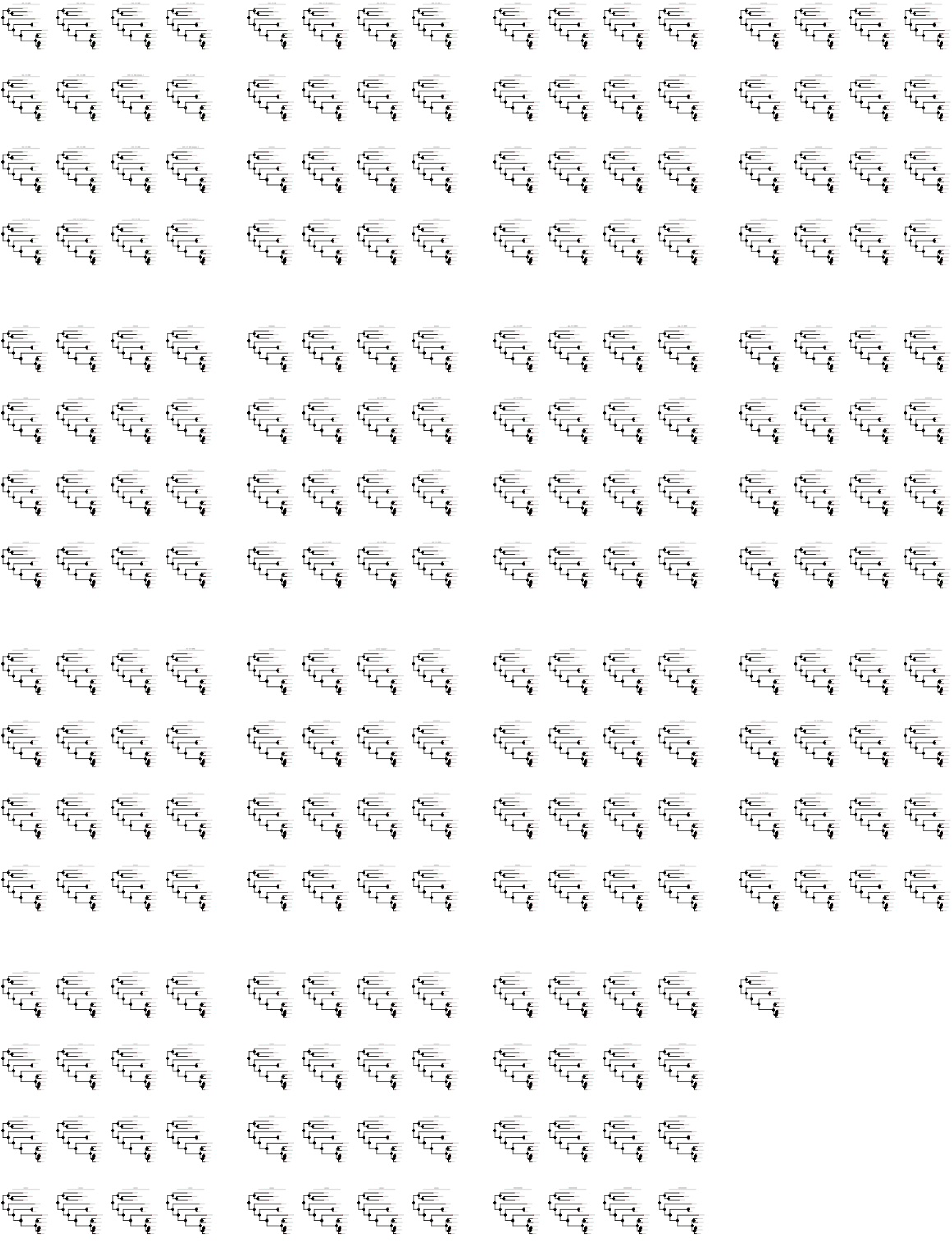
All possible phylogenetic reconstruction of ancestral miRNA presence and absence for 3 miRNA classes using a phylogenetic probabilistic graphical model with universal parameters of λ = 0.292 and μ = 0.694. These parameters were computed by running the phylogenetic reconstruction algorithms on all mirtrons and miRNAs pooled together. These trees illustrate how the method’s maximum likelihood reconstruction performs for all possible configurations of extant miRNAs presence and absence per alignment. Blue text indicates count of alignments with this particular configuration in each class. Summary of miRNA birth and death (Figure 6) are based upon these estimates of ancestral miRNA presence and absence.

**Supplementary Figure S19:**
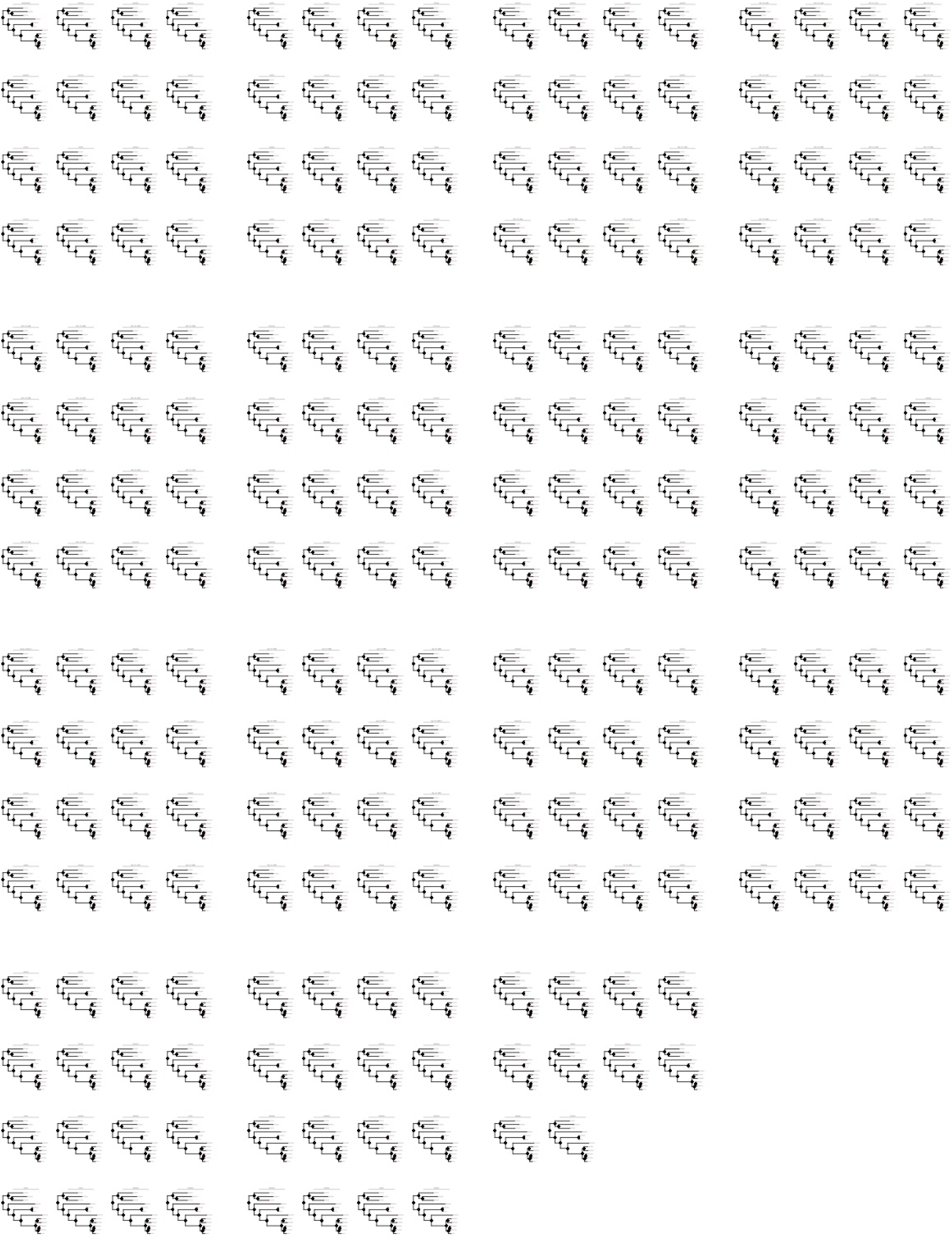
Individual phylogeny of extant and inferred ancestral miRNA presence and absence for all canonical miRNAs that are not in the testis-restricted clustered subclass.

**Supplementary Figure S20:**
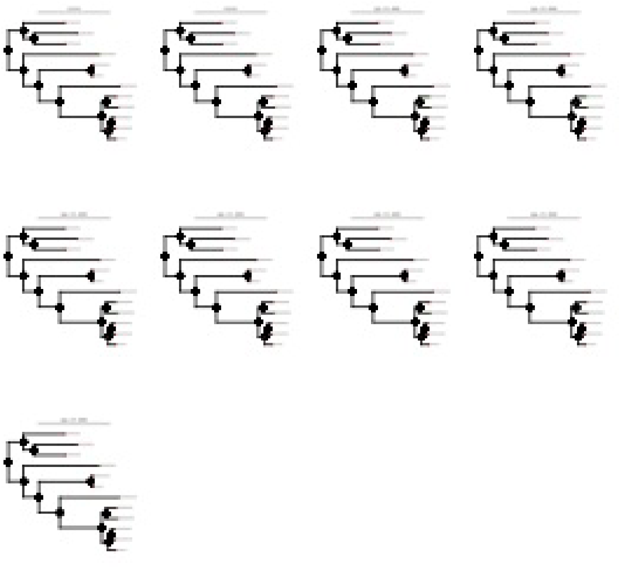
Individual phylogeny of extant and inferred ancestral miRNA presence and absence for all mirtrons.

**Supplementary Figure S21:**
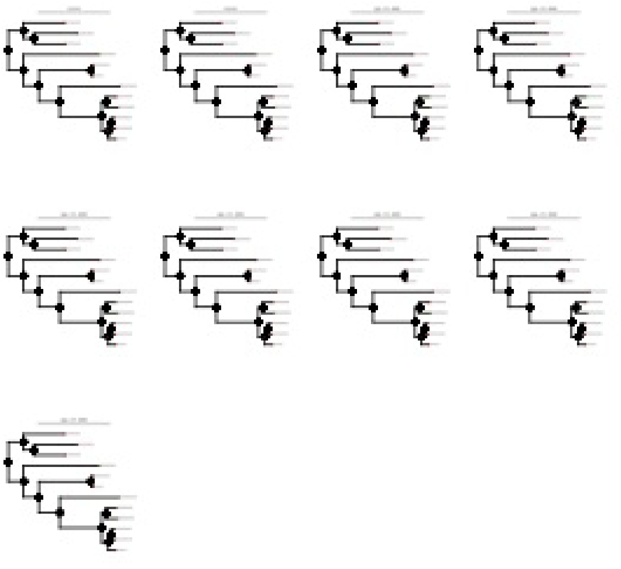
Individual phylogeny of extant and inferred ancestral miRNA presence and absence for all testis-restricted clustered (TRC) canonical miRNAs.

**Supplementary Figure S22:**
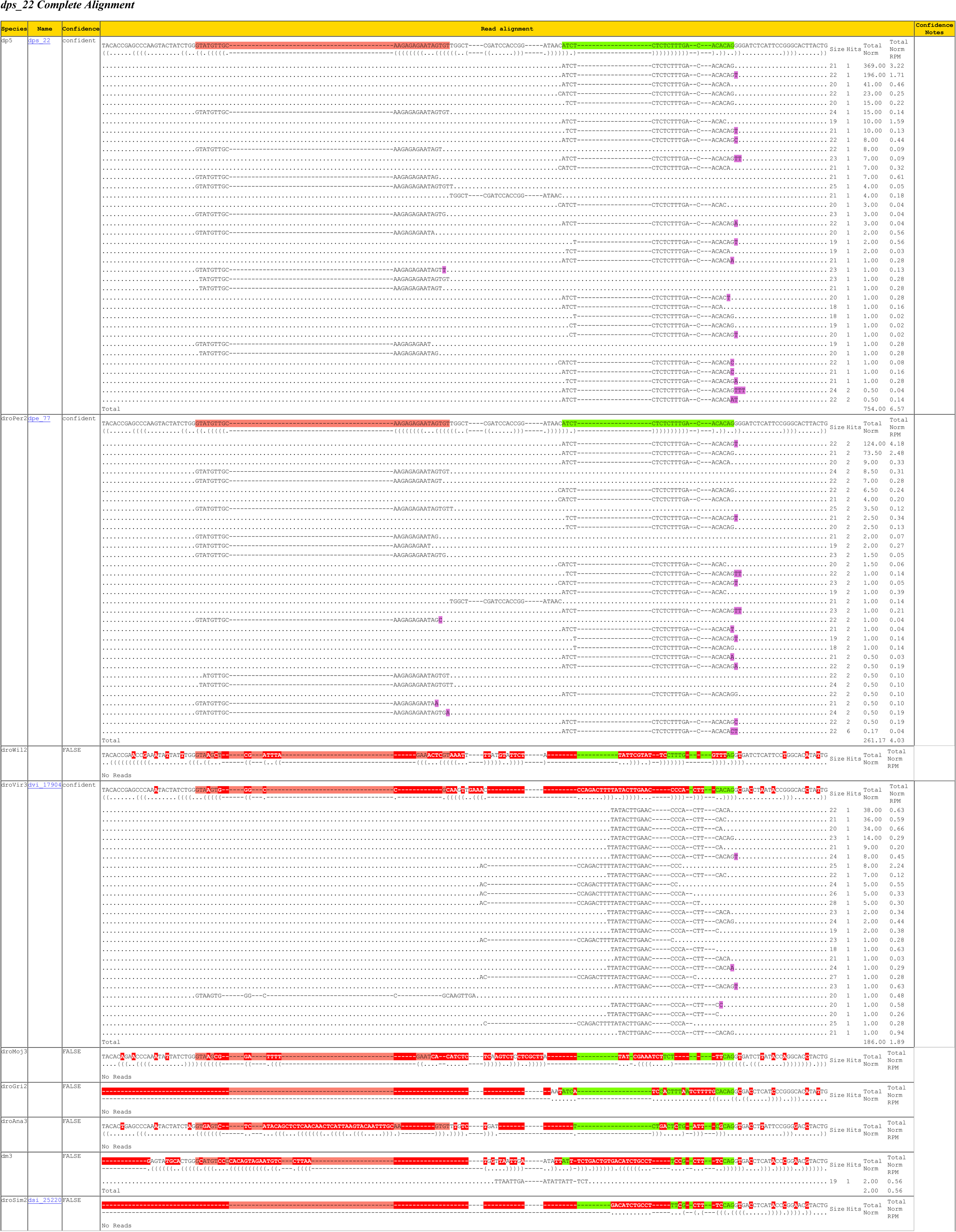
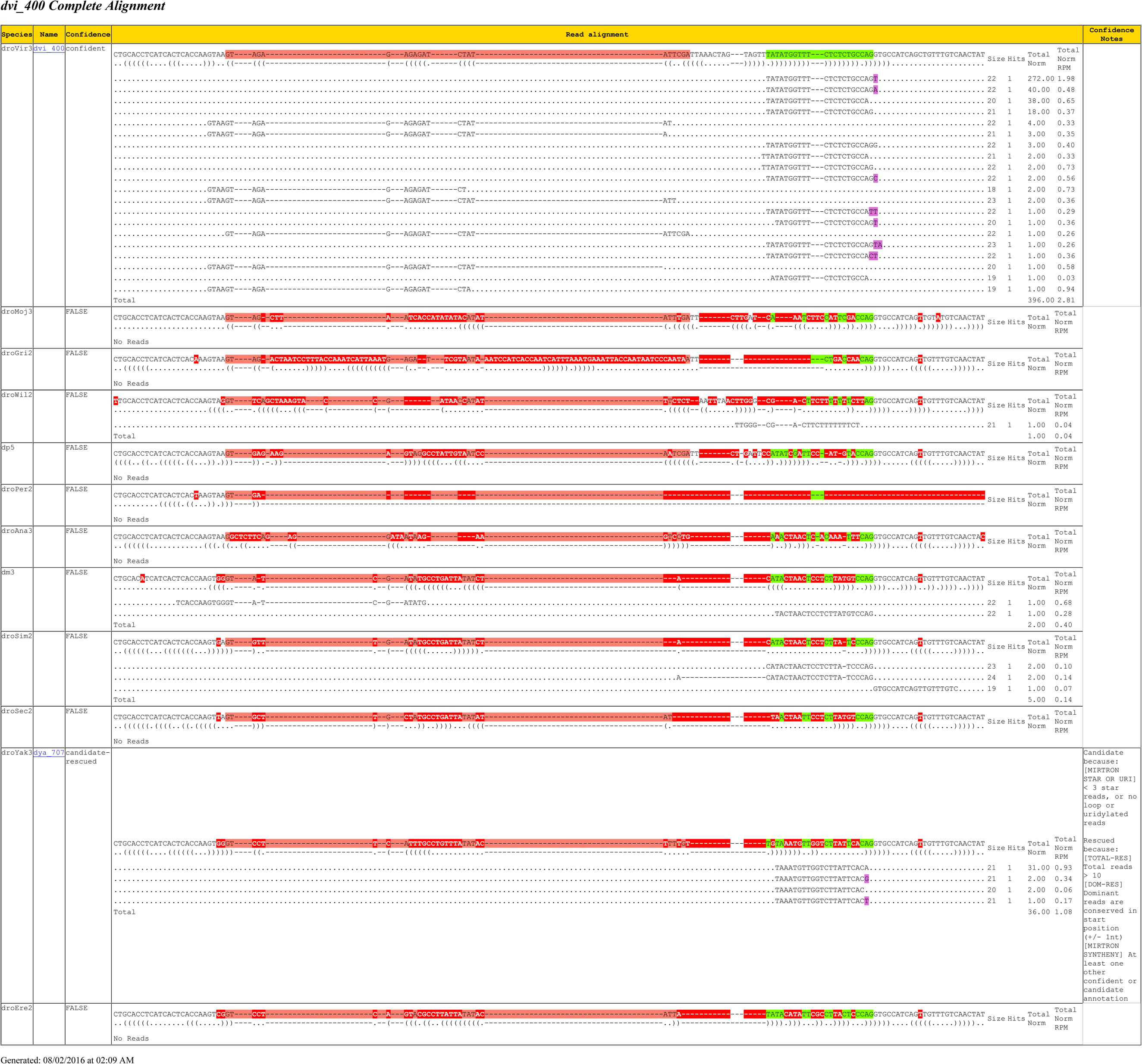
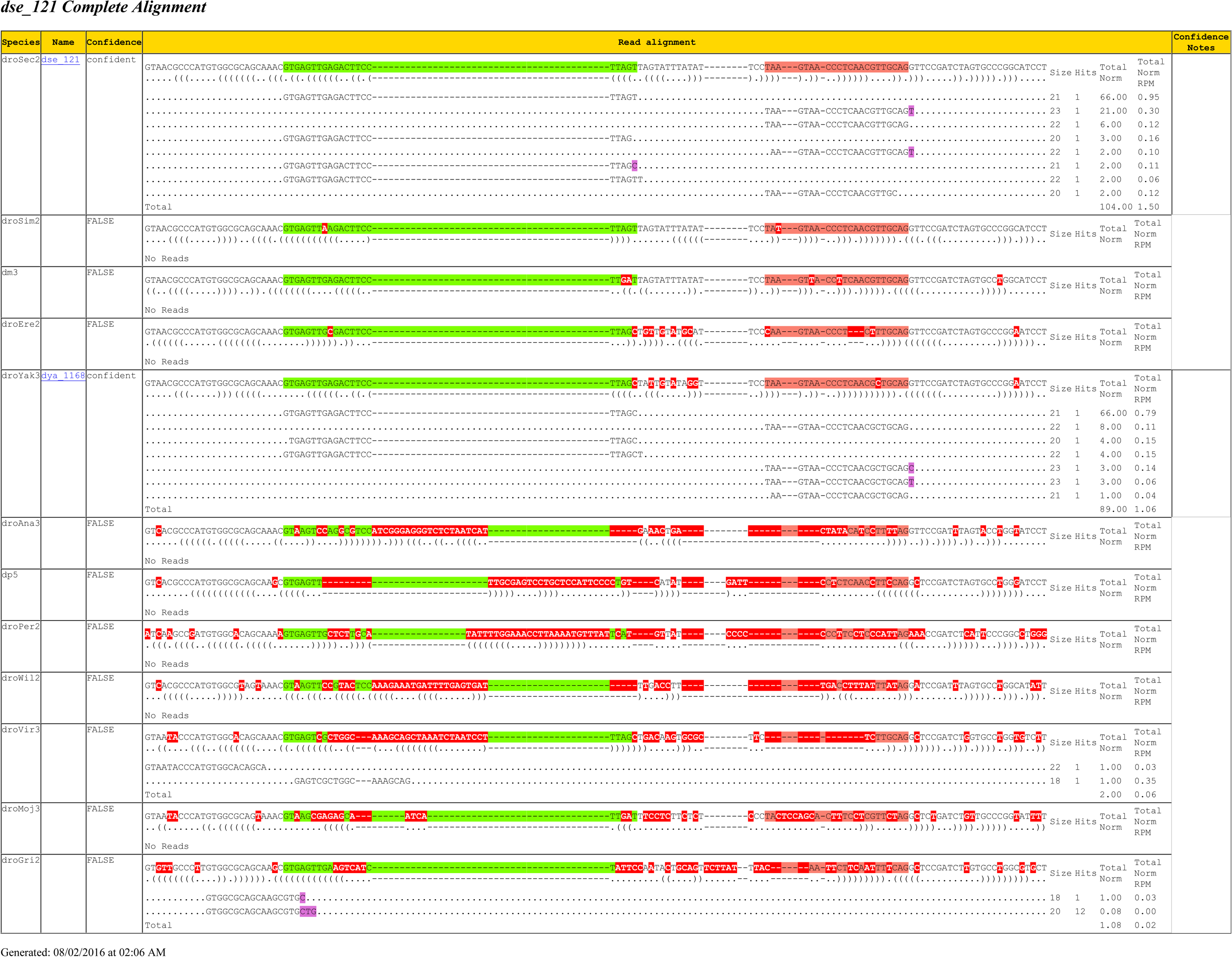
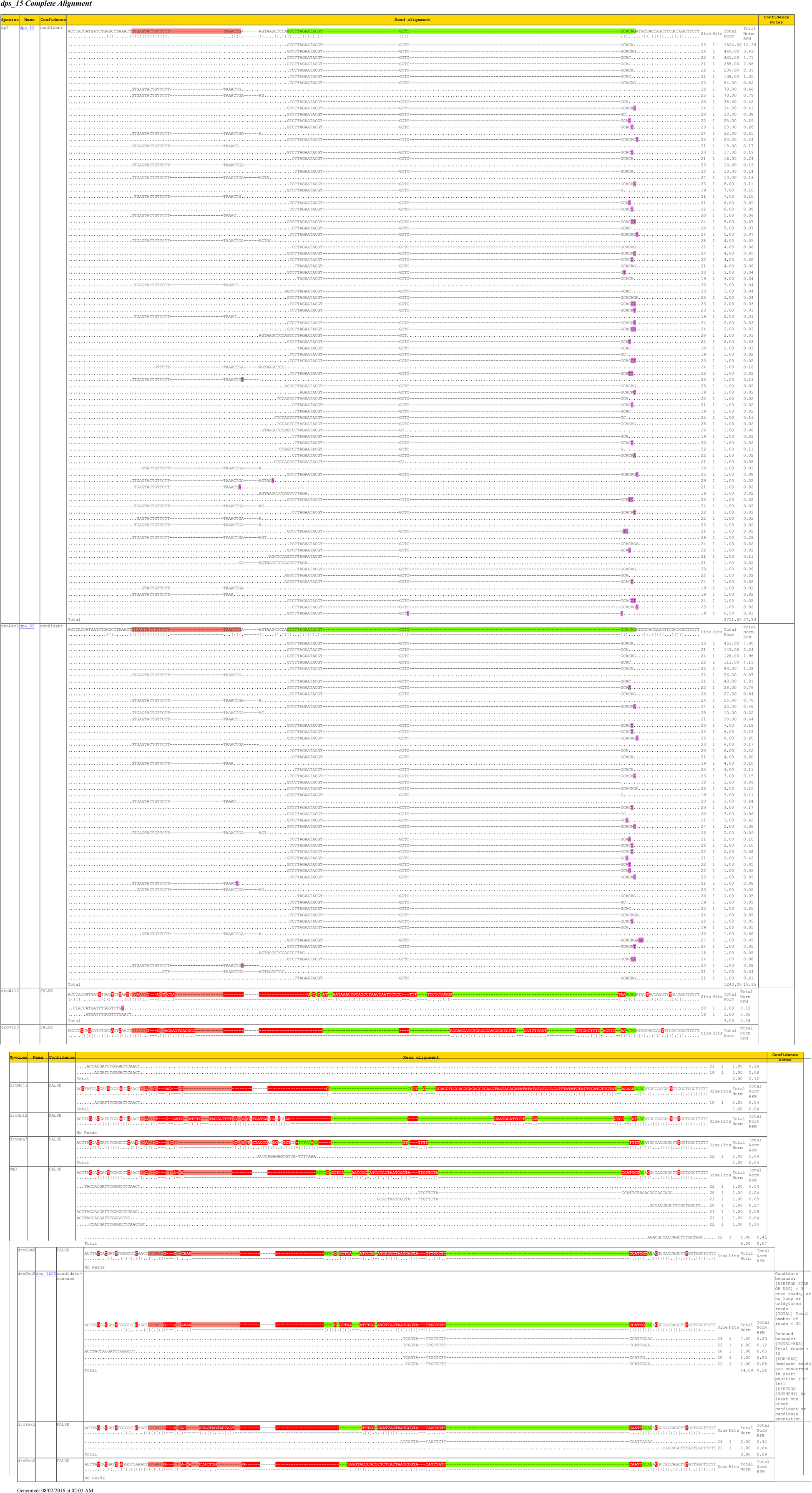
Examples of mirtrons with atypical emergence and decay patterns. Expression profiles and mirtron alignments are shown per example to highlight the non-clade specificity of mirtron presence.

**Supplementary Figure S23:**
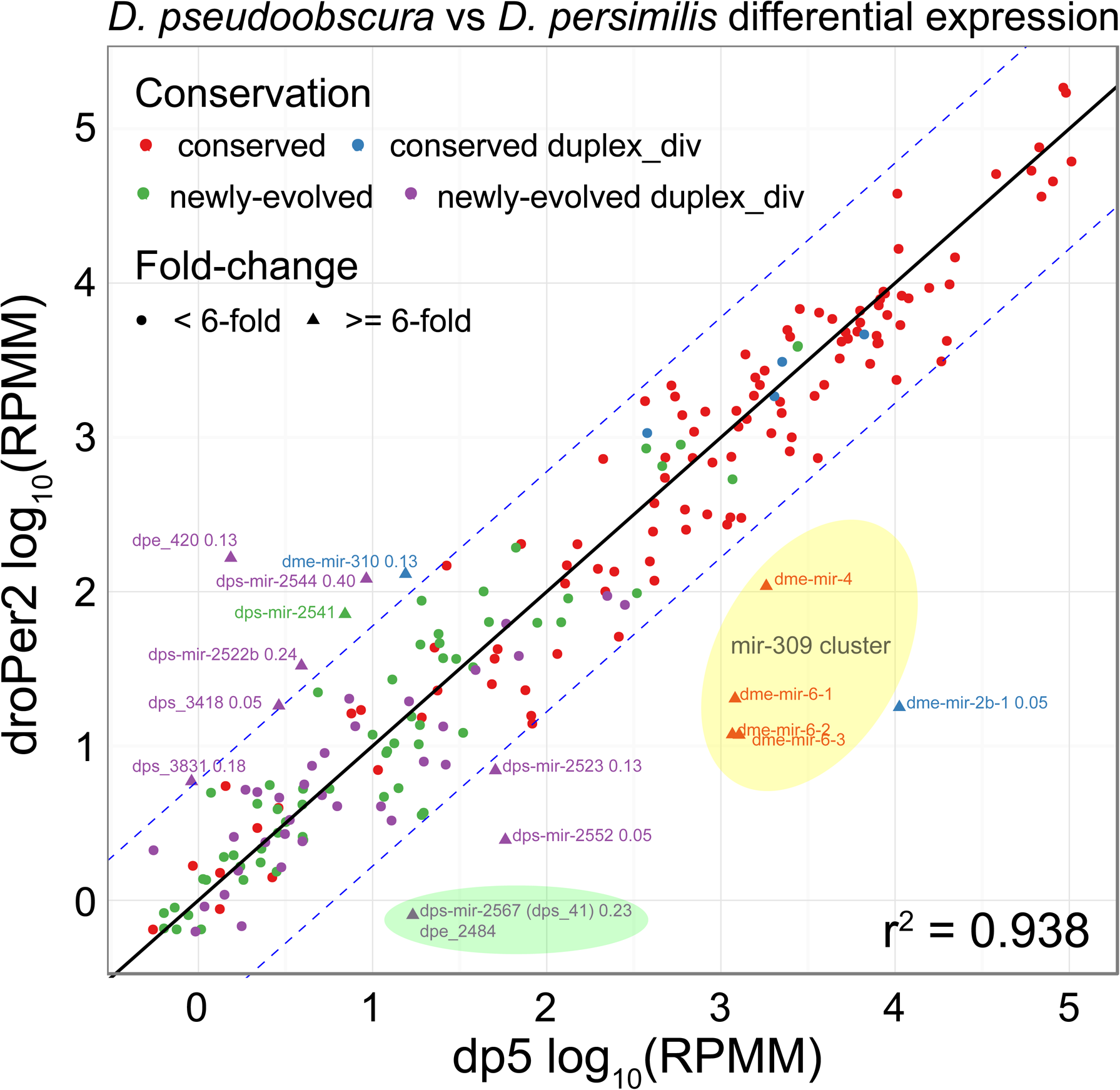
Differential expression of miRNAs between obscura group species. Scatterplot depicting the correlation of miRNA expression of all D. pseudoobscura and D. persimilis ortholog pairs (RPMM = Reads Per Million Mapped MiRNA Reads). All miRNA alignments with orthologs in both species are shown. Points that lie on or near the diagonal represent similarly expressed ortholog pairs. Orthologs with > 6-fold log10(RPMM) difference (denoted by the blue-dashed line and labeled points) are examples of significantly differentially expressed orthologs. Points are colored by miRNA age, and shapes represent miRNAs with or without miR:miR* duplex region substitutions (fraction of duplex sites with substitutions are labeled). Note that the mir-309 cluster (yellow) loci are expected to be expressed in the very early embryo, and given that the embryo development and timing were not controlled in library preparation, their differential accumulation may not be genuine. Amongst loci changed by >6-fold, dme-mir-2b-1 and dme-mir-310 are deeply conserved, but all others are specific to the obscura group species.

**Supplementary Figure S24:**
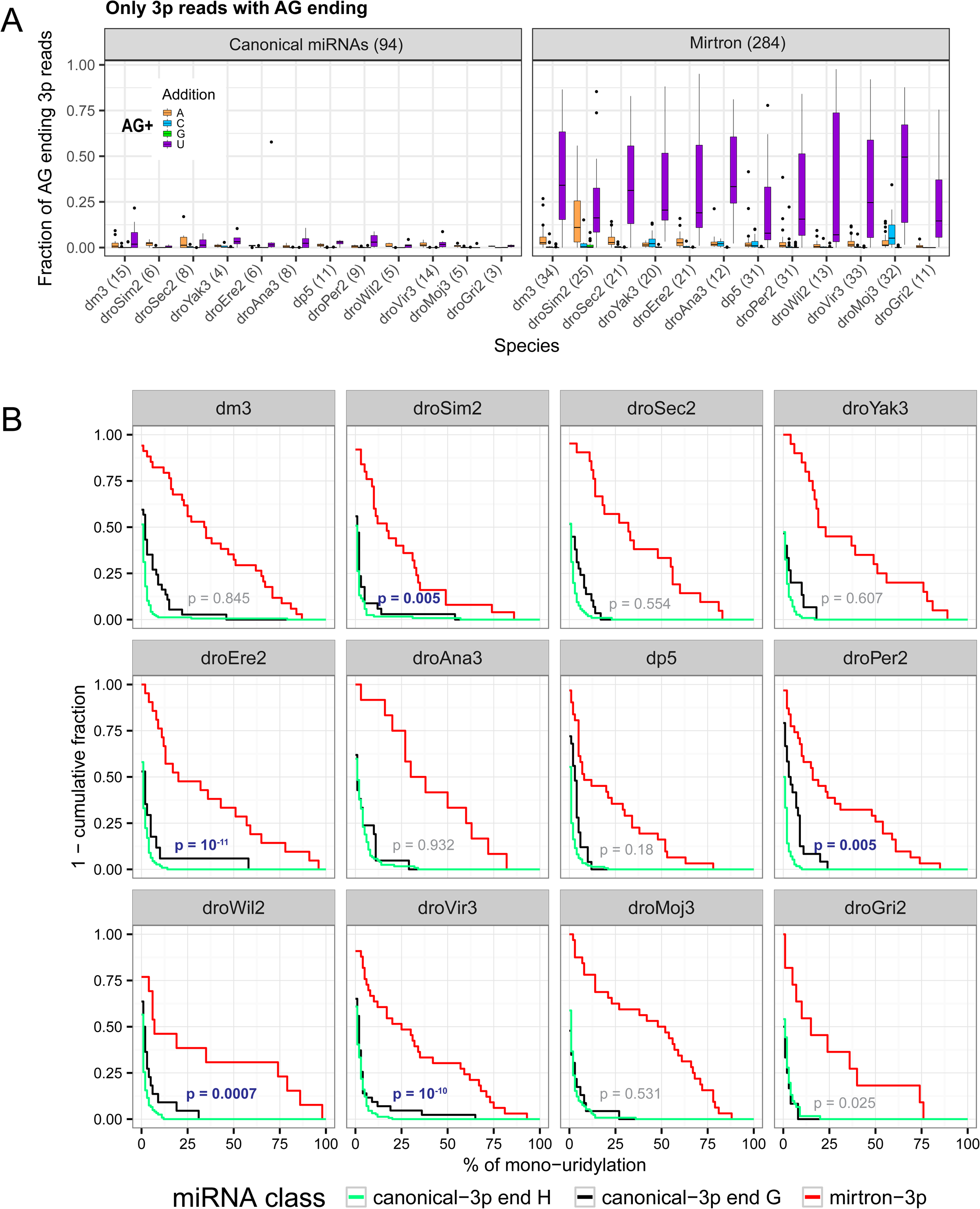
3’ end untemplated nucleotide additions for canonical miRNAs and mirtrons in 12 Drosophila species. (A) Proportion of AG ending 3’ arm miRNAs and mirtrons that contain reads within mono-A, C, G or U untemplated additions. Error bars represent the standard error of the mean. More mirtrons contain untemplated uridylation than comparable 3’ end AG-ending canonical miRNAs. (B) Species-specific empirical cumulative distribution function of mono-uridylation for mirtrons and canonical miRNAs with 3’ end ‘G’ nucleotide or non-‘G’ nucleotides (i.e. IUPAC ambiguity code ‘H’). P-value computed from two-tailed Wilcoxon Rank Sum Test between canonical 3’-end ‘H’ miRNAs and mirtrons. Significant differences in mono-uridylation distributions between these two classes are noted in blue text. P-values from comparisons between canonical 3’ end ‘H’ miRNAs and 3’ end ‘G’ miRNAs are all nonsignificant and not shown.

